# Marine sediments illuminate Chlamydiae diversity and evolution

**DOI:** 10.1101/577767

**Authors:** Jennah E. Dharamshi, Daniel Tamarit, Laura Eme, Courtney Stairs, Joran Martijn, Felix Homa, Steffen L. Jørgensen, Anja Spang, Thijs J. G. Ettema

**Author notes:** These authors contributed equally (Daniel Tamarit, Laura Eme). Correspondence to: Thijs J. G. Ettema.

## Abstract

The bacterial phylum Chlamydiae, which is so far comprised of obligate symbionts of eukaryotic hosts, are well-known as human and animal pathogens^1-3^. However, the Chlamydiae also include so-called environmental lineages^4-6^ that primarily infect microbial eukaryotes^7^. Studying environmental chlamydiae, whose genomes display extended metabolic capabilities compared to their pathogenic relatives^8-10^ has provided first insights into the evolution of the pathogenic and obligate intracellular lifestyle that is characteristic for this phylum. Here, we report an unprecedented relative abundance and diversity of novel lineages of the Chlamydiae phylum, representing previously undetected, yet potentially important, community members in deep marine sediments. We discovered that chlamydial lineages dominate the microbial communities in the Arctic Mid-Ocean Ridge^11^, which revealed the dominance of chlamydial lineages at anoxic depths, reaching relative abundances of up to 43% of the bacterial community, and a maximum diversity of 163 different species-level taxonomic unit. Using genome-resolved metagenomics, we reconstructed 24 draft chlamydial genomes, thereby dramatically expanding known interspecies genomic diversity in this phylum. Phylogenomic and comparative analyses revealed several deep-branching Chlamydiae clades, including a sister clade of the pathogenic Chlamydiaceae. Altogether, our study provides new insights into the diversity, evolution and environmental distribution of the Chlamydiae.

---

During a previous metagenomics study aimed at exploring microbial diversity of deep marine sediments from the Arctic Mid-Ocean Ridge^12^, we detected several Chlamydiae-related contiguous sequences (contigs). This finding prompted us to systematically screen marine sediment cores from a region surrounding Loki’s Castle hydrothermal vent field (Fig. 1a). We extracted DNA from 69 sediment samples (Supplementary Table 1) from various core depths, followed by screening using Chlamydiae-specific 16S rRNA gene primers. Chlamydiae were identified in 51 (74%) of the samples ranging in depth from 0.1 to 9.4 meters below seafloor (mbsf). We investigated the chlamydial relative abundance and diversity in 30 of these samples using bacterial-specific 16S rRNA gene amplicon sequencing, resulting in the identification of 252 operational taxonomic units (OTUs; clustered at 97% identity; Supplementary Data 1) that could be reliably assigned to Chlamydiae. This analysis revealed notable differences in chlamydiae relative abundance and diversity between samples (Supplementary Data 2), with individual samples showing relative abundances of up to 43% of the total bacterial community and up to 163 OTUs (Fig. 1b). Furthermore, we found that 155 of the 252 chlamydiae OTUs from our samples (hereafter referred to as “marine sediment chlamydiae”) could be identified in at least two samples. We further investigated the diversity of these marine sediment chlamydiae by performing a phylogenetic analysis in which the presently discovered chlamydial OTUs were placed amidst a recently compiled dataset of diverse chlamydiae 16S rRNA gene sequences^6^. This analysis revealed that the overall diversity of marine sediment chlamydiae spanned, and expanded, the known chlamydial diversity (Fig. 1c), revealing several deeply branching clades.

**Figure 1.**
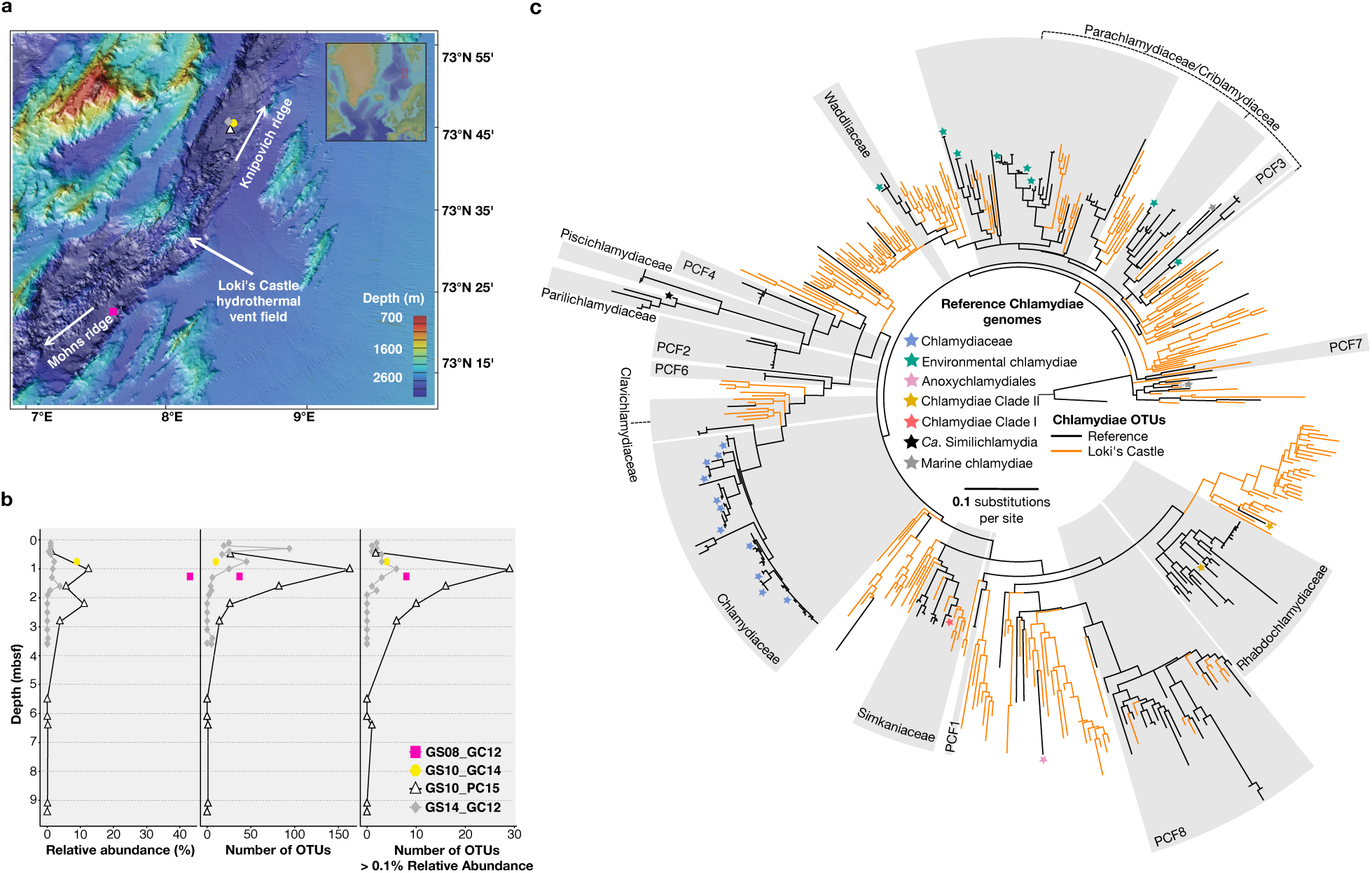
*Chlamydiae are diverse and abundant in Loki’s Castle marine sediments*. **a**, Bathymetric map of sediment core sampling locations taken northeast (GS10_GC14, GS10_PC15 and GS14_GC12) and southwest (GS08_GC12) of Loki’s Castle hydrothermal vent field. **b**, Chlamydial relative abundance, OTU number and abundant OTUs, as a factor of depth (meters below seafloor, mbsf) based on bacterial 16S rRNA gene amplicons. **c**, Maximum-likelihood (ML) phylogenetic tree (520 taxa, 476 sites) of marine sediment chlamydiae 16S rRNA gene sequences from amplicon OTUs (orange) and a reference (black) dataset, rooted using PVC taxa as an outgroup, inferred using IQ-TREE with the GTR+R7 model of evolution. Previously sequenced chlamydial genomes and metagenomic bins are labelled with stars.

To obtain genomic information from these marine sediment chlamydiae, we employed a genome-resolved metagenomics approach (Supplementary Fig. 1) in which we generated 249.6 gigabase pairs (Gbp) of paired-end reads from four sediment samples with a high relative abundance and diversity of chlamydial OTUs (Supplementary Fig. 2). Sequence assembly generated 5.85 Gbp of contigs larger than 1 kilobase pair (kbp). To assess the diversity of Chlamydiae-related sequences in these metagenome assemblies, we performed phylogenetic analyses of contigs containing 16S rRNA gene sequences or at least five genes of a conserved 15-ribosomal protein gene cluster. These analyses revealed numerous Chlamydiae-related contigs (Supplementary Fig. 2), most of which represented novel lineages that were distantly related to known chlamydiae. Contigs were binned into 24 highly complete (median 95% completeness, Supplementary Table 2) metagenome-assembled genomes (MAGs) on the basis of their tetranucleotide frequencies and patterns of sequence coverage across samples. They differed markedly in predicted genome size (1.33-1.99 Mbp), GC-content (26.4-48.9%) and gene content (Fig. 2, Supplementary Figs. 3 and 4).

**Figure 2.**
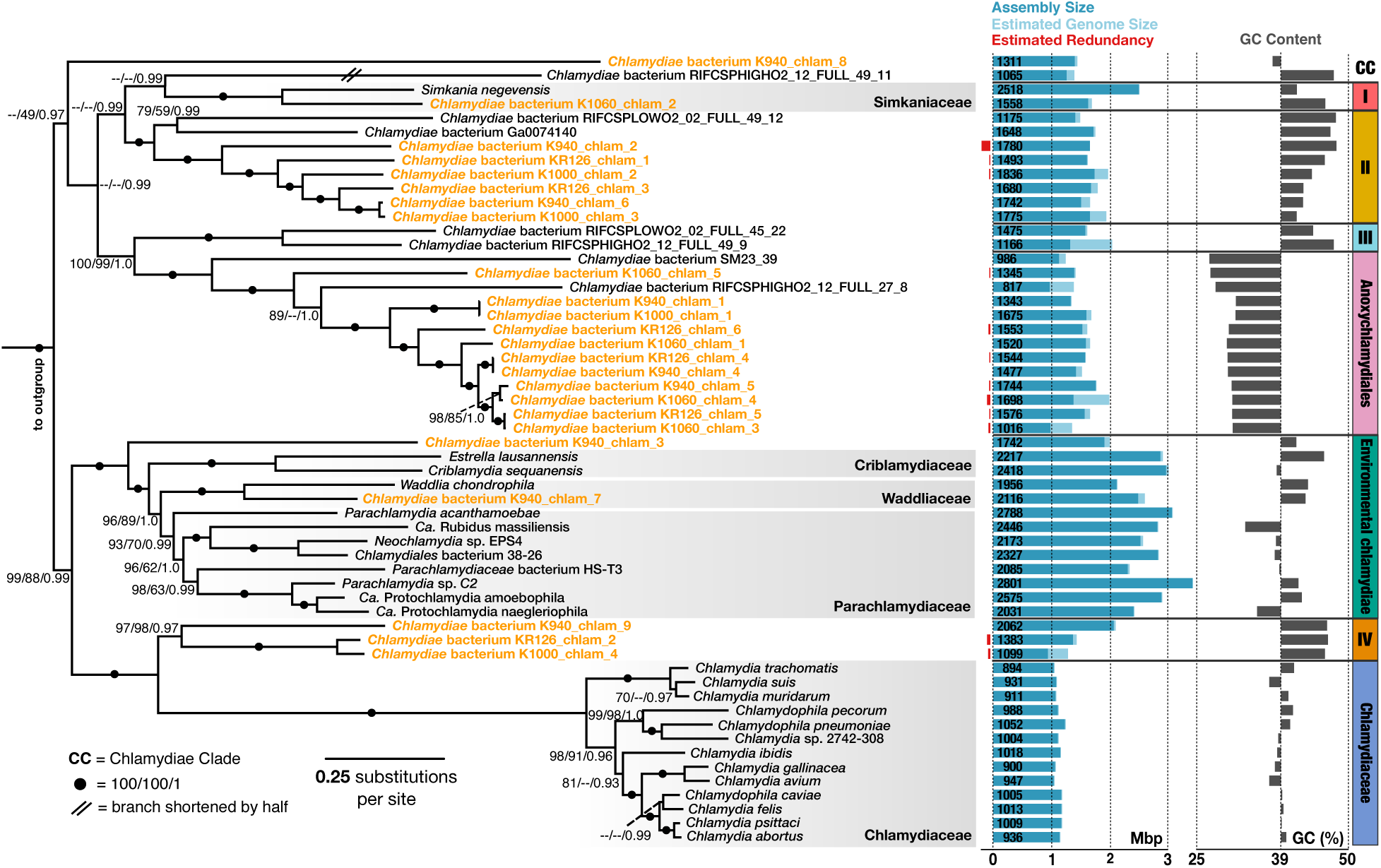
*Marine sediment lineages span Chlamydiae species phylogeny*. Bayesian phylogenetic tree of 38 concatenated single-copy conserved marker proteins in which marine sediment chlamydiae and previously characterized chlamydial representatives are shown in orange and black fonts, respectively. Previously established chlamydial families are shaded in grey. Branch support values were mapped onto the tree in the following order: non-parametric bootstrap support values (BV) for the full alignment (8006 sites) and reduced alignment (6005 sites after removal of the top 25% compositionally heterogeneous sites), each under the LG+C60+G+F derived PMSF approximation estimated by IQ-TREE, and posterior probability (PP) support values under the CAT+GTR+G4 model of evolution inferred with Phylobayes. For each species, the genome bin size, estimated genome size and redundancy are reported in Mbp along with the number of predicted open reading frames, and GC content.

To robustly infer the evolutionary relationships of the marine sediment chlamydiae to known lineages (Supplementary Table 3), we performed phylogenomic analyses of concatenated conserved marker protein sequence datasets. These analyses, which were designed to minimize potential long branch attraction and compositional bias artefacts, revealed that the newly reconstructed genomes form five new clades of high taxonomic rank (referred to as Chlamydiae Clades (CC) I-IV and Anoxychlamydiales; Fig. 2, Supplementary Fig. 3, Supplementary Discussion). While these phylogenomic analyses form clades that are in agreement with results based on previously available chlamydial genome data^13^, the branching order of these clades inferred differs considerably (Supplementary Discussion). Most marine sediment chlamydiae were placed into one of two well-supported, deeply branching clades, CC-II and Anoxychlamydiales, which also include four MAGs associated with estuary sediment, aquifer groundwater and a drinking water treatment plant^14-16^ (Fig. 2). Anoxychlamydiales are unique among Chlamydiae as they comprise members with gene repertoires indicative for an anaerobic lifestyle and will be treated in more detail in a complementary study (*manuscript in prep.*). Together, CC-I, CC-II, CC-III and Anoxychlamydiales form a superclade that is mainly comprised of uncultivated members represented by MAGs (Fig. 2, Supplementary Fig. 3). While the lifestyles of most members of this superclade remain elusive, the distinctive gene repertoires of the various clades point at functional differences (Supplementary Figs. 3 and 4). A second chlamydial superclade is comprised of the environmental chlamydiae, CC-IV and Chlamydiaceae (Fig. 2, Supplementary Fig. 3) and includes five of the marine sediment chlamydiae MAGs. While two of these MAGs affiliate with the environmental chlamydiae, which comprise symbionts of single-celled eukaryotes such as amoebozoa^7^, three MAGs resolve into a previously unknown clade referred to as CC-IV. Previous studies have suggested that Chlamydiaceae, a family that includes important animal pathogens such as the human pathogen *Chlamydia trachomatis*^17^, represents a deep-branching clade of the Chlamydiae phylum^13^. However, our phylogenetic analyses strongly support that Chlamydiaceae and CC-IV represent sisterclades, which together share a common ancestor with environmental chlamydiae (Fig. 2). Hence our results indicate that Chlamydiaceae represent a clade that was formed relatively late in chlamydiae evolution (Fig. 2), and that features specifically associated with pathogenicity in the Chlamydiaceae evolved much more recently than previously assumed.

**Figure 3.**
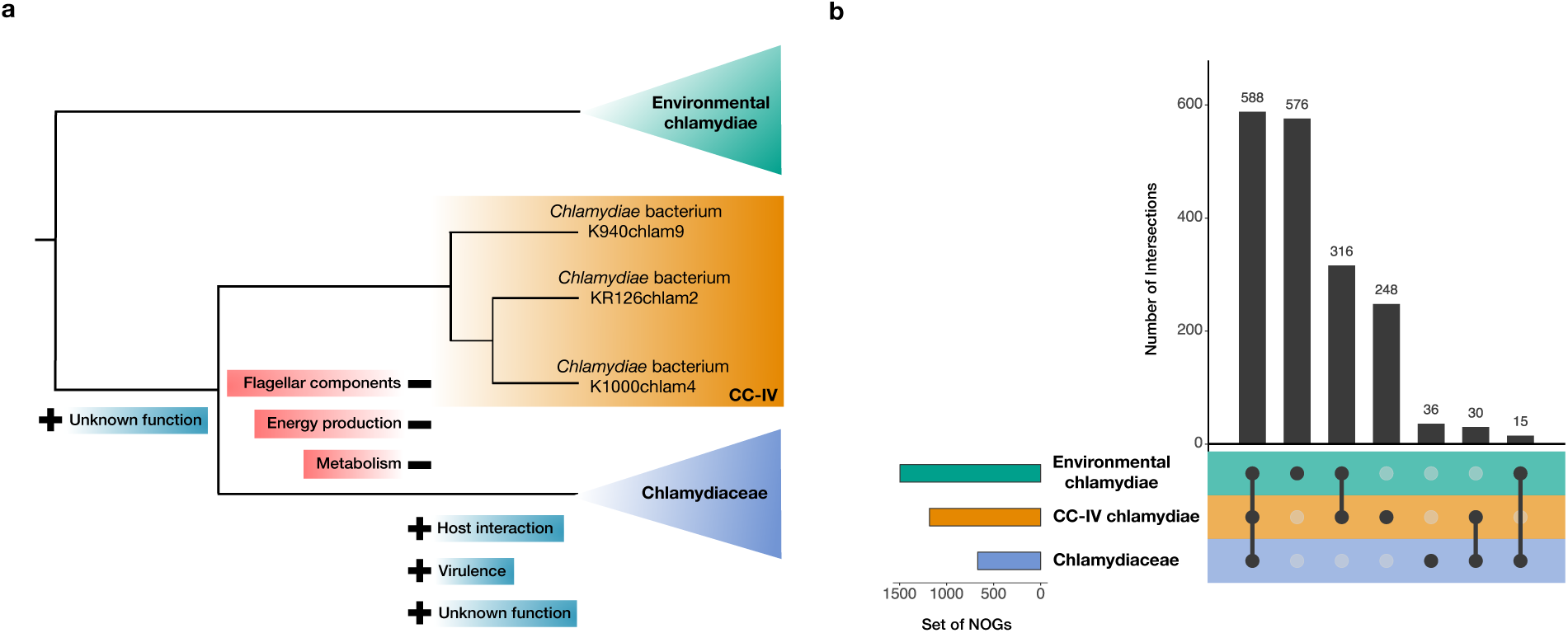
*Gene content evolution in Chlamydiaceae*. **a**, Schematic overview of lost and acquired cellular features, based on presence and absence patterns of NOGs (Supplementary Data 4), in pathogenic Chlamydiaceae, inferred from sister clade CC-IV. **b**, Plot showing intersections of NOGs conserved (in a third of lineages affiliated with each clade, Supplementary Table 2 and 3) across environmental chlamydiae, CC-IV and Chlamydiaceae.

Uncovering the sister relationship of CC-IV and the pathogenic Chlamydiaceae allowed us to re-evaluate the evolutionary events leading to their emergence (Fig. 3a). The genome sizes of CC-IV MAGs, while generally smaller than those of environmental chlamydia (Fig 2.), are larger than those of Chlamydiaceae (Fig. 2), suggesting that the latter have been subjected to genome reduction, a feature often observed in animal pathogens^18^. However, when considering only the gene set conserved across environmental chlamydiae, these differences are less drastic (Fig. 3b), and distribution patterns of Clusters of Orthologous Group (COG) categories found in CC-IV lineages more closely resemble that of environmental chlamydiae than Chlamydiaceae (Supplementary Fig. 5). Since their divergence from CC-IV, Chlamydiaceae have experienced reductive genome evolution (Fig. 3a-b, Supplementary Fig. 5), including the loss of several core components of central carbon metabolism, and *de novo* biosynthesis of nucleotides and amino acids (Supplementary Fig. 4, Supplementary Discussion). At the same time, they have acquired a small number of genes primarily linked to host-interaction and virulence, in addition to a notable set of conserved genes with unknown functions (Supplementary Fig. 6, Supplementary Discussion). It is likely that some of these unique features of the Chlamydiaceae are linked to the emergence of host-specificity. Further comparative analyses of CC-IV and Chlamydiaceae genomes revealed that only seven gene families are uniquely shared between these clades (Supplementary Fig. 7, Supplementary Discussion). This gene set is highly conserved across the Chlamydiaceae family, hinting at their putative importance in their adaptation to a pathogenic lifestyle. In addition, phylogenetic analyses of gene families with multiple homologs reveal gene duplication events have occurred both before and after the divergence of the Chlamydiaceae from CC-IV (Supplementary Fig. 7, Supplementary Discussion). However, the exact functions of these genes in Chlamydiaceae are currently unknown and our analyses suggest that they should represent a priority for future functional investigations. CC-IV genomes also encode homologs of several flagellar proteins (Supplementary Discussion), which clustered with flagellar components recently identified within distantly-related chlamydiae sampled from marine waters^19^ (Supplementary Fig. 8), indicating motility as a sharp difference between CC-IV and Chlamydiaceae.

All previously known members of the Chlamydiae, including environmental lineages, are obligate intracellular symbionts of eukaryotic hosts^20^ and display an obligate host-association for replication^10^. Similar to previously characterized lineages^20^, the herein identified marine sediment chlamydiae encode various homologs of proteins associated with this typical chlamydial lifestyle (Supplementary Fig. 4 Supplementary Discussion). For example, the marine sediment chlamydiae contain NF-T3SS (Supplementary Figs. 8 and 9) and NF-T3SS-specific effectors (Supplementary Data 3, Supplementary Discussion). NF-T3SS components are typically present in obligate bacterial symbionts that have a eukaryotic host, but they are also observed in free-living lineages (Supplementary Discussion). Similarly, most of these genomes contain genes encoding other secretion systems (Supplementary Fig. 10 and 11, Supplementary Discussion), some of which have been described to participate in adhesion and invasion in Chlamydiaceae. However, these systems have been shown to have alternative functions in other bacteria (Supplementary Discussion). Nucleotide transporters (NTTs), and in particular ATP/ADP transporters, are a characteristic feature of Chlamydiae and other bacteria typically associated with eukaryotic hosts^21^. All marine sediment chlamydiae genomes encode multiple NTT homologs (Supplementary Fig. 4 and 12, Supplementary Discussion), including possible ATP/ADP NTTs, which cluster together with other chlamydial sequences in phylogenetic analyses (Supplementary Fig. 12). Pathways for the *de novo* biosynthesis of nucleotides and amino acids are often incomplete in obligate intracellular symbionts^8,18^. Indeed, despite identifying chlamydial lineages with the genetic capacity to synthesize both purine and pyrimidine nucleotides *de novo*, we did not observe lineages capable of *de novo* synthesis of all nucleotides and amino acids (Supplementary Fig. 4 Supplementary Discussion).

Since the genomes of these marine sediment chlamydiae encode the typical features of host-dependency found in previously characterized chlamydiae^9^, we expected to find indications of potential eukaryotic hosts in the marine sediment samples. While attempts to amplify 18S rRNA genes from DNA isolated from these sediment samples were unsuccessful, screening the four marine sediment metagenome datasets resulted in the identification of a few 18S rRNA gene fragments (Supplementary Table 4, Supplementary Discussion). However, no such sequences could be identified in the metagenome of the marine sediment sample with the highest Chlamydiae relative abundance (Fig. 1b). Given the absence of obvious host candidates, we explored the possibility that these genome sequences derive from persistent marine sediment chlamydiae cells, such as from elementary bodies, which can survive outside of host cells and even remain metabolically active, despite being unable to replicate^22,23^. Estimation of replication rates based on sequence read-coverage^24^ indicated that these MAGs are derived from actively dividing marine sediment chlamydiae (Fig. 4b). Thus, our results suggest that at least some of these marine sediment chlamydiae are not associated with eukaryotic hosts.

**Figure 4.**
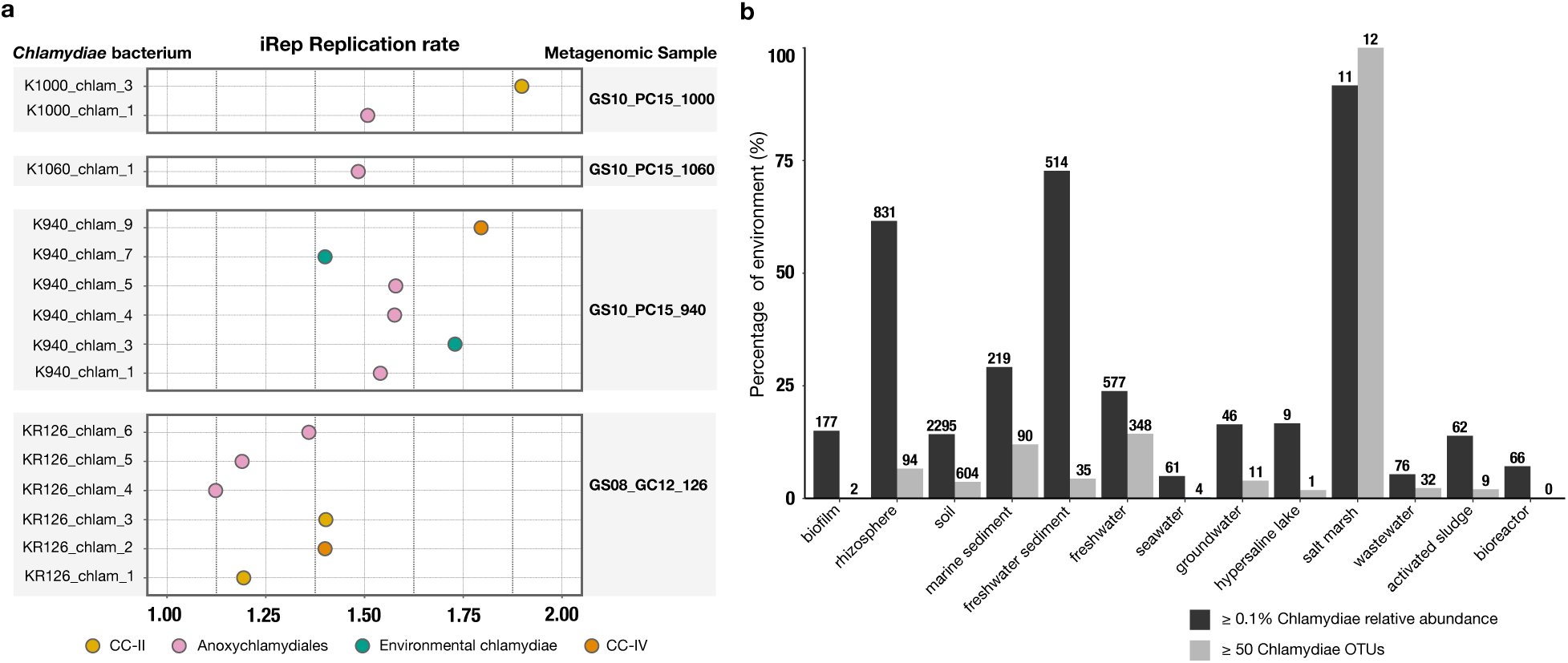
*Estimated iRep replication rate of marine sediment chlamydiae, and bar plot showing presence of chlamydiae in diverse environments*. **a**, Replication rates of marine sediment chlamydiae inferred using iRep. Indicates the proportion of the microbial population represented by the metagenome-assembled genome that is actively dividing. **b**, Percentage of samples from selected environments which contain a chlamydial relative abundance of ≥ 0.1%, or ≥ 50 OTUs based on publicly available 16S rRNA gene amplicon datasets. Values above each bar represent the absolute number of samples.

Finally, we investigated chlamydial diversity in environments other than marine sediments. An analysis of publicly available 16S rRNA gene amplicon datasets from a variety of environments revealed that fresh water, ground water, salt marshes and wastewater often harbour diverse and abundant chlamydiae (Fig. 4b). These findings likely represent considerable underestimations of chlamydial diversity and relative abundance since mismatches in commonly used 16S rRNA primer sets (e.g., Earth Microbiome Project^25^, Supplementary Table 5) generally do not capture known Chlamydiae diversity (Supplementary Discussion).

Our work reports the existence and genomic characterization of an extended diversity of Chlamydiae-related lineages in deep marine sediments and provides insights into the evolution and diversification of the Chlamydiae phylum. Using sophisticated phylogenomic methods, we used a robust phylogenomic framework for investigating Chlamydiae evolution. Furthermore, we identified several new Chlamydiae clades of high taxonomic rank, including a sister clade of the pathogenic Chlamydiaceae, which provided insights into the early evolution of this family. Altogether, our findings indicate that Chlamydial diversity and abundance has been underappreciated in environmental surveys, and our observations represent a shift in our view of the environmental distribution of Chlamydiae. These results indicate differences in basic ecology and lifestyle across this phylum, and contribute to a comprehensive understanding of its evolution, including the emergence of host dependency and pathogenicity.

## Acknowledgements

We thank M. Horn, L. Guy, S. Abby, L. Juzokaite, A. E. Lind, K. Zaremba-Niedzwiedzka and E. C. Fernandez for technical assistance and/or for useful advice and discussions. We also acknowledge the help from chief scientist R. B. Pedersen, the scientific party and the entire crew on board the Norwegian research vessel G.O. Sars during the summer 2008, 2010 and 2014 expeditions. All sequencing was performed by the National Genomics Infrastructure sequencing platforms at the Science for Life Laboratory at Uppsala University, a national infrastructure supported by the Swedish Research Council (VR-RFI) and the Knut and Alice Wallenberg Foundation. We thank the Uppsala Multidisciplinary Center for Advanced Computational Science (UPPMAX) at Uppsala University and the Swedish National Infrastructure for Computing (SNIC) at the PDC Center for High-Performance Computing for providing computational resources. This work was supported by grants of the European Research Council (ERC Starting grant 310039-PUZZLE_CELL), the Swedish Foundation for Strategic Research (SSF-FFL5) and the Swedish Research Council (VR grant 2015-04959) to T.J.G.E. C.W.S. is supported by a European Molecular Biology Organization long-term fellowship (ALTF-997-2015) and the Natural Sciences and Engineering Research Council of Canada postdoctoral research fellowship (PDF-487174-2016). Funding was received from the European Union’s Horizon 2020 research and innovation program under the respective Marie Sklodowska-Curie grant agreements 625521 (to A.S.) and 704263 (to L.E.). A.S. is supported by the Swedish Research Council (VR starting grant 2016-03559) and the NWO-I foundation of the Netherlands Organisation for Scientific Research (WISE fellowship).

## Author Contributions

T.J.G.E. and A.S. conceived the study, and J.E.D. and T.J.G.E. performed the experimental design. S.L.J. provided environmental samples. S.L.J. and J.E.D. performed DNA extractions and performed PCR-based screening. J.E.D. generated 16S rRNA gene amplicons. F.H. and J.E.D. performed metagenomic sequence assemblies. J.E.D., J.M. and F.H. performed genome-resolved metagenomics analyses. J.E.D., F.H. and T.J.G.E. analyzed 16S rRNA gene amplicon sequence data. J.E.D., C.S., D.T., L.E., J.M., and A.S. analyzed genomic data and performed phylogenetic analyses. J.E.D., C.S., L.E., D.T., A.S. and T.J.G.E. interpreted the obtained data and results. T.J.G.E. and J.E.D wrote, and all authors edited and approved, the manuscript.

## Methods

### Sample acquisition

Sediment cores were retrieved from the Arctic Mid-Ocean Ridge near Loki’s Castle^11^ hydrothermal vent field over multiple sampling expeditions: GS08_GC12 in 2008 (3.3 m)^26^, GS10_GC14 (2 m) and GS10_PC15 (11.2 m) in 2010^27^, and GS14_GC12 (3.6 m) in 2014. Sediment samples at various depths (Supplementary Table 1) were collected on-board and immediately frozen for prospective microbiological analysis. Geochemistry and sediment characteristics of these samples have been published previously^26,27^, except in the case of GS14_GC12.

### Sample screening and bacterial 16S rRNA gene amplicon sequencing

DNA was extracted using the PowerLyzer® PowerSoil® DNA kit in conjunction with the PowerLyser® 24 homogenizer as per manufacturer’s instructions (MOBIO) with minor modifications using 0.5-0.7 g of sediment and the addition of polyadenosine to increase DNA yield^28^. Taxonomic coverage of all primer pairs, for both sample screening and amplicon sequencing, was tested *in silico* using SILVA TestPrime^29^ with the SSU r132 RefNR database (Supplementary Table 5). All primers used have been published previously^29,30^ except for Chla-310-a-20 which was designed with PRIMROSE^31^ (Supplementary Table 5). Reaction conditions for each primer pair are found in Supplementary Table 5. Primer pair Chla-310-a-20 and S-*-Univ-1100-a-A-15^29^, which amplifies an approximately 800 bp region of the 16S rRNA gene, was used to screen 69 Loki’s Castle marine sediment samples (Supplementary Table 1) for Chlamydiae. Primer pair 574*f^30^ and 1132^30^ were used to screen for 18S rRNA genes in these same sediment samples. The latter primer pair did not yield positive amplification products, in spite of its broad taxonomic coverage of known eukaryotic 18S rRNA genes (Supplementary Table 5).

Thirty sediment samples with positive PCR screening results for Chlamydiae were selected for further investigation. Bacterial-specific 16S rRNA gene amplicons were sequenced using a two-step PCR approach. Primer sequences and reaction conditions are reported in Supplementary Table 5. In the first step, a ca. 500 bp region of the 16S rRNA gene was amplified in triplicate, to account for random PCR drift^32^, using bacterial primers (S-D-0564-a-S-15 and S-D-Bact-1061-a-A-17^29^). For each reaction, replicates were pooled and purified using magnetic AMPure XP beads (Agencourt). In the second PCR step, libraries were constructed using adaptor sequences and reaction conditions from the TruSeq DNA LT Sample Prep Kit (Illumina), before sequencing with Illumina MiSeq (2×300 bp). Using cutadapt^33^ v. 1.10 sequence reads shorter than 100 bases were filtered out, 3’ ends trimmed to a minimum Phred quality score of 10, and primer sequences removed. Forward and reverse reads were merged using VSEARCH^34^ v. 1.11.1 (--fastq-minovlen 16), de-replicated (--derep_fulllength), and clustered into centroid OTUs (threshold = 97%). Chimeras were detected and removed using UCHIME^35^ with the SILVA123.1_SSUref_tax:99 database^36^. Taxonomy was assigned using the LCAClassifier^37^ from CREST-2.0.5 with silvamod106 as the reference database.

### Metagenome sequencing and assembly

The Fast DNA Max Spin kit and Fast DNA Spin kit (for GS10_PC15_940 only, 4 replicates pooled) were used according to manufacturer’s protocols (MP Biomedical), with the addition of polyadenosine, to extract DNA from sediment samples chosen for further analysis (GS08_GC12_126, GS10_PC15_940, GS10_PC15_1000, and GS10_PC15_1060). Libraries were prepared using the Nextera DNA Library Prep kit (Illumina) with 25 ng of input DNA, and sequenced with Illumina HiSeq in rapid-mode (2x 250 bp). For GS10_PC15_1060, reads from three separate HiSeq runs and undetermined reads (reads with barcode mismatches) from one run were combined. Quality control to remove low-quality reads was performed using Trimmomatic^38^ 0.35 with the options: SLIDINGWINDOW:4:12, MINLEN:50, ILLUMINACLIP:TruSeq Illumina Universal Adaptors. FastQC^39^ v.011.4 was used to visually evaluate sequence quality before and after processing. Using the fq2fa program from the IDBA-UD^46^ package, paired reads were interlaced and Ns removed, using the options ‘merge’ and ‘filter’ (except for GS10_PC15_1060 reads, where Ns were retained, as it improved the assembly). Trimmed reads from each sample were subjected to iterative *de novo* assembly using IDBA-UD 1.0.9 (minimum k-mer size = 20 and maximum k-mer size = 124, except for GS08_GC12_126 for which maximum k-mer size = 100). Assembly quality and statistics (Supplementary Fig. 2) were assessed using QUAST^40^ v3. Open reading frames (ORFs) across assembled contigs in each metagenome were called with Prodigal^41^ v.2.6.3.

### Assesment of metagenome microbial composition

To investigate the microbial composition of the samples, ‘ribocontigs’, i.e. a contig encoding at least 5 of 15 ribosomal proteins found in an operon conserved across prokaryotes^42^, were identified in the metagenomic assemblies using the RP15 pipeline^43^. As part of this, a maximum-likelihood (ML) phylogeny using the LG+C60+G model of evolution (Supplementary Fig. 2) was inferred from a dataset consisting of concatenated ribosomal proteins extracted from metagenomic ribocontigs and 90 phylogenetically diverse reference taxa including bacteria and archaea^44^ in addition to PVC representative species (Supplementary Tables 3 and 6).

### Obtaining Chlamydiae metagenome assembled genomes

A differential coverage binning approach was used to obtain metagenome-assembled genomes (MAGs). For each metagenome assembly, contig coverage was estimated using pseudoalignment with Kallisto^45^ 0.42.5, with sequence reads from each of the four samples. Differential coverage profiles were generated for each assembly using a freely available script (github.com/EnvGen/toolbox/tree/master/scripts/kallisto_concoct/input_table.py), provided by Johannes Alneberg. To give more statistical weight to longer contigs and to reduce the impact of chimeric sequences^46^, contigs larger than 20 kb were split into 10 kb fragments. CONCOCT^46^v.0.40 was used to cluster contigs within each focal assembly into MAGs, using their differential coverage profile^47^, tetranucleotide frequency, and different contig length cut-offs (1 kb, 2 kb and 3 kb). Due to the large diversity in sample GS10_PC15_1060, the maximum number of bins was adjusted (1500 bins for 1 kb, and 1000 bins for 2 and 3 kb length cut-offs). Putative chlamydial metagenomic bins were identified by phylogenomic analyses of concatenated ribosomal proteins encoded on ribocontigs (Supplementary Fig. 2). Completeness and redundancy was assessed using the micomplete^48^ pipeline (without weighting), provided by Lionel Guy (https://bitbucket.org/evolegiolab/micomplete) using a custom marker set (Supplementary Table 7) corresponding to genes present in complete Chlamydiae genomes (Supplementary Table 3). For each ribocontig, the corresponding metagenomic bin with the highest completeness and lowest redundancy across the CONCOCT iterations was selected for further analysis. Chlamydial metagenome bins were subjected to manual cleaning using mmgenome^49^, resulting in medium and high quality MAGs (Supplementary Table 2) based on MIMAG standards^56^. Differential coverage across samples, GC content, linkage, the presence of chlamydial-specific marker proteins^20^, and single-copy bacterial marker proteins (Supplementary Table 7) were visualized using mmgenome^50^. Contigs with profiles diverging from the majority of contigs were removed. Linkage information, i.e. information about which contigs are connected by read pairs, was calculated with read mapping using Bowtie2^51^, followed by application of the script bam_to_linkage.py from CONCOCT^46^. After generating the final MAGs, contigs that had been split into 10 kb contig fragments were joined again if more than half of the fragments of a specific contig were assigned to a specific MAG. Otherwise, all fragments of the contig in question were discarded.

### Selection of published PVC genomes for comparative and phylogenetic analysis

Selection of PVC superphylum representatives was facilitated by a phylogenetic analysis of ribocontigs from PVC member genomes available in NCBI (as of February 6^th^, 2017). Using the RP15 pipeline^43^, a maximum-likelihood (ML) phylogeny of these ribocontigs was inferred, using RAxML^52^ 8.2.4 under the PROTCATLG model of evolution. Branch support was estimated through 100 rapid bootstrap replicates (‘-f a’) (Supplementary Fig. 13). Phylogenetically diverse representatives from non-Chlamydiae PVC phyla were selected (Supplementary Table 6) to be used as an outgroup for phylogenomic analyses. With the exception of Chlamydiaceae (for which we used genomes classified as reference or representative in NCBI), all Chlamydiae genomes (Supplementary Table 3) were used in protein phylogenies and comparative genomics analyses. For determining interspecies relationships within the Chlamydiae phylum (Supplementary Fig. 13), only one representative was kept whenever several chlamydial genomes had a near-identical phylogenetic placement (Supplementary Table 3). Chlamydial species representative MAGs and single-cell assembled genomes (SAGs), which were made available on NCBI subsequently (between February 6^th^, 2017 and April 18^th^, 2018, Supplementary Table 3), were included in comparative genomics and phylogenetic analyses (Supplementary Fig. 4).

### Protein clustering and gene annotation

Gene features of marine sediment chlamydiae MAGs (Supplementary Table 2) were annotated with Prokka^53^ v1.12, using a version that allows for partial gene prediction (GitHub pull request #219). All protein sequences from both chlamydiae in NCBI (Supplementary Table 3) and from those in this study (Supplementary Table 2) were searched against databases as follows: top hits with and excluding Chlamydiae against the *nr* database and taxonomic classification (Lowest Common Ancestor (LCA) algorithm, ‘-f 102’) using blastp (--more-sensitive) from DIAMOND^54^ aligner v0.9.19.120; PFAM (PF)^55^ and Interpro (IPR)^56^ domains, and MetaCyc^57^ and KEGG^58,59^ pathway annotations, were assigned using Interproscan^60^ version 5.22-61.0; KEGG ‘KO’ numbers were assigned using GhostKOALA^61^. Protein sequences were also mapped to the eggNOG orthologous groups^62^ version 4.5 using eggNOG-mapper^63^, at both the universal ‘-d NOG’ and bacterial level ‘-d BACT’. The presence of proteins of interest across Chlamydiae genomes were assessed using these annotations and database searches (Supplementary Data 3). The presence of amino acid and nucleotide *de novo* biosynthesis and central carbon metabolism pathways (Supplementary Data 3) were manually investigated using KEGG^58^ KO number assignments.

### Detection of flagellar genes, secretion systems, NF-T3SS secreted proteins, eukaryotic-like domains and subcellular targeting signals in the host

Genes related to secretion systems and the flagellum were detected for most chlamydiae using MacSyFinder^71^ with the protein models built by Abby et al.^64^, using the mode ‘gembase’ for complete genomes and ‘unordered’ for incomplete genomes. We predicted NF-T3SS secreted proteins, eukaryotic-like domains (ELD), and putative subcellular targeting signals to eukaryotic cellular compartments for all predicted proteins from chlamydiae (Supplementary Table 2 and 3) and from nine well-characterized PVC representatives with free-living lifestyles. For this, we used EffectiveDB^65^ in ‘genome mode’, i.e. enabling the prediction of secretion systems and the discovery of novel ELD (Supplementary Data 3).

### Phylogenetic analyses

Unless otherwise stated, gene or protein sequences were aligned using MAFFT-L-INS-i^74^ v7.271 and trimmed with trimAl^66^ v1.4 (--gappyout). Identical sequences were removed, and alignments were inspected manually. ML phylogenetic analyses were performed using IQ-TREE^67^ 1.5.3 with automated model selection^68^ among the following models: the empirical LG model^69^ the empirical profile mixture models (C10 to C60) combined with the LG exchangeability matrix (e.g., LG+C10)^70^, with or without empirically determined amino acid frequencies (+F), and free or gamma-distributed rates (+R or +G)^71^. Bootstrap support values were inferred from 1000 ultrafast bootstrap (ufBV) replicates and from 1000 replicates of the SH-like approximate likelihood ratio test (SH-aLRT). All unprocessed phylogenetic trees can be found in Supplementary Data 4.

### Phylogenetic inference of interspecies relationships within the Chlamydiae phylum

#### Identification and phylogenetic analysis of 16S rRNA gene amplicon and metagenome sequences

Barrnap^53^ 0.8 was used to identify 16S/18S rRNA genes in the metagenomic assemblies, three iterations with ‘kingdom’ set to ‘euk’, ‘arc’ and ‘bac’ were run, with ‘reject’ set to 20% of the rRNA gene. Sequences were taxonomically classified using LCAClassifier^37^ from CREST-3.0.5 with silvamod128 as the reference database. The microbial composition of each metagenome can be found in Supplementary Table 4. A reference dataset of near-full length chlamydiae 16S rRNA gene sequences from various metagenomic and amplicon sequence databases^6^ was used for determining the phylogenetic placement of these sequences. ML phylogenetic inference (Fig. 1c, Supplementary Fig. 2) was performed with the GTR+R7 model of evolution^72^ (based on model selection).

#### Selection of single-copy marker proteins

We selected marker proteins using NOGs that were present in a single-copy in 95% of near-complete PVC genomes (Supplementary Fig. 14). For each of the 149 markers protein initially identified, alignments were generated and manually curated to remove divergent sequences prior to final alignment and trimming with trimAl^66^ v1.4 (--automated1). ML phylogenies were inferred for each alignment, and proteins displaying patterns of vertical inheritance were selected. Due to strain microdiversity, it is not uncommon to have contigs from two different, but closely related, lineages together in a MAG, resulting in redundancy. For each marker, in cases of multiple copies from the same MAG, alignments and corresponding single-gene phylogenies were manually inspected to determine if it represented a paralog, a redundant gene copy, or partial sequences from the same gene. If redundant sequences overlapped with non-identical regions, all sequences from the same genome were removed; if they were placed at the end of a contig and shared an identical overlapping region (longer than 30 nucleotides), the sequences were merged; and if they were partial non-overlapping protein fragments, the longer fragment was selected and the shorter one removed. This inspection resulted in 126 remaining protein markers which were subjected to discordance filtering (or “χ^2^ trimming”) to remove markers with the most conflicting phylogenetic signal^73^. The 20% of markers whose taxon bipartition profiles were least concordant with the others, resulting in a high discordance score (Supplementary Fig. 15), were removed. This final set of 98 marker proteins was concatenated and used for species-tree reconstruction (Supplementary Table 8).

#### Phylogenetic inference of interspecies relationships using concatenated marker proteins

Chlamydial species representatives (released prior to February 6^th^, 2017, Supplementary Table 3) and marine sediment chlamydiae MAGs (Supplementary Table 2) were used to investigate interspecies phylogenomic relations within the Chlamydiae phylum, using other PVC representatives as an outgroup (Supplementary Table 6). Marker proteins were separately aligned and trimmed before being concatenated into a supermatrix. ML inference was performed on a supermatrix of all 98 identified single-copy marker proteins (28,286 amino acid positions), and a sub-selection of the 55 (14,212 amino acid positions) and 38 (7,894 amino acid positions) markers with the highest representation among lineages (Supplementary Table 8), using the LG+C60+ G+F model of evolution. The tree topologies inferred were similar across all three datasets (Supplementary Fig. 16), indicating that the 38 marker protein sub-selection was sufficient for further inferences. Several more in-depth phylogenetic analyses were applied to the supermatrix of 38 marker proteins (Fig. 2), using a smaller outgroup to allow for more computationally intensive analyses (Supplementary Table 6). PMSF is a site-heterogeneous mixture model that can closely approximate complex mixture models such as LG+C60+G+F while reducing computational time several-fold^74^, making full bootstrapping practical. A ML phylogeny was inferred using the PMSF model implemented in IQ-TREE^67^ 1.5.5, with a guide-tree inferred using the LG+C60+G+F model of evolution, with 100 nonparametric bootstrap replicates (BV). The same analysis was performed on the alignment, after it was subjected to χ^2^-trimming. Here, the proportion of most heterogeneous sites were was removed in a step-wise fashion from 5% to 95% of sites, as previously described^48,75^. The resulting χ^2^ test statistics for each lineage under the various heterogenous site removal treatments were visualized for the chlamydiae and PVC outgroup (Supplementary Fig. 17) and 25% removal was chosen as the treatment which best lowered compositional heterogeneity while retaining the largest number of informative sites. Bayesian analysis of this alignment was performed under the CAT+GTR+Γ4 model with PhyloBayes MPI^76^ 1.7a. Four independent Markov chain Monte Carlo chains were run for ∼55,000 generations, after which, three chains converged (maxdiff = 0.12; burn-in = 15,000). Subsequently, these 38 markers were updated with additional Chlamydiae species representatives (released between February 6^th^, 2017 and April 18^th^, 2018, Supplementary Table 3). Single-protein phylogenies and alignments including these lineages were inferred and manually inspected, as described above, followed by concatenation of each of the 38 trimmed alignments. A robust ML phylogeny was inferred based on the resulting supermatrix using the selected LG+C60+PMSF model, as described above (Supplementary Fig. 3).

### Phylogenetic analyses of proteins of interest

#### Flagellar genes and NF-T3SS

Sequences identified for each NF-T3SS/flagellar gene (see above) were gathered, separately aligned and trimmed with BMGE^77^ v. 1.12 (-m BLOSUM30). ML phylogenies were inferred with IQ-TREE^67^ v1.6.5 with model selection among the following models: the empirical LG model^69^ the empirical profile mixture models (C10 to C60) combined with the LG exchangeability matrix (e.g., LG+C10)^70^, with or without empirically determined amino acid frequencies (+F), and specific free rates (+R0, +R2, +R4 or +R6)^71^. An additional ML reconstruction was run with the same alignments using the PMSF^74^ approximation of the selected model and the previously obtained tree as a guide tree, with 100 BV. These reconstructions were used to reclassify sequences between the NF-T3SS/flagellar homologues and outgroups. Phylogenies of individual components were largely congruent (Supplementary Data 4), allowing us to reconstruct a concatenated phylogeny using the separately trimmed alignments of the *sctJ, sctN, sctR, sctS, sctT, sctU*, and *sctV* homologues with IQ-TREE^67^ v1.6.5 as above (Supplementary Data 4).

#### Nucleotide transporters

The diversity and phylogenetic placement of nucleotide transporter (NTT) proteins from marine sediment chlamydiae was investigated. The region corresponding to the NTT PF^78^ domain (PF03219) with a single-domain structure was predicted using hmmscan from the HMMER v3.1 toolkit (http://hmmer.org) and extracted from the full protein sequences. An initial phylogeny (not shown) was reconstructed based on this conserved region using FastTree2^79^, and indicated two monophyletic clades separated by a long branch as identified previously^80^. Further phylogenetic analyses were performed separately on the clade containing “canonical NTT” and the “other NTT” clade (Supplementary Fig. 12). NTT proteins containing a HEAT domain were also investigated (Supplementary Fig. 12). Proteins homologous to the *Chlamydia trachomatis* query were retrieved from marine sediment chlamydiae using BLASTP and were combined with NTT-HEAT sequences identified in a prior study^80^. All three NTT datasets were each aligned before being trimmed with TrimAl^81^ v1.4, using the ‘gappyout’ option for the “canonical NTT” alignment and the ‘automated1’ option for the “sister NTT” and NTT-HEAT alignments. The LG+F+R8 model of evolution was selected for ML phylogenetic inference, except for the NTT-HEAT alignments, where the LG+F+R6 model was selected instead. Chlamydiae NTTs that have been functionally characterized were annotated in the resulting phylogenies (Supplementary Fig. 12, Supplementary Discussion).

#### Proteins conserved in CC-IV and Chlamydiaceae

NOGs^62^ and PF^78^ domains identified uniquely in clade CC-IV and Chlamydiaceae among chlamydial lineages (Supplementary Table 3) were compiled (Supplementary Fig. 6 and 7) and phylogenetic investigation of these sequences with the PF domains PF04518 and PF05302 performed (Supplementary Fig. 7). Hmmalign, from the HMMER v3.1 toolkit (http://hmmer.org), was used to extract the region corresponding to the domains from sequences, which were aligned and trimmed as outlined previously. ML phylogenies were inferred using the PMSF model implemented in IQ-TREE^67^ 1.5.5, with a guide-tree inferred using the LG+C20+G+F model of evolution, with 100 BV. An hmmsearch against the *nr* database confirmed that these PF domains are only present in CC-IV and Chlamydiaceae chlamydiae.

### Determination of replication rates

We used iRep^24^ to determine the replication rate of the microbial population represented by each MAG. The tool uses differences in sequencing coverage that arise bi-directionally across the genomes of replicating bacteria, due to the single origin of replication, to infer a population-level rate of replication^24^. Since iRep calculations require MAGs that are estimated to be 75% complete with 5X coverage, we analyzed only 15 of the 24 MAGs that met these criteria. Sequenced reads from each metagenome were mapped to corresponding assembled contigs using Bowtie2^51^ with the ‘reorder’ option, before applying iRep^24^ v.1.10 with default settings (Fig. 4a).

### Chlamydial environmental diversity

Using the Integrated Microbial NGS platform (IMNGS)^82^ (accessed March 5^th^, 2018), which systematically screens prokaryotic 16S rRNA gene amplicon datasets deposited as sequence read archives in NCBI, the percentage of samples from select environments with a relative abundance of over 0.1% Chlamydiae, and a percentage with at least 50 OTUs was assessed (Fig. 4b).

### Data visualization

Plots in figures were made with R v.3.2.2 (R Development Core Team, 2008) using the packages ggplot2^83^ and gplots^84^. NOG absence and presence profiles across chlamydial genomes were evaluated using top NOG hits identified by eggNOG mapper^63^ as described above (Supplementary Data 3). A binary distance matrix of NOG presence patterns was hierarchically clustered using hclust with the ‘average’ agglomeration method in R, and a heatmap generated using heatmap.2 from the gplots^84^ package (Supplementary Fig. 3). Intersection plots were implemented using the R package UpSetR^85^ (Fig. 3b). Synteny figures for NF-T3SS/flagellar systems (Supplementary Figs. 8, 9, 10 and 11) were generated using the R package GenoPlotR^86^. Phylogenetic trees were visualized and edited using Figtree^87^ v1.4.2 and iTOL^88^. Protein domains were visualized and mapped to phylogenetic trees using iTOL^88^. Bathymetry was uploaded and visualized using GeoMapApp V. 3.6.6 (http://www.geomapapp.org). Figures were made and edited using Inkscape and Adobe Illustrator.

### Data availability

Raw sequence reads for both 16S rRNA gene amplicons and metagenomes have been deposited to the NCBI Sequence Read Archive repository under BioProject PRJNA504765. Whole Genome Shotgun projects for metagenome assemblies GS08_GC12_126, GS10_PC15_940, GS10_PC15_1000 and GS10_PC15_1060 have been deposited at DDBJ/ENA/GenBank under the accessions SDBU00000000, SDBV00000000, SDBS00000000 and SDBT00000000, respectively. The versions described in this paper are versions SDBU01000000, SDBV01000000, SDBS01000000 and SDBT01000000. Accessions for MAGs generated in this study can be found in Supplementary Table 2, and are linked to BioProject PRJNA504765. Files containing sequence datasets, alignments, all 16S rRNA gene amplicon OTUs and additional data generated in this study are archived at the Dryad Digital Repository: https://datadryad.org/resource/XXX.

