## Supplementary Information for "Marine sediments illuminate Chlamydiae diversity and evolution"

### Supplementary Discussions

#### 1. Evolutionary relationships within the Chlamydiae phylum

We performed several in-depth phylogenomic analyses to reconstruct interspecies relationships within the Chlamydiae phylum. To build upon previous work[1-3], we have increased taxon sampling and put a particular emphasis on applying state-of-the-art approaches aiming to detect and alleviate potential phylogenetic artifacts that can be caused by long-branching taxa and sequence composition heterogeneity (see Methods).

Our phylogenomic analyses in maximum likelihood and Bayesian frameworks allowed us to resolve seven well-supported Chlamydiae Clades (CC) of putatively high taxonomic rank. These include five newly identified clades, CC-I through CC-IV and Anoxychlamydiales, which are primarily composed of uncultured chlamydial lineages represented by metagenome-assembled genomes (MAGs). The phylogenetic placement of most lineages, and deep-branching relationships between chlamydia clades were well-resolved and consistent across phylogenomic reconstructions (Fig. 2, Supplementary Figs. 3 and 16), with the exception of a few long-branching lineages (see below).

##### *1.1 Resolving deep evolutionary relationships between chlamydial clades*

Overall, within previously identified clades, our analyses recovered shallow evolutionary relationships that were consistent with recent work[3]. However, there are notable differences with regard to the inferred deeper evolutionary relationships. In particular, previous work has suggested that the Chlamydiaceae (denoted as the order Chlamydiales[3]) are deeply branching[1-4] and comprise a sister group of all other chlamydial lineages (corresponding to C-I, CC-II, CC-III, Anoxychlamydiales and environmental chlamydiae members)[2,3], which was tentatively classified as the order Parachlamydiales[3].

In contrast, all our phylogenomic reconstructions strongly support a sister relationship of the Chlamydiaceae with CC-IV, which together form a sister clade of the environmental chlamydiae. Altogether, this group forms a sister relationship with the second major radiation in the Chlamydiae, comprised of CC-I, CC-II, CC-III and Anoxychlamydiales lineages (Fig. 2, Supplementary Figs. 3 and 16).

Our results differ from prior analyses due to the inclusion of CC-IV, which is composed of three newly identified metagenome assembles genomes (MAGs) from Loki's Castle marine sediments, and the use of phylogenetic inference methods aimed at minimizing artifacts such as long-branch attraction (LBA). For instance, the branch leading to the Chlamydiaceae family is relatively long, which may in part be due to the evolutionary transition to a parasitic lifestyle with a restricted animal host range[6,7]. The inclusion of CC-IV in our analyses shortens the long branch to the Chlamydiaceae and may thus alleviate phylogenetic reconstruction artefacts that were previously attracting the latter to the base of the phylum.

Investigations of the evolution of the Chlamydiae and inferences on the nature of the chlamydial ancestor have been based on the assumption that the Chlamydiaceae represent the earliest diverging lineage within this phylum[1,2]. Thus, conclusions from these analyses will need to be re-examined based on the herein updated phylogeny of the Chlamydiae.

##### *1.2 Phylogenetic placement of long-branching chlamydial lineages*

In a recent study, the long-branching orphan lineage Chlamydiae bacterium RIFCSPHIGHO2_12_FULL_49_11 was inferred as the second deepest-branching lineage within Chlamydiae (after the divergence of *Ca.* Similichlamydia epinephelii) and was proposed to form the new order *Candidatus* Novochlamydiales[3].

In agreement with this, our initial maximum-likelihood (ML) phylogenies suggested the placement of Chlamydiae bacterium RIFCSPHIGHO2_12_FULL_49_11, followed by

K940_chlam_8 at the base of Chlamydiae, although support for the early divergence of these representatives was weak (BV = 42 and BV = 61, respectively) (Fig. 2, Supplementary Data 4). When 25% of the most heterogeneous sites were removed, both lineages became nested inside a larger clade composed of CC-I, II, III and Anoxychlamydiales, although with poor support (Fig. 2, Supplementary Data 4). However, in our Bayesian phylogenetic inference based on the CAT-GTR model, a complex model of protein evolution that minimizes the effects of LBA[5], the placement of the two lineages within the larger clade of CC-I, II, III and Anoxychlamydiales was highly supported (posterior probability (PP) = 0.97, Fig. 2). For instance, *Chlamydiae* bacterium RIFCSPHIGHO2_12_FULL_49_11 was placed within a well-supported clade with CC-I (PP = 0.99, Fig. 2), suggesting that the early divergence of this representative may indeed have been the result of LBA. Thus, our analyses indicate that *Chlamydiae* bacterium RIFCSPHIGHO2_12_FULL_49_11[2] does not represent a deep-branching *Chlamydiae* order but may instead be closely related to the Simkaniaceae family.

During the process of our analyses, several other chlamydial MAGs and Single-cell Assembled Genomes (SAGs) were publicly released (Supplementary Table 3). We reconstructed a ML phylogeny including these lineages, which was congruent with our prior analyses (Supplementary Fig. 3). One of these MAGs, representing *Candidatus* Similichlamydia epinephelii, was placed as a sister lineage to all other members of the Chlamydiae with high support (Supplementary Fig. 3). This position is consistent with other recent phylogenomic analyses of the Chlamydiae[2-4]. *Ca.* S. epinephelii is a member of the candidate chlamydial family *Candidatus* Parilichlamydiaceae, which is composed of chlamydial fish pathogens that cause epitheliocystis[6]. This taxon emerges on a long-branch, which is not surprising given the accelerated rate of evolution observed in many pathogens. Future phylogenetic analyses with an improved taxonomic sampling might better resolve the phylogenetic placement of *Ca*. Parilichlamydiaceae by alleviating potential phylogenetic artifacts.

##### *1.3 Genome characteristics and gene content variation across the Chlamydiae phylum*

Genome characteristics (e.g., genome size and GC content) and gene content vary widely between different clades of the Chlamydiae (Fig. 2, Supplementary Fig. 3). Nearly all genomic information available for CC-I, CC-II, CC-III and Anoxychlamydiales is represented by MAGs from Loki's Castle marine sediments (Supplementary Table 2) or other recent metagenomic surveys (Supplementary Table 3), which together represent over half of all chlamydial diversity. With the exception of *Simkania negevensis*, all characterized chlamydial lineages obtained through co-cultivation are part of the environmental chlamydiae and Chlamydiaceae (Supplementary Table 3).

CC-II has an unusually high GC-content among chlamydiae, and branches as a sister group to the *S. negevensis*-containing[7] CC-I clade (Fig. 2, Supplementary Fig. 3). The estimated genome sizes of CC-II MAGs derived from this metagenomic study and others (1.6-2.0 Mbp), and CC-I lineages and *Chlamydiae* bacterium K1060_chlam_2 (1.7 Mbp), are all distinctly smaller than the *S. negevensis* genome (2.6 Mbp), indicating differences in their cell biology and lifestyle. The genomes of CC-I and CC-II lineages appear to have experienced reductive genome evolution as their genomes display smaller median intergenic space in comparison to all other chlamydiae, particularly in comparison with the genomes of many environmental chlamydiae (Supplementary Fig. 3).

CC-III is composed solely of MAGs derived from recent metagenomic studies (Supplementary Table 3).

Anoxychlamydiales is dominated by marine sediment chlamydiae (Fig. 2, Supplementary Fig. 3), including nearly half of the MAGs obtained from Loki's Castle marine sediments,

*Chlamydiae* bacterium RIFCSPHIGHO2_12_FULL_27_8 (derived from Rifle Colorado aquifer groundwater[8]) and *Chlamydiae* bacterium SM23_39 (derived from the sulfate-methane transition zone of White Oak River sediments[9]). The Anoxychlamydiales are characterized by an exceptionally low GC content (26-31%) and have the largest median intergenic spaces of members of the sub-clades including CC-I, CC-II, CC-III and Anoxychlamydiales. Gene content within this clade is highly conserved in comparison with other newly identified chlamydial lineages (Supplementary Fig. 3).

Only two marine sediment chlamydiae MAGs appeared to belong to the environmental chlamydiae clade of well-characterized chlamydial symbionts of protists[10,11] (Fig. 2, Supplementary Fig. 3). *Chlamydiae* bacterium K940_chlam_3 represents the deepest branching lineage of the environmental chlamydiae, and *Chlamydiae* bacterium K940_chlam_7 represents a sister taxon to *W. chondrophila*[12]. These representatives have estimated genome sizes of 2.0 and 2.6 Mbp respectively, which is consistent with the range of genome sizes represented by other members of the environmental chlamydiae (2.1-3.4 Mbp). The environmental chlamydiae clade displays the most varied patterns in gene content and the largest median intergenic space across their genomes (mean of 84 bp) (Supplementary Fig. 3). These factors point to gene acquisition events, which could be a result of the amoeba-associated lifestyles of members of this clade[13].

CC-IV is composed solely of three marine sediment chlamydiae MAGs which have higher GC-content (47%) than other chlamydiae (mean of 39%) and forms a well-supported sister clade of the Chlamydiaceae (Fig. 2, Supplementary Fig. 3). The gene content within CC-IV representatives is less conserved than within members of the Chlamydiaceae (Supplementary Fig. 3). Furthermore, CC-IV members have larger estimated genome sizes (1.3-2.1 Mbp) than the Chlamydiaceae (1-1.2 Mbp), with *Chlamydiae* bacterium K940_chlam_9 having a particularly large genome size in comparison (2.1 Mbp).

#### 2. Insights into the evolution of pathogenicity in Chlamydiaceae

The Chlamydiaceae family, recently reviewed in[14,15], includes important animal and human pathogens. The well-known human pathogen *Chlamydia trachomatis* is the causative agent of sexually transmitted genital tract infections and trachoma (i.e., preventable blindness), while *Chlamydophila pneumoniae* can cause acute respiratory infections. In addition, *C. psittaci*, *C. abortus*, and *C. felis* all have zoonotic potential and can also cause disease in humans[14]. Based on our revised Chlaymidiae phylogeny and the expanded genomic sampling of members of this phylum, here we provide insights into the emergence and evolution of the Chlamydiaceae family.

##### *2.1 Chlamydiaceae evolved later in Chlamydiae evolution, and through genome reduction*

The Chlamydiaceae were thought to be an early-diverging group within the Chlamydiae phylum[1-4]. The expanded genomic sampling of chlamydial diversity and use of sophisticated phylogenomic methods herein has allowed us to propose a new phylogeny for the Chlamydiae, including the Chlamydiaceae (Supplementary Discussion 1). Specifically, we consistently recover a strongly supported sister-relationship between Chlamydiaceae and CC-IV, which together form a clade sister to the amoeba-associated environmental chlamydiae (Fig. 2, Supplementary Fig. 3, Supplementary Fig. 16).

The CC-IV clade is solely comprised of uncultured lineages identified in Loki's Castle marine sediments. When compared to Chlamydiaceae, CC-IV lineages have larger genomes (1.1-1.2 Mbp and 1.3-2.1 Mbp, respectively) and higher GC contents (37–41% and 47%, respectively). These two clades also differ significantly in their conservation of gene content (Supplementary Fig. 3) while the gene content of Chlamydiaceae is highly conserved, it is highly variable between the three obtained CC-IV lineages.

Our analyses of the presence and absence patterns of Non-supervised Orthologous Groups (NOGs) support previous reports which indicated that the evolution of the Chlamydiaceae family was characterized by massive gene loss[14,16-18], consistent with observed genome size reduction in the branch leading to this family (Fig. 2, Fig. 3a-b, Supplementary Data 3). When comparing gene content between environmental chlamydiae, CC-IV, and Chlamydiaceae, we found 576, 248 and 36 NOGs respectively, conserved uniquely within each clade (Fig. 3b,). When considering NOGs found exclusively in the Chlamydiaceae and not in other chlamydiae, the set of protein families uniquely conserved in this group dropped further to 15 (Supplementary Fig. 6). We also identified 13 PF domains conserved across Chlamydiaceae, which are not present in the genomes of other chlamydial lineages. The acquisition of the small set of proteins conserved in the Chlamydiaceae which are not found in other chlamydiae, may have played a role in the evolution of the clade. In addition, the proportion of NOGs assigned to each Cluster of Orthologous Groups of proteins (COG) functional category was generally smaller in Chlamydiaceae relative to environmental chlamydiae and CC-IV lineages (Supplementary Fig. 5). The latter observation was most prevalent in COGs with the largest underrepresentation in functional categories related to metabolism (e.g., energy production and conservation, carbohydrate transport and metabolism, and inorganic ion transport and metabolism; Supplementary Fig. 5). Indicating, that in particular a loss of functions related to metabolism may have contributed to Chlamydiaceae evolution.

##### *2.2 Chlamydiaceae display reduced metabolic capacities relative to CC-IV*

Several metabolic pathways appear to have been lost specifically in Chlamydiaceae relative to CC-IV and environmental chlamydiae (Supplementary Fig. 4, Supplementary Data 3). These pathways include proline biosynthesis, and the UMP biosynthesis pathway necessary for pyrimidine biosynthesis (KEGG module: M00051). Furthermore, Chlamydiaceae members lack genes for a hexokinase (or any glucokinase) and for the first three enzymes of the tricarboxylic acid cycle (TCA) cycle (i.e., citrate synthase, aconitase and isocitrate dehydrogenase)[19]. Consequently, they depend on their host for metabolic exchange of TCA cycle intermediates and glucose-6-phosphate[19]. In contrast, a glucokinase and a complete TCA cycle are found in virtually all genomes of environmental[19] and CC-IV chlamydiae. These patterns suggest that all pathways mentioned above were present in the common ancestor of environmental chlamydiae, CC-IV and Chlamydiaceae, and subsequently lost in Chlamydiaceae.

Many flagellar components are present in CC-IV and individual chlamydial lineages branching at the base of the environmental chlamydiae and CC-IV/Chlamydiaceae clades while only a few components were identified in Chlamydiaceae (Supplementary Discussion 3). Phylogenetic analyses of these flagellar components indicate that these were already present in the last common ancestor of Chlamydiaceae, environmental and CC-IV chlamydiae (Supplementary Fig. 8, Supplementary Data 4, Supplementary Discussion 3). Yet, the few subunits present in the Chlamydiaceae seem to have been co-opted to act alongside their NF-T3SS[20].

##### *2.3 Gain of virulence and host interaction factors in Chlamydiaceae evolution*

In comparison to other lineages of the Chlamydiae, Chlamydiaceae genomes are characterized by a set of unique and functionally annotated core genes (Fig. 3b, Supplementary Fig. 6) that encode proteins associated with host-interaction and virulence.

In particular, and in agreement with previous studies[17], we observed an expansion of Polymorphic Outer Membrane Protein families (POMPs) uniquely in members of the Chlamydiaceae[14]. Polymorphic outer membrane proteins (POMPs) allow for niche-specific adhesion of chlamydial cells to their animal hosts, and also aid in immune system evasion through their antigenic diversity[1]. Several additional outer membrane proteins are also conserved across Chlamydiaceae (Supplementary Fig. 6).

Another striking example of a protein uniquely conserved among members of the Chlamydiaceae is the carbohydrate-selective porin (OprB) protein (Supplementary Fig. 6), which is a component of the outer membrane complex of membrane proteins that are surface-exposed in Chlamydiaceae EBs[21]. Besides, all Chlamydiaceae encode one arginine decarboxylase which most likely functions in the reduction of arginine reserves during host cell infection, but could also protect from nitrosative stress[22]. Furthermore, most Chlamydiaceae uniquely encode a Membrane Attack Complex PerForin (MACPF)[14,23]. While the exact function of the MACPF in Chlamydiaceae is unclear, it may assist in the acquisition and processing of lipids derived from the host, play a role in host immune system avoidance or facilitate host entry through pore formation[14].

Finally, Chlamydiaceae are characterized by a large number of highly conserved NOGs and protein family (PF)[24] domains with unknown function (Supplementary Fig. 6), some of which could play a role in the pathogenic lifestyle of members of this family.

##### *2.4 Gene acquisition events unique to CC-IV and Chlamydiaceae*

Members of the CC-IV and Chlamydiaceae (Supplementary Fig. 7a) appeared to encode seven gene families (by NOG or PF domain) absent in all other Chlamydiae lineages, which were likely gained prior to the divergence of the two former clades. Despite their presence in all representative Chlamydiaceae genomes investigated here, the function of most of these proteins is unknown (barring the exception of COG0400 in the genome of *Chlamydia* sp. 2742-308). Their maintenance across Chlamydiaceae, despite massive gene loss during evolution of this family (Fig. 3a-b), suggests that these proteins play important roles in their pathogenic lifestyles. To further investigate their evolutionary history, we inferred single-gene tree phylogenies for two of these protein families (PF04518 and PF05302), which are thus far taxonomically restricted to CC-IV and Chlamydiaceae (see Methods).

While only one protein with the PF domain PF04518 is found in CC-IV member *Chlamydiae* bacterium K940_chlam_9, genomes of Chlamydiaceae encode four or five proteins with this domain (Supplementary Fig. 7a). A phylogenetic analysis of proteins assigned to PF04518 (Supplementary Fig. 7a) revealed that this gene family appears to have undergone several gene duplication events, after the divergence of CC-IV and Chlamydiaceae and prior to the diversification of the latter, resulting in four distinct gene copies (Supplementary Fig. 7b). Each of these copies belongs to one of four different highly supported clades (BV > 98) (Supplementary Fig. 7b). Genes encoding proteins from clades 1 and 2, as well as from clade 3 and 4, are localized, respectively, in a gene cluster in the genomes of Chlamydiacaeae. A subset of proteins assigned to cluster 4 experienced an additional gene duplication event in *Chlamydia trachomatis*, *Chlamydia muridarum* and *Chlamydia suis*, and form a distinct sub-clade (clade 5) within clade 4 (Supplementary Fig. 7b). The previous investigation of this protein family in Chlamydiaceae has shown that its members contain a Non-Flagellar Type III Secretion System (NF-T3SS) signal[25] and appear to be secreted by the NF-T3SS as effectors[26]. They may act by targeting nuclear functions, since they are found in the nucleus of infected host cells[25,26]. The function of proteins with the PF04518 domain in Chlamydiaceae was likely neo-functionalized by the above described duplication events. Understanding the function of the single copy protein in Chlamydiae bacterium K940_chlam_9 could help determine the ancestral function of the protein and how it impacted the evolution of Chlamydiaceae pathogenicity.

We also observed conserved gene duplications between the CC-IV and Chlamydiaceae in the case of proteins with the domain PF05302 (Supplementary Fig. 7a). In this case, a phylogenetic analysis revealed three distinct clades, with one copy from *Chlamydiae* bacterium K940_chlam_9 and Chlamydiaceae members found in each (Supplementary Fig. 7c). All three copies are organized together in the genomes of both *Chlamydiae* bacterium K940_chlam_9 and Chlamydiaceae members (Supplementary Fig. 7c). Together, these results indicate that this gene family underwent several gene duplication events prior to the divergence of CC-IV and Chlamydiaceae. *Chlamydia trachomatis* PF05302 domain-containing homologs CT847 (clade 3) and CT849 (clade 1) (Supplementary Fig. 7c, Supplementary Data 4) have both been characterized as T3SS substrates, and likely effectors[27,28]. *Chlamydia trachomatis* homolog CT847 appears to interact with mammalian Grap2 Cyclin D-Interacting Protein (GCIP), a protein involved in the eukaryotic cell cycle[27]. Examining the role of the three proteins with the domain PF05302 in *Chlamydiae* bacterium K940_chlam_9 could help in elucidating their ancestral functions. Thereby aiding in understanding the contributions of this protein family to Chlamydiaceae evolution.

Future, more fine-grained investigations of gene content evolution in members of the Chlamydiae, with the inclusion of CC-IV members, will be crucial to better understand the evolutionary trajectories that led to the ecological success of Chlamydiaceae as animal pathogens.

#### 3. Secretion systems and flagella in Chlamydiae

##### *3.1 Detection of secretion systems, flagella and effectors*

The secretion of proteins and other molecules by secretion systems is important for host association, microbial interactions and relation with the environment. We screened all available Chlamydiae genomes for type I to VI secretion systems (T1SS to T6SS), flagella and related genes with MacSyFinder[29] (see Supplementary Data 3). Most chlamydiae genomes were found to contain genes for T1SS, T2SS, T3SS and T5SS, while only a few encoded T4SS and flagella genes. We discuss each of these systems below, following the gene nomenclature proposed by Abby *et al.*[30].

**T1SS.** T1SSs are simple one-step protein secretion systems that are formed by three components: an inner membrane ABC transporter, an outer membrane component and a bridging membrane fusion protein. All three components were found in most of the surveyed chlamydiae. However, the membrane fusion protein was not detected in most CC-IV and CC-III lineages, and all three components were absent in most Chlamydiaceae (Supplementary Data 3).

**T2SS.** T2SS are complex protein secretion systems formed by outer membrane, inner membrane and pseudopilus apparatuses, and a cytoplasmic ATPase[31,32]. The most commonly detected homologs for this system in Chlamydiae were GspD, GspF and GspG, respectively, which represent the central core proteins of the three above-mentioned T2SS structural apparatuses. We also detected the cytoplasmic ATPase GspE in a few Chlamydiae genomes, and PilB (a homolog of GspE in type 4 pili) was often detected in those genomes where GspE was missing (Supplementary Data 3). The core genes *gspDEFG* were generally co-located in tandem (Supplementary Fig. 10). Other T2SS components, such as the minor pseudopilins GspHIJK (labeled as 'mandatory' by MacSyFinder) and other non-essential proteins (labeled as 'accessory' by MacSyFinder) were often absent. However, situated immediately upstream of *gspDEFG*, we detected either the minor pseudopilin genes *gspHIJK*, or genes of similar length patterns with a significant e-value using BLAST (Supplementary Fig. 10). Taken together, these results suggest that chlamydiae harbour a variant of the classical T2SS. Furthermore our observations agree in part with Peabody *et al.*[31], who described the presence of genes *gspCDEFG* in *Chlamydia* and *Chlamydophila* genomes: we were unable to detect GspC, while Peabody *et al.* were unable to detect the minor pseudopilins GspHIJK.

**Non-flagellar T3SS (NF-T3SS), flagellum and T3SS-secreted effectors.** NF-T3SSs are complex protein secretion systems generally with eukaryotic host interactions, and have previously been shown to be essential for virulence in Chlamydiaceae (reviewed in e.g.,[33,34]). NF-T3SS components evolved through exaptation of proteins constituting the bacterial flagellum[35], which complicates their unambiguous detection and annotation. The components screened in the present study include the outer membrane ring secretin (SctC), the inner membrane ring (SctJ), the secretion apparatus (SctRSTUV), the sorting platform (SctQ) and the cytoplasmic ATPase (SctN). SctC is unique to the NF-T3SS, while the other components share homology with flagellar proteins. To evaluate whether the chlamydial homologs were NF-T3SS or flagellar genes, we performed phylogenetic analyses of various individual genes, as well as of concatenated alignments, using as reference the SctN sequences published by Abby and Rocha[35] and the other discussed NF-T3SS sequences used in Abby *et al.*[30]. Similar to previous studies[35], our phylogenetic analyses (Supplementary Fig. 8, Supplementary Data 4) place a myxococcal NF-T3SS system as sister to all other bacterial NF-T3SS sequences. The latter then diverge on one hand into all of the chlamydial sequences and, on the other, the rest of bacteria. Our analyses retrieve the monophyly of the main chlamydial clades, which is in overall agreement with the species tree. These results confirm that the NF-T3SS is found across Chlamydiae.

The NF-T3SS genes are distributed over three gene clusters, one containing *sctN*, *sctQ* and *sctC*, one containing *sctJ*, *sctR*, *sctS* and *sctT*, and one containing *sctU* and *sctV* (Supplementary Fig. 9). The gene order and neighborhood of these clusters is highly conserved in all Chlamydiae genomes, as has been shown previously for environmental chlamydiae and Chlamydiaceae[8,9]. The gene order conservation allowed us to detect *sctC* homologs in many genomes where this gene was not detected by MacSyFinder: we identified significant BLAST hits to known SctC sequences in their expected position near the *sctN* and *sctQ* genes. The three gene clusters were interspersed and flanked with other conserved genes on the same strand. While their function could not be determined in most cases, a gene situated between *sctQ* and *sctC* encoded a serine/threonine protein kinase that has been suggested to participate in NF-T3SS protein secretion[10]. Altogether, we hypothesize that the NF-T3SS genes were acquired by the common ancestor of Chlamydiae and have since been inherited vertically.

In contrast to the ubiquity of NF-T3SS, we found flagellar genes only in a handful of genomes, including CC-IV and four marine chlamydiae related to the environmental chlamydiae and Chlamydiaceae clades[36,37]. Although many chlamydiae genomes were found to contain putative homologs of the flagellar proteins *sctN* and *sctQ* genes, most turned out not to be associated with flagellar function. On one hand, most proteins detected as flagellar SctQ homologs branched instead with NF-T3SS genes in phylogenetic analyses (ufBV=94%; SH-aLRT=92%) or within a clade extremely distantly related to flagellar homologs (Supplementary Data 4). Phylogenetic analyses of chlamydial SctN homologs similarly revealed that these often branched with non-flagellar ATPases (Supplementary Data 4). However, phylogenetic analyses were inconclusive regarding the putative function of a group of proteins annotated as flagellar SctN homologs by MacSyFinder in Chlamydiaceae: while they are more closely related to flagellar sequences than to other homologs, they are not nested within them (Supplementary Data 4). Putative flagellar homologs of SctV in Chlamydiaceae also form long-branching clades related to known flagellar sequences, indicating these proteins could represent divergent flagellar homologs (Supplementary Data 4). Remarkably, flagellar homologs of SctN (FliI) and SctV (FlhA) in Chlamydiaceae have been shown to interact with the NF-T3SS protein complex, suggesting they have been co-opted to a new function in protein secretion[20].

In contrast, the above-mentioned marine chlamydiae and the CC-IV *Chlamydiae* K940_chlam_9 and KR12_chlam_2 were found to contain a large array of flagellar genes (Supplementary Figs. 4 and 8; Supplementary Data 3). Even though CC-IV *Chlamydiae* bacterium K1000_chlam_4 contained only a copy of the flagellar homolog of *sctR (fliP)*, it is possible that this genome encodes a full flagellar gene set, since this MAG is relatively incomplete (Fig. 2, Supplementary Table 2) and the contig containing *fliP* ends right after this gene, and thus before the expected location of the flagellar homologs of *sctS (fliQ)* and *sctT (fliR)* genes. The flagellar genes of CC-IV and the marine chlamydiae listed above form a well-supported clade in the phylogeny of concatenated NF-T3SS genes and in single-gene trees of SctJ, SctR (Supplementary Fig. 8), SctN and others (Supplementary Data 4). Therefore, given the species phylogeny obtained in the present study (Fig. 2, Supplementary Fig. 3), these results suggest that gene sets for the flagellum were present in the common ancestor of the environmental chlamydiae, Chlamydiaceae and CC-IV clades, but were ultimately lost in the former two groups but retained in their CC-IV and unclassified relatives. However, further phylogenetic analyses that adequately model the extreme divergence of the Chlamydiaceae genes of putative flagellar origin will be required to verify these inferences.

Since we were able to predict the existence of a NF-T3SS in several chlamydiae, we used EffectiveDB[38] to predict T3SS-secreted proteins, eukaryotic-like domains (ELD), and putative subcellular targeting signals to eukaryotic cellular compartments. None of the non-chlamydial PVC bacteria were predicted to have any T3SS secreted proteins. In contrast, 9% to 28% of chlamydial proteomes were predicted to possess a T3SS-associated signal peptide. However, we could not identify major differences between various subclades of chlamydiae regarding most features predicted by EffectiveDB (Supplementary Data 3). A notable exception concerns the prediction of CCBD (conserved chaperone-binding domain) motifs, which are usually found in the N-terminal region of T3SS-secreted proteins and have been shown to serve as binding site of chaperones facilitating the correct selection and unfolding of T3SS-dependent effector proteins. We found that Anoxychlamydiales members were enriched in proteins predicted to have a CCBD motif (average $4.58 \pm 0.78\%$) compared to other chlamydiae (2.52

± 1.20%). However, the significance of this result is difficult to assess given that these lineages do not appear to be enriched in predicted T3SS-secreted proteins. Intriguingly, five chlamydiae were not predicted to have any T3SS-secreted proteins, although all of these organisms are predicted to have a NF-T3SS. In addition, *Verrucomicrobium spinosum*, a verrucomicrobium recently described to have a NF-T3SS[39], was also not predicted to have any T3SS-secreted proteins or CCBD motifs. This suggests that predictive tools such as EffectiveDB are currently unable to model the entire diversity of proteins motifs that are recognized by the T3SS and their chaperones, and that differences between chlamydial lineages in terms of their T3SS-secreted proteins will have to be revisited when more sensitive predictive tools become available.

**T4SS.** T4SSs are versatile systems generally involved in contact-dependent translocation of proteins and DNA. We detected most of the Type F T4SS genes in a subset of Chlamydiae genomes: *Waddliaceae* bacterium SP13, *Chlamydiae* bacterium K1060_chlam_2 (CC-I), *Chlamydiae* bacterium K1000_chlam_3 (CC-II), *R. massiliensis*, *Parachlamydia* spp. and *Protochlamydia* spp (Supplementary Fig. 11). This patchy distribution of T4SS genes in chlamydiae is consistent with the idea that these genes were recently acquired via horizontal gene transfer (HGT).

**T5SS**. T5SS are two-step protein secretion systems, generally substrate-specific and containing one to three components. T5SS classical autotransporters (type 5a secretion systems, T5aSS) and translocators (type 5b secretion system, T5bSS) were identified in most CC-I, II, III and Anoxychlamydiales members, while most environmental chlamydiae and Chlamydiaceae only contained T5aSS autotransporters.

**Other systems**. Finally, while a few homologs were detected for other systems such as T6SS, Tad pili and type IV pili (T4P), these remained largely incomplete in all genomes, indicating these inferences likely represent false positives.

##### *3.2 Putative functions of secretion systems in marine sediment Chlamydiae*

The observation made above largely corroborates findings made in previous studies, which indicate that most Chlamydiae contain T1SS, T3SS and T5aSS, and provide new evidence for the presence of T2SS and the sparse presence of T4SS and flagella[3,17,30,40]. The presence of secretion systems is commonly interpreted in the light of host-symbiont dynamics. However, the lack of identified eukaryotes in the presented samples (see Supplementary Discussion 6) raises the possibility that these perform alternative functions in at least some of the newly discovered chlamydial lineages.

In Chlamydiaceae, T2SS, T3SS and T5aSS are typically linked to host adhesion, invasion and manipulation[15,41,42]. Similarly, environmental chlamydiae have also been shown to express secretion systems during infection of microbial eukaryotes[43,44]. However, whether their function is conserved throughout the Chlamydiae phylum remains unclear. For example, a recent transcriptomic study found that the expression of T3SS is higher in reticulate bodies than in elementary bodies in *C. abortus*, but found the opposite pattern in *W. chondrophila*[44]. The exact nature of the cell cycle and the function of secretion systems in these lineages are yet to be elucidated. Furthermore, the extracellular stage of the chlamydial cell cycle is considerably understudied[45], and little is known about potential chlamydial interactions with other microbes.

Despite the traditional link between secretion systems and host-association, some of these systems have been described to target prokaryotes. For example, T6SS are typically used for bacteria-bacteria interactions[46], and gram positive T7SS have been described to target bacteria under certain conditions[47]. T1SS, T4SS and T5bSS, all present in chlamydial genomes, have also been shown to target bacterial cells[48-51]. In *Legionella pneumophila*, T2SS and T4P facilitate biofilm formation and retention[52,53], and the former is involved in sliding motility[54] and extracellular survival in freshwater. Taken together, this suggests that the presence of various secretion systems does not necessarily imply interactions with eukaryotic hosts.

In line with this, NF-T3SS have also been described in bacteria that are not known to interact with eukaryotes[55]. For example, a NF-T3SS has been identified in *Verrucomicrobium spinosum*, a generally free-living organism (though it has been shown to have detrimental effects when experimentally inoculated in fruit flies and *Caenorhabditis elegans*[39]). NF-T3SS has also been found alongside T6SS, chemotaxis and flagellar genes, in strains of *Vibrio* and *Aeromonas* associated with microbial biofilms, but which are not known to associate with eukaryotes[39]. The presence of NF-T3SS in Myxococcales is particularly interesting, given that they are not associated with a host and contain a highly divergent version of the NF-T3SS, which represents a sister to all other NF-T3SS and lacks various genes generally associated with this system[35]. The chlamydial version of the NF-T3SS, which is placed as sister clade to all non-myxococcal NF-T3SS (Supplementary Fig. 8) may hold functional similarities with the more divergent Myxococcales NF-T3SS[35].

In conclusion, these observations indicate that the presence of various secretion systems does not necessarily imply a eukaryote-associated lifestyle. Alternatively, proteins secreted by secretion systems could play a role in growth in biofilms, in the interaction with other microbial groups or in the modification of the environment. Studies about the biology of chlamydial elementary bodies, as well as visualisation of representatives of the newly discovered lineages will be instrumental to answer this question.

**4. Phylogenetic diversity of chlamydial nucleotide transporters.**

Nucleotide transporters (NTTs) belong to the 'ATPases Associated with diverse cellular Activities' (AAA) family of proteins[56] and can transport a range of metabolites, including ATP, the cofactor nicotinamide adenine dinucleotide ($NAD^+$), ribonucleotides and deoxyribonucleotides across a membrane. NTTs are found in diverse lineages in the tree of life. In plastid-bearing eukaryotes, the ATP/ADP NTT proteins are essential for the import of ATP into the organelle from the cytosol[57,58]. Some obligate intracellular pathogenic eukaryotes (e.g., Microsporidia[16,56]) and obligately symbiotic bacteria (e.g., members of the Chlamydiaceae and *Rickettsia*[59]) use NTTs to import ATP and other nucleotides from their eukaryotic hosts. A recent investigation of NTT phylogenetic diversity found that NTT homologs are also found in a diverse set of free-living organisms[60].

All chlamydial genomes investigated to date, including those from marine sediment chlamydiae, encode multiple NTT homologs (Supplementary Fig. 4, Supplementary Data 3), although the number of homologs varies across different representatives of this phylum. Depending on the clade, we observed two (CC-IV and Chlamydiaceae), four (CC-I, CC-II and environmental chlamydiae) or five (Anoxychlamydiales) distinct NTT paralogs (Supplementary Data 3). To classify the NTTs of marine sediment chlamydiae related to those from characterized lineages, we reanalysed the phylogenetic diversity of NTT proteins across the tree of life.. For this, we used a previous analysis that resolved the NTT superfamily[60]: "canonical NTTs", "other NTTs" and in addition, proteins with NTT-HEAT domains.

***4.1 Phylogenetic diversity of "canonical NTTs"***

"Canonical NTTs" have a single TLC (PF03219) protein domain architecture, and include most functionally characterized NTTs, such as the ATP/ADP transporters from plastids, Microsporidia, *Rickettsia* and Chlamydiae (Supplementary Fig. 12a). Phylogenetic analyses of these "canonical NTTs" resolved nine distinct groups of chlamydial sequences (Supplementary Fig. 12a). Most ATP/ADP transporters of primary and secondary plastid-bearing lineages (e.g., archaeplastids, diatoms, haptophytes, red algae and brown algae) formed a strongly supported clade (ufBV = 95, Supplementary Data 4). Notably, in spite of the cyanobacterial origin of plastid[61], the closest prokaryotic homologs to plastid derived ATP/ADP transporters are represented by homologs of Chlamydiae (Supplementary Fig. 12a). As previously hypothesized[59,60,62,63], this suggests that plastid-bearing lineages acquired the NTT gene from a chlamydial-like donor early in the evolution of this organelle. These chlamydial ATP/ADP translocases formed a clade (ufBV = 91, Supplementary Data 4), referred to as cluster 9 (Supplementary Fig. 12a). Cluster 9 contains only one representative sequence from each major chlamydial clade, except for Anoxychlamydiales members, each of which has two paralogs. The topology within clade 9 (Supplementary Data 4) is consistent with the organismal phylogeny (Fig. 2), suggesting that this protein has evolved vertically within the phylum. The substrate specificities of several proteins in cluster 9 have been experimentally characterized. *Chlamydia trachomatis* (*Ct*)[56], *Candidatus* Protochlamydia acanthamoebae (*Pam*)[59] and *Simkania negevenis* (*Sn*)[59,64] encode ATP/ADP-specific NTT1 antiporters that participate in energy parasitism during infection of their hosts. Interestingly, *Ct*NTT1[65] can also act as an $NAD^+$/ADP antiporter, providing a mechanism through which members of the Chlamydiaceae can acquire $NAD^+$, which they are unable to synthesize[66],[67].

Clusters 6 and 7, were each composed of several representatives of the environmental chlamydiae, and branched closely to alphaproteobacterial lineages, including ADP/ATP transporters from *Rickettsia*[59] suggesting they may have similar functional roles in chlamydiae. Cluster 8 includes CC-I, one representative from Anoxychlamydiales, and several representatives from the environmental chlamydiae (Supplementary Fig. 12a). *Sn*NTT3, found in cluster 8, has been characterized and is a proton-independent general NTP transporter, which is also capable of transporting the deoxyribonucleotide triphosphate dCTP[64].

Five different chlamydial clusters, clusters 1-5, together formed a well-supported group (ufBV = 100). Cluster 5 includes CC-II and environmental chlamydiae (Supplementary Fig. 12a) and *Pam*NTT2, a proton-independent transporter of all four canonical ribonucleoside triphosphates[68]. Cluster 4 is composed solely of sequences from environmental chlamydiae (Supplementary Fig. 12a) and includes *Pam*NTT3, a proton-energized symporter that transports

UTP[68]. Cluster 3 is composed of members of CC-I, CC-II, CC-III, Anoxychlamydiales and includes *Sn*NTT2, a proton-dependent symporter of GTP and ATP[64]. Cluster 2 NTTs comprises sequences from CC-IV and Chlamydiaceae, and includes *Ct*NTT2, a proton-driven symporter of all four NTPs[56] (Supplementary Fig. 12a). Cluster 1 contains homologs from environmental chlamydiae, Anoxychlamydiales and some members of CC-II including *Pam*NTT5, a proton-energized symporter that transports both GTP and ATP[68] (Supplementary Fig. 12a).

Finally, we observe a clear functional conservation within the experimentally characterized $H^+$-driven symporters, all of which group together in clusters 1, 2, 3 and 4 (ufBV = 88 Supplementary Fig. 12a, Supplementary Data 4). This suggests that phylogenetic reconstructions of NTT homologues possess a predictive power in terms of mode of transport, but not substrate specificity.

##### *4.2 Phylogenetic diversity of "other NTTs" with a single-domain architecture*

A bacterial-dominated group of "other NTTs" has been described to have a single TLC (PF03219) protein domain architecture, and most proteins in this group have yet to be functionally characterized (Supplementary Fig. 12b). In the phylogenetic analysis of those "other NTTs", we recovered two groups of chlamydial sequences (Supplementary Fig. 12b). Cluster 11 includes representatives from environmental chlamydiae and Anoxychlamydiales (Supplementary Fig. 12b) and branches as a sister clade to a clade comprising sequences from the *Candidatus* Dependentiae (formerly TM6) phylum (Supplementary Fig. 12b). *Ca.* Dependentiae have reduced genomes and are thought to lead host-associated lifestyles with eukaryotic hosts[69]. Cluster 10 includes members of CC-I (including *Sn*NTT4, whose substrate specificity could not be determined in a prior study[64]), CC-II, and Anoxychlamydiales and forms a maximally supported clade composed largely of Deltaproteobacteria, including several *Bdellovibrio*-and-like-organisms (BALOs). BALOs have a predatory lifestyle whereby they invade the periplasm of other gram-negative bacteria to harvest nutrients[70]. However, this invasive lifestyle is not obligate, as BALOs can grow axenically under nutrient rich conditions.

##### *4.3 Phylogenetic diversity of "NTT-HEAT" proteins*

"NTT-HEAT" family NTTs[60] have an additional C-terminal HEAT domain (PF13646, PF02985) and are found across a wide-range of free-living bacteria, though none have been functionally characterized thus far (Supplementary Fig. 12c). HEAT domains are involved in protein-protein interactions[71], and thus may alter the function of the NTT domain in NTT-HEAT proteins[60]. These putative NTTs are hypothesized to facilitate inter-microbial nutrient exchange during bacteria-bacteria interactions[60]. For example, these NTTs could be involved in multicellular development in Cyanobacteria, which have proteins with this domain architecture[60]. A ML phylogeny of proteins with this domain architecture (Supplementary Fig. 12c) recovered one chlamydial clade, cluster 12, which includes the Chlamydiaceae, CC-IV, CC-III, some environmental chlamydiae and several members of Anoxychlamydiales.

##### *4.4 NTTs in marine sediment chlamydiae*

NTTs identified in marine sediment chlamydiae from this study clustered together with other chlamydial homologs (Supplementary Fig. 12). Despite distinct phyletic distribution patterns of the NTTs found in different chlamydiae clades, the ubiquity of NTT homologues in different chlamydial lineages gives weight to the proposed ancient origin of NTTs within the Chlamydiae phylum[59,60]. Marine sediment chlamydiae, appear to host a similar set of NTTs as other members of the phylum, including homologs closely related to functionally characterized NTTs. Due to the promiscuous functions of NTTs however[59,67], we cannot predict the substrate specificity of homologs found in marine sediment chlamydiae. Although they have homologs of the canonical ATP/ADP transporter (Supplementary Fig. 12a), functional characterization is necessary to determine their substrate specificity.

#### 5. Genomic potential for de novo biosynthesis of nucleotides and amino acids across Chlamydiae

Many host-associated bacteria, and particularly obligate intracellular bacteria, are able to acquire essential amino acids and nucleotides from their hosts. Obligate symbionts often undergo genome reduction and lose the ability to produce these compounds *de novo*[72]. Here, we discuss the ability of the marine sediment chlamydiae genomes for *de novo* amino acid and nucleotide biosynthesis (Supplementary Fig. 4, Supplementary Data 3).

##### *5.1 De novo biosynthesis of amino acids*

Similar to characterized chlamydiae[19], the investigated MAGs seem to be generally auxotrophic for many amino acids. In fact, no chlamydiae representative with the capacity to synthesize all amino acids has been identified thus far. Below, we discuss some examples of more extensive amino acid and nucleotide biosynthesis capabilities in specific Chlamydiae lineages.

**Proline.** Environmental chlamydiae do not have the coding potential to synthesize all amino acids *de novo*[19], but generally encode a larger set of amino acid biosynthetic capabilities than Chlamydiaceae (e.g., proline biosynthesis). We identified all genes for proline *de novo* biosynthesis in more than half of environmental chlamydiae genomes (9/15; including marine sediment lineage *Chlamydiae* bacterium K940_chlam7), and in one member of CC-IV (*Chlamydiae* bacterium K940_chlam_9). This observation is in line with a scenario in which proline biosynthesis was present in the common ancestor of the CC-IV, Chlamydiaceae and environmental chlamydiae, and was subsequently lost in the Chlamydiaceae.

**Aromatic amino acids.** The ability to synthesize aromatic amino acids (i.e., tryptophan, phenylalanine, and tyrosine) displays a punctuated distribution among the analysed chlamydiae. Seemingly, only *S. negevensis* is capable of synthesizing all three amino acids[19]. The genomes of *Chlamydiae* bacterium RIFCSPLOWO2_02_FULL_45_22 and its close relatives (Supplementary Fig. 13, Supplementary Data 3) encode the potential for phenylalanine and tyrosine biosynthesis and a near-complete pathway for tryptophan biosynthesis. While *Waddliaceae* bacterium SP13, *Chlamydiales* bacterium SCGC AG-110-P3 and *Parachlamydia* sp. C2 each encode a complete pathway for the biosynthesis of tryptophan, some Chlamydiaceae[19] were found to encode a near-complete pathway (Supplementary Fig. 4).

**Other amino acids.** Chlamydiae generally do not encode pathways for the biosynthesis of arginine, methionine, histidine, leucine, isoleucine and valine. We identified some notable exceptions, including a complete leucine biosynthesis pathway in *Parachlamydia* sp. BC.030 and a near-complete histidine biosynthesis pathway (7/8 components; Supplementary Data 3) in *Chlamydiae* bacterium RIFCSPLOWO2_02_FULL_45_22 and close relatives (Supplementary Fig. 13, Supplementary Data 3).

##### *5.2 De novo biosynthesis of nucleotides*

**Pyrimidine.** Chlamydiaceae and most members of the environmental chlamydiae are auxotrophic for the *de novo* biosynthesis of both purine and pyrimidine nucleotides[19]. Previous work has identified pyrimidine biosynthesis (i.e., uridine monophosphate (UMP) biosynthesis from glutamine; KEGG module M00051)[73,74] in *W. chondrophila* WSU 86-1044[25] and *Criblamydia sequanensis* CRIB-18[73,74]. We identified a complete pyrimidine biosynthesis pathway in members of CC-II (*Chlamydiae* bacterium Ga0074140), CC-III (*Chlamydiae* bacterium CG10_big_fil_rev_8_21_14_0_10_42_34, CG10_big_fil_rev_8_21_14_0_10_35_9), CC-IV (*Chlamydiae* bacterium K940_chlam_9) and environmental chlamydiae (*Chlamydiae* bacterium K940_chlam_7, K940_chlam_3, *Chlamydiales* bacterium SCGC AG-110-M15 and *Waddliaceae* bacterium SP13). Further, we identified near-complete pathways in additional chlamydiae members of CC-II (*Rhabdochlamydia helvetica* T3358 and *Chlamydiae* bacterium K940_chlam_2), other CC-IV lineages, and environmental chlamydiae (*Chlamydiae* bacterium K940_chlam_3).

**Purine.** Compared to pyrimidine biosynthesis, purine biosynthesis is more sparsely distributed among the Chlamydiae. The first evidence for a near-complete *de novo* purine biosynthesis pathway (i.e., inosine monophosphate (IMP) biosynthesis from glutamine; KEGG module M00048) in Chlamydiae was recently described in *Chlamydiales* bacterium SCGC AG-110-M15[36]. We additionally identified the complete pathway for IMP biosynthesis in *Chlamydiae* bacterium CG10_big_fil_rev_8_21_14_0_10_35_9 and *Waddliaceae* bacterium SP13, and partial biosynthesis pathways (i.e., all but the PurE and PurK encoding genes; Supplementary Data 3) in *Chlamydiales* bacterium SCGC AG-110-M15 and *Chlamydiae* bacterium K940_chlam_3 (environmental chlamydiae). The former represents a relatively incomplete SAG (Supplementary Table 3), such that the presence of these genes cannot be ruled out.

Interestingly, all genomes which encode a complete or near-complete purine *de novo* biosynthesis pathway also encode a complete or near-complete pathway for *de novo* pyrimidine biosynthesis. This observation suggests that some chlamydiae might not rely on a host for these essential metabolites. Future analyses aimed at inferring the evolutionary histories of these pathways will help to determine whether nucleotide biosynthesis was ancestrally present in Chlamydiae or rather represents a derived trait acquired by horizontal gene transfer.

#### 6. Eukaryotes in Loki's Castle marine sediments

All chlamydiae characterized to date represent obligate symbionts with eukaryotic hosts and have a characteristic biphasic lifecycle (intracellular host-associated phase and extracellular elementary body phase). Our analyses revealed that the marine sediment chlamydiae encode key host-association features (e.g., NF-T3SS; Supplementary Data 3, Supplementary Discussion 3) and elementary body factors (e.g., early upstream reading frame transcription factor and histone-like development protein; Supplementary Fig. 4, Supplementary Data 3) and are predicted to be auxotrophic for some nucleotides and amino acids (Supplementary Fig. 4, Supplementary Data 3, Supplementary Discussion 5). These observations would suggest that marine sediment chlamydiae might be host-associated and prompted a thorough search for eukaryotes in these sediments.

Indeed, active populations of fungi, protists and macrofauna have been observed in marine sediments[75-78]. Using universal eukaryotic primer sets, we failed to amplify 18S rRNA gene sequences from the marine sediment samples (Supplementary Table 5), in line with previous analyses of Loki's Castle marine sediments[79]. However, we were able to identify several 18S rRNA gene sequences in the obtained metagenomic data (see below; Supplementary Table 4), suggesting that eukaryotes might represent low-abundant community members of these anaerobic marine sediments. Yet, it has been shown that eukaryotic DNA from overlying water columns can be deposited and well-preserved in marine sediments under anoxic conditions[80,81]. In the present study, we were unable to determine whether the observed eukaryotic DNA sequences were derived from live cells capable of hosting chlamydiae. Below we expand on the identified 18S rRNA gene sequences and discuss their potential sources.

Several 18S rRNA gene sequences from samples GS10_PC15_940 (contig-124_471961 and contig-124_482067) and GS10_PC15_1000 (contig-124_27583) were classified as mammalian (Supplementary Table 4). These sequences most likely represent human contamination introduced during sampling, DNA extraction or during sequencing.

In the GS10_PC15_1060 sample we identified an 18S rRNA gene sequence that likely derives from a flatworm (order Rhabdocoela, contig-124_364989). *Chlamydiae* bacterium

K1060_chlam_2, which corresponds to the only Simkaniaceae-like MAG derived from these marine sediments, was also obtained from this sample. Since several of the previously described Simkaniaceae are known symbionts of marine worms[82,83], it is possible that *Chlamydiae* bacterium K1060_chlam_2 might be a symbiont of the Rhabdocoela-related flatworm observed in this sediment layer.

In the GS10_ PC15_940 sample we uncovered 18S rRNA gene sequences that likely derive from an ichthyosporean (contig-124_299197 and contig-124_207553), a green algae (*Micromonas* contig-124_372972) and a chloroplast genome (contig-124_152295). This sample was shown to also contain the actively replicating (Fig. 4a) and highly abundant *Chlamydiae* bacterium K940_chlam_7 (Supplementary Discussion 7), which is most closely related to the protist-associated environmental chlamydiae[10,11]. This raises the possibility that one of the aforementioned eukaryotes could represent a host for *Chlamydiae* bacterium K940_chlam_7. However, given that *Micromonas* is a phototroph it is unlikely that these cells are active in dark marine sediments. Moreover, there have been no reported cases of chlamydiae capable of infecting Archaeplastida (algae and land plants). Alternatively, it is possible that the observed ichthyosporean might represent the host organism of *Chlamydiae* bacterium K940_chlam_7. Yet, little is known about the ecology of ichthyosporeans in marine sediments, and, to our knowledge, there have been no reported cases of chlamydiae capable of infecting ichthyosporeans so far.

Eukaryotes present at low abundances in the samples could have been missed by our sequencing efforts. However, in general, the eukaryotic sequences identified in the samples appear insufficient to account for overall patterns in chlamydial diversity and abundance across all samples. No eukaryotic sequences were identified in sample GS08_GC12_126, where Anoxychlamydiales lineages were found to be exceptionally abundant (Supplementary

Discussion 7). Thereby suggesting that these particular chlamydial lineages may not depend on a eukaryotic host.

**7. Abundance and diversity of chlamydial lineages in Loki's Castle marine sediments**

In a previous study of Loki's Castle marine sediments[108], we detected the presence of Chlamydiae. We further investigated the relative abundance and diversity of Chlamydiae in these sediments using amplicon sequencing of samples taken from four different sediment cores at various depths (Supplementary Table 1, Supplementary Data 2). All of the samples with high chlamydial abundances were isolated from sediment depths found below (but within 1.2 m of) the oxic/anoxic transition zone, which is found at various depths below the seafloor in sediment cores GS08_GC12(0.38 mbsf)[74], GS10_PC15 (1.0 mbsf)[75], and GS10_GC14[75] (0.4 mbsf). The highest diversity of chlamydial OTUs (over 0.1% relative abundance) were observed in anoxic sediment layers (Fig. 1b). When considering individual OTUs found across the marine sediment amplicons, 30 were found to be present in at least five samples (Supplementary Data 2), indicating that a large fraction of the observed chlamydial lineages are commonly found in this environment.

***7.1 Anoxychlamydiales lineages are abundant microbial community members in Loki's Castle marine sediments***

We were unable to link 16S rRNA gene fragments to most of the Anoxychlamydiales MAGs reconstructed in this study (a problem often encountered in genome-resolved metagenomic studies[112,113]). However, in the phylogenetic analysis of the obtained 16S rRNA amplicon sequences (Supplementary Data 4), we identified 17 OTUs that formed a highly supported clade (ufBV = 98) with Anoxychlamydiales member *Chlamydiae* bacterium SM23_39. The OTU abundance of this group mirrors the presence of Anoxychlamydiales bins in the metagenomes from the same samples, indicating that these OTUs represent Anoxychlamydiales 16S rRNA gene sequences. Two of these OTUs (OTU_5_19291 and OTU_255_442) were highly abundant and widespread across sediment samples. OTU_5_19291 is found in 18 samples, with highest relative abundance in all four sediment cores past the oxic/anoxic transition zone. The relative abundance of OTU_5_19291 is above 1% in 7 samples. In one exceptional case it was the most abundant OTU in the GS08_GC12_126 sample, representing 40% of bacterial relative abundance. OTU_255_442, like OTU_5_19291 was most abundant (1.3%) in sample GS08_GC12_126.

##### *7.2 Environmental chlamydiae lineages are abundant in sample GS10_PC15_940*

In our amplicon survey of GS10_PC15_K940, we identified an abundant OTU (OTU_64_1912; 4.8% abundance), which likely corresponding to the *Chlamydiae* bacterium K940_chlam_7 MAG, as they both affiliate with the *Waddliaceae* family in phylogenetic analyses (Fig. 1b, Fig. 2, Supplementary Fig. 3, Supplementary Data 2). *Waddlia chondrophila* is a known animal pathogen[12,84], and *Waddliaceae* family members have been identified both in animal-associated and environmental samples[85]. The wide distribution of these organisms in diverse environments suggests these species could naturally infect protists like other environmental chlamydiae[10,11]. If so, this raises the possibility that the *Waddliaceae*-related *Chlamydiae* bacterium K940_chlam_7 (Supplementary Discussion 6) might be a symbiont of the eukaryotes detected in sample GS10_PC15_K940.

#### 8. Underestimation of environmental abundance and diversity of Chlamydiae

##### *8.1 The environmental distribution of Chlamydiae*

To assess if the high environmental relative abundance and diversity of Chlamydiae identified here (Supplementary Table 1, Supplementary Discussion 7) is unique to Loki's Castle marine sediments, we surveyed chlamydial abundance and diversity in other environments using the Integrated Microbial NGS (IMNGS) platform[86]. IMNGS allows for large-scale taxonomic analysis of 16S rRNA gene amplicon datasets deposited in the Sequence Read Archive (SRA). Using this platform, we identified 13 environments that were enriched for chlamydial diversity (>50 OTUs) and/or abundance (>0.1% relative abundance; Fig. 4b, Supplementary Data 3).

A large proportion of rhizosphere samples (831, corresponding to 62% of samples) and soil samples (2295, corresponding to 14% of samples) harbour relative abundances of Chlamydiae above 0.1%, though comparatively fewer had a high taxonomic richness as based on OTU numbers. Several salt marsh samples were found to contain large numbers of chlamydial OTUs, indicating that this environment represents an unexplored reservoir for uncultured Chlamydiae diversity. Approximately 16% of groundwater samples also appear to harbor a large relative chlamydial abundance, which is congruent with a recent study in which 17 MAGs were assembled from groundwater that resolved five distinct chlamydial lineages[8]. Other environments, including wastewater, activated sludge and bioreactor samples, also contain chlamydial relative abundances above 0.1% of the total microbial community. Interestingly, previous studies have retrieved several chlamydial MAGs affiliated with both CC-II and environmental chlamydiae from such environments (Supplementary Discussion 1, Supplementary Table 3)[87-89]. In addition, some biofilm samples were found to harbour higher (>0,1% relative abundance) chlamydial abundances (15% of samples) and could be an additional environment of interest for studying uncultured chlamydial lineages.

Furthermore, we found that 24% and 14% of freshwater samples contained relative abundances above 0,1% and included more than 50 OTUs, respectively. Samples from freshwater sediments generally also contain high relative abundances of Chlamydiae, but do not necessarily harbour high chlamydial diversity, which is similar to observations made for seawater and marine sediment samples (Fig. 4b, Supplementary Data 3). These findings are in line with a study that revealed a broad taxonomic and phylogenetic diversity of chlamydiae in various environments, particularly from plant, soil and freshwater environments[90]. Altogether, our analyses underline that several environments harbor high diversity and relative abundances of uncultured Chlamydiae, even though primer sets used in environmental surveys were not optimal for detection of chlamydiae (see 8.2).

##### *8.2 Underestimation of chlamydial diversity and abundance in environmental surveys*

Schulz et al.[90] recently reported that taxonomic diversity estimates differ significantly between amplicon and metagenomic surveys. In particular, they observed that taxonomic richness and diversity of Chlamydiae was more pronounced in metagenomic data when compared to amplicon studies. This may be the result of the common use of primers, which do not amplify a large fraction of representatives of the Chlamydiae phylum[91]. For example, the widely used 16S rRNA gene primer sets 515FB and 806RB from the Earth Microbiome Project[92], only capture 0.7% of the characterized chlamydial diversity without any mismatches (though they do capture 95% if allowing a single mismatch (Supplementary Table 5)). Similarly, the universal primer pair A519F/Uni1391R captures less than 1% of chlamydial diversity (Supplementary Table 5). In the present study we therefore used a bacterial-specific primer pair (S-D-0564-a-S-15/SD-Bact-1061-a-A-17) that is predicted to capture ~94% of the presently known chlamydial diversity without mismatches (Supplementary Table 5). Indeed, when comparing the relative abundances of OTUs generated by 16S rRNA amplicon sequencing using the S-D-0564-a-S-15/SD-Bact-1061-a-A-17 and A519F/U1391R primer pairs on sediment core GS08_GC12[93], we found that chlamydial OTUs represented 43% relative abundance using the former primer pair, and less than 1% relative abundance when using the latter. Similarly, while no chlamydial sequences were detected previously in GS10_GC14_75 using the A519F/U1391R primer pair[79], a similar analysis with the S-D-0564-a-S-15/SD-Bact-1061-a-A-17 primer pair recovered 8.9% relative chlamydial abundance.

*8.3 Using culture-independent methods to explore chlamydial genomic diversity*

As evidenced by the present study, culture-independent methods have great potential for expanding genomic representation within the Chlamydiae phylum[94]. The majority of the so far characterized chlamydiae have been isolated by means of co-cultivation (Supplementary Table 3), thus selecting for representatives that can replicate in the respective eukaryotic host. However, most newly identified chlamydial lineages are represented by genome data only which is derived from cultivation-independent studies (Supplementary Table 3). Chlamydiae-targeted studies using cultivation-independent approaches have resulted in the first chlamydial SAGs[36] and the first chlamydial MAGs from metagenomic-based projects targeting animal host-associated populations[2,4]. A number of chlamydial MAGs have also been recently retrieved from whole microbial community metagenomic sequencing efforts of diverse environments, including drinking water treatment plants[88,89], a bioreactor[87], aquifer groundwater[8], a cold-water geyser[95], oceanic waters[37] and river estuary sediment[9]. Altogether, the Chlamydiae phylum is severely understudied at the genomic level, and the future exploration of the microbial communities in additional environments in which Chlamydiae are represented (e.g, see 8.1) will likely yield genomic data from diverse and abundant chlamydial lineages.

#### Supplementary Figures

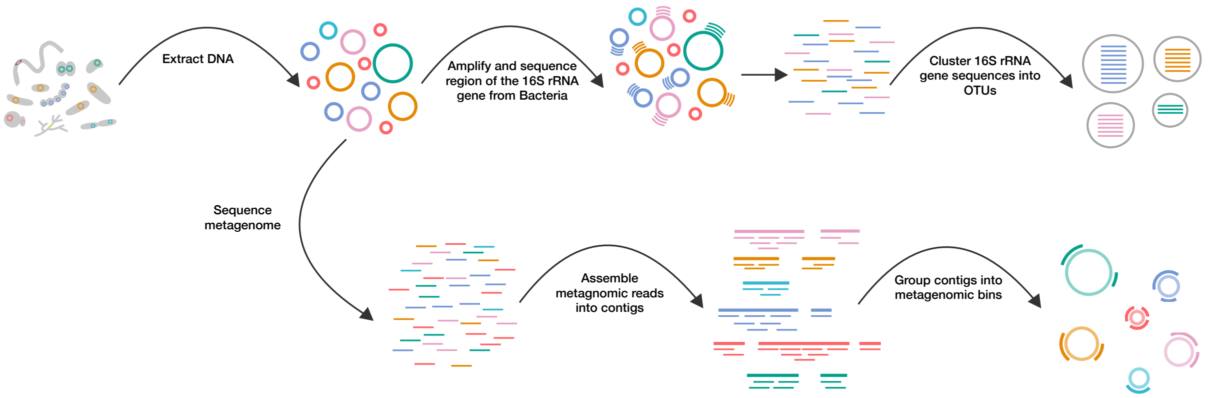

**Supplementary Figure 1.** *Overview of sequencing methods.* For amplicon sequencing, DNA was extracted from 69 marine sediment samples taken near Loki's Castle hydrothermal vent field. These were used as a template for bacterial-specific amplification of an approximately 500 bp region of the 16S rRNA gene and sequenced on an Illumina MiSeq sequencer. These sequences were clustered at the 97% level to generate operational taxonomic units (OTUs). For metagenomic sequencing, DNA was extracted from 4 samples with a high abundance and diversity of chlamydiae. Sequence libraries were prepared and sequenced with Illumina HiSeq and resulting reads of each metagenome were assembled into contigs using IDBA-UD. A differential coverage genome binning approach (using CONCOCT), followed by manual curation, was used to obtain metagenome assembled genomes (MAGs).

**a**

| Sample ID | Metagenome ID | Gbp Sequenced | Gbp Assembled (≥ 1 kb) |
|---|---|---|---|
| GS08_GC12_126 | KR126 | 16.4 | 1.10 |
| GS10_PC15_940 | K940 | 63 | 1.30 |
| GS10_PC15_1000 | K1000 | 53.8 | 1.07 |
| GS10_PC15_1060 | K1060 | 116.4 | 2.38 |

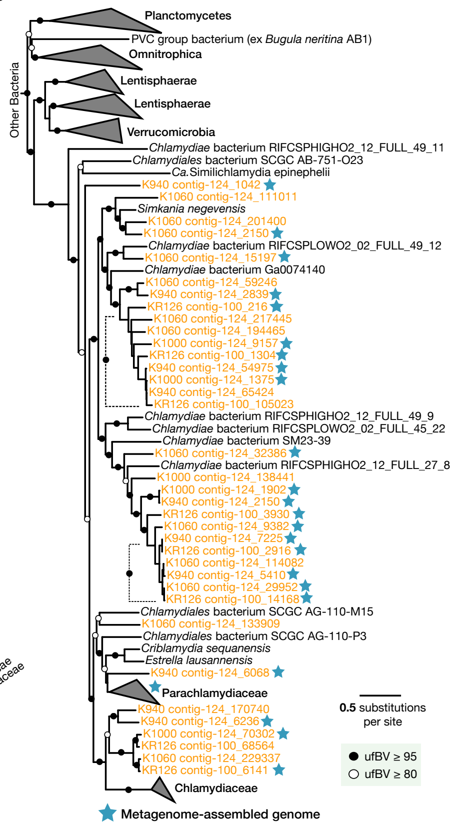

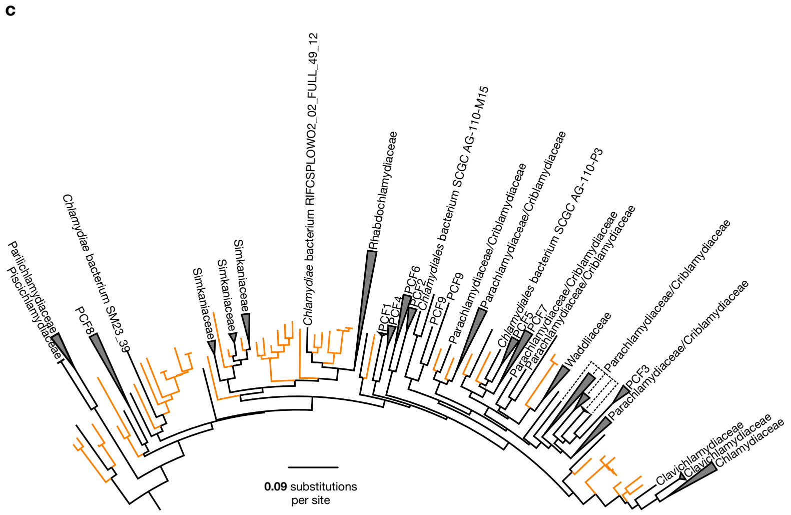

**Supplementary Figure 2.** *Marine sediment metagenome sequencing statistics and chlamydiae diversity*. **a**, Sequencing statistics and sample identifiers for the four sediment samples used for metagenomic sequencing, including the number of Gbp assembled for each metagenome. **b**, Maximum likelihood (ML) tree estimated using an alignment of fifteen ribosomal proteins (at least five of which had to be present) from reference taxa (black, collapsed clades in grey) and marine sediment chlamydiae (orange), under the LG+C60+G model of evolution implemented with IQ-TREE (180 taxa, 2308 sites). Black and white circles represent bipartition values greater than 95 and 80 percent, respectively, from 1000 ultrafast bootstraps (ufBV). Dotted lines indicate the ufBV for all branches in the indicated clade. Sequences corresponding to metagenome-assembled genomes retrieved in this study are indicated with a blue star. **c**, ML phylogeny of chlamydial 16S rRNA gene fragments identified in Loki's Castle metagenomes (orange) in the context of a reference chlamydial dataset (black, collapsed clades in grey), inferred using IQ-TREE with the GTR+R7 model of evolution (344 taxa, 1554 sites).

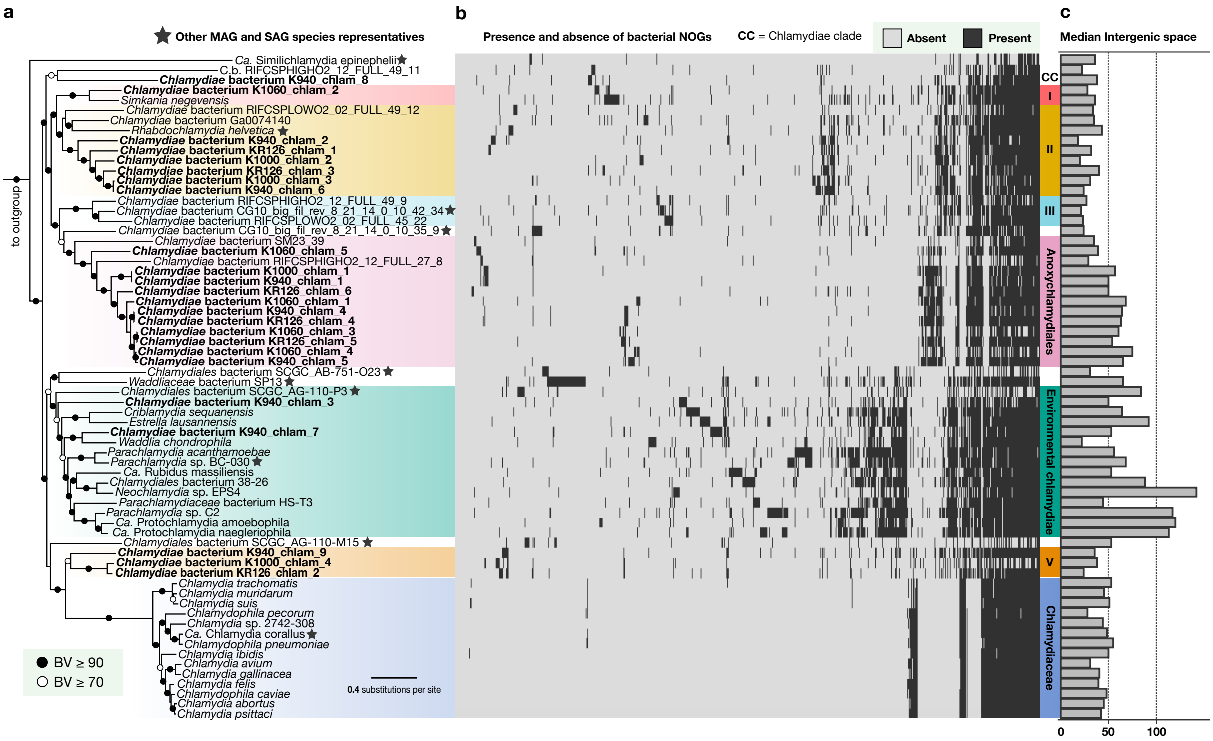

**Supplementary Figure 3.** *Species phylogeny and gene content variation across the Chlamydiae phylum.* **a**, Phylogenetic tree was estimated using a concatenated alignment of 38 single-copy marker proteins, using IQ-TREE under the PMSF approximation of LG+C60 (8072 sites). Bipartitions are labeled with black and white circles representing non-parametric bootstrap values (BV) greater or equal to 90 and 70, respectively. The phylogeny includes other metagenome assembled genome (MAG) and single-cell assembled genomes (SAG) chlamydiae species representatives (stars, see Methods, Supplementary Table 3). **b**, Presence (in dark grey) and absence (in light grey) of NOGs found across all chlamydial lineages, with delineated Chlamydiae clades indicated. **c**, Median intergenic space in bp across chlamydial genomes.

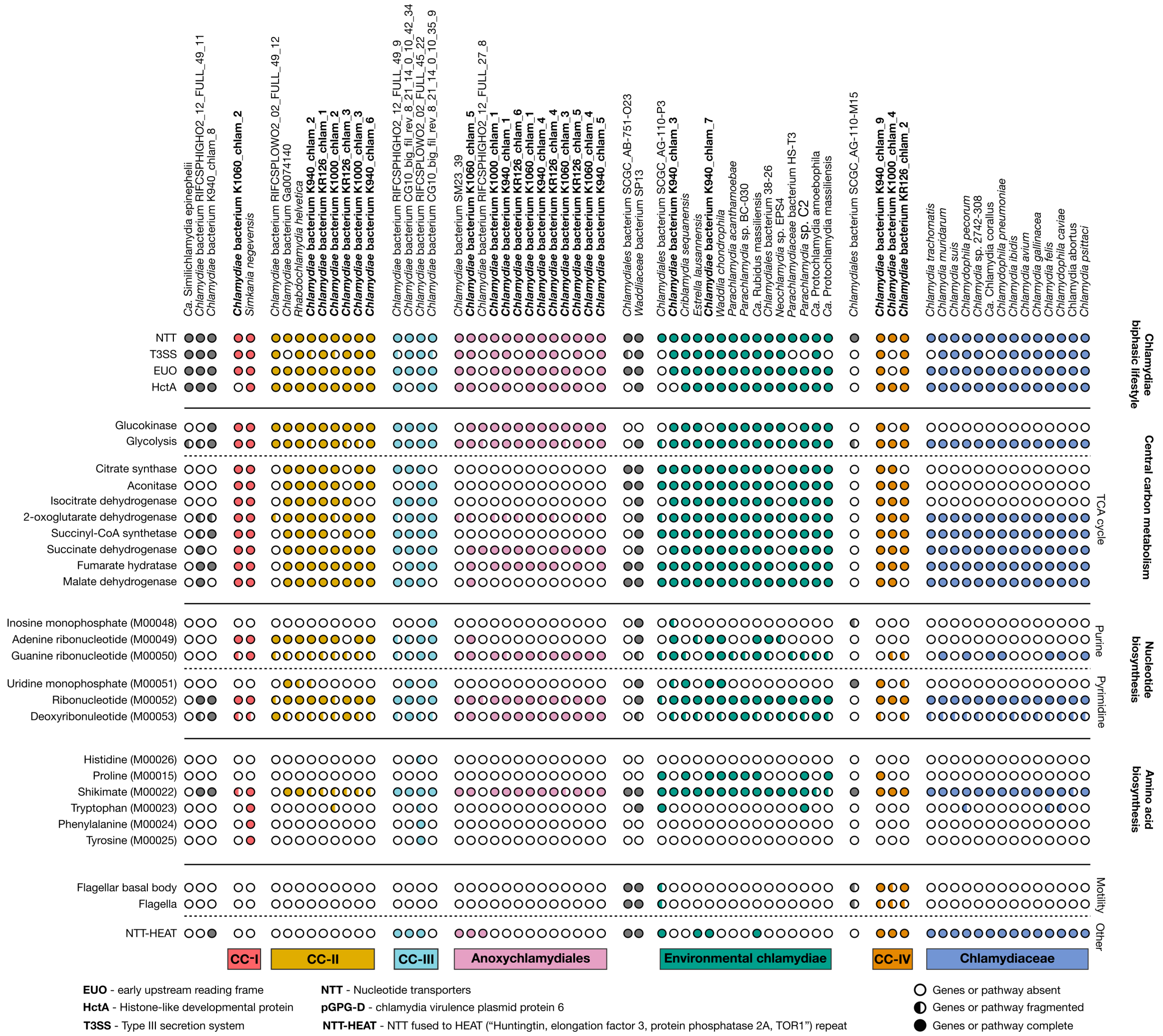

**Supplementary Figure 4.** *Overview of selected protein content across* Chlamydiae. Presence of selected proteins and pathways including traits associated with the chlamydiae biphasic lifecycle, components of central carbon metabolism and nucleotide and amino acid biosynthesis, across *Chlamydiae* species representatives color-coded according to *Chlamydiae* clades. Where relevant the corresponding KEGG pathway module is indicated in brackets.

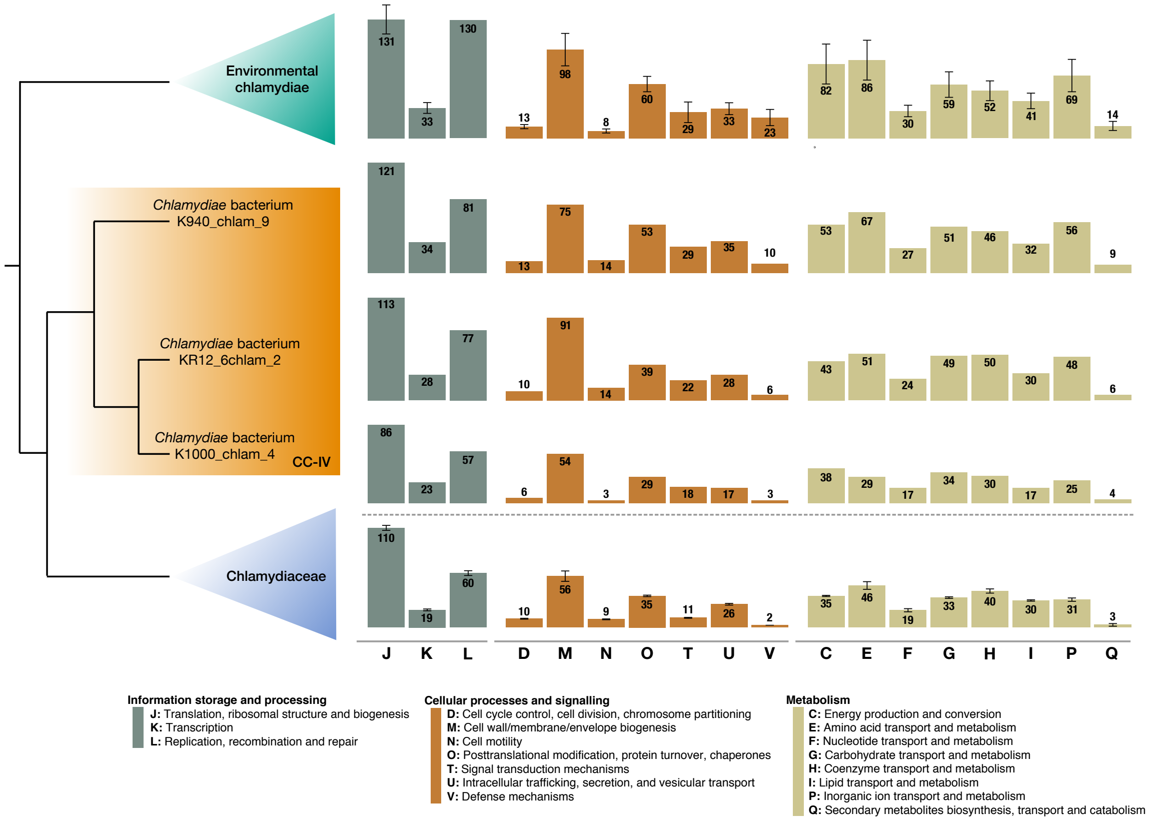

**Supplementary Figure 5.** *COG category distribution patterns*. Distributions of the number of NOGs assigned across COG categories for environmental chlamydiae (mean and standard deviation), CC-IV, and Chlamydiaceae (mean and standard deviation).

NOG or PF Present (dark grey) / NOG or PF Absent (light grey)

| NOG or PF Domain | Description | *Chlamydia trachomatis* | *Chlamydia muridarum* | *Chlamydia suis* | *Chlamydophila pecorum* | *Chlamydia* sp. 2742-308 | *Ca.* Chlamydia corallus | *Chlamydophila pneumoniae* | *Chlamydia ibidis* | *Chlamydia avium* | *Chlamydia gallinacea* | *Chlamydia felis* | *Chlamydophila caviae* | *Chlamydia abortus* | *Chlamydia psittaci* |
|---|---|---|---|---|---|---|---|---|---|---|---|---|---|---|---|
| **Host Interaction and Adhesion** | | | | | | | | | | | | | | | |
| 0ZM85 | Polymorphic membrane protein - family A | 1 | 1 | 1 | | 1 | 1 | 1 | 1 | 1 | 1 | 1 | 1 | 1 | 1 |
| 0Y0KC | Polymorphic membrane protein - family B/C | 2 | 2 | 2 | 1 | 1 | 1 | 2 | 1 | 1 | 1 | 1 | 1 | 1 | 1 |
| 0XV92 | Polymorphic outer membrane protein - family D/E/F/G/H | 5 | 5 | 6 | 11 | 14 | 19 | 13 | 22 | 4 | 5 | 21 | 13 | 10 | 13 |
| 0Y3IR | Polymorphic membrane protein - family G | 1 | 1 | 1 | 2 | 1 | 1 | 1 | 1 | 1 | 1 | 1 | 1 | 1 | 1 |
| PF03503 | Chlamydia cysteine-rich outer membrane protein 3 | 1 | 1 | 1 | 1 | 1 | 1 | 1 | 1 | 1 | 1 | 1 | 1 | 1 | 1 |
| PF05745 | Chlamydia 15 kDa cysteine-rich outer membrane protein (CRPA) | 1 | 1 | 1 | 1 | 1 | 1 | 1 | 1 | 1 | 1 | | 1 | 1 | 1 |
| PF04156 | IncA protein | 4 | 5 | 3 | 1 | | 2 | 9 | 1 | 1 | 1 | 8 | 10 | 8 | 10 |
| PF17628 | Inclusion membrane protein D | 1 | 1 | 1 | | 1 | | | | | | | 1 | | |
| **Virulence Factors** | | | | | | | | | | | | | | | |
| 0Y2RT | Porin AaxA/Carbohydrate-selective porin OprB | 1 | 1 | 1 | 1 | 1 | 1 | 1 | 1 | 1 | 1 | 1 | 1 | 1 | 1 |
| COG1945 | Arginine decarboxylase | 1 | 1 | 1 | 1 | 1 | 1 | 1 | 1 | 1 | 1 | 1 | 1 | 1 | 1 |
| 0Y3HQ | Membrane attack complex (MAC) perforin | 1 | 1 | 1 | 1 | | 1 | 1 | 2 | 1 | | 2 | | 1 | 2 |
| 0ZUEX | Adherence factor/cytotoxin | 4 | 3 | 2 | 2 | | | | 2 | | 1 | 1 | 1 | | 1 |
| PF05475 | Pgp3 C-terminal domain | | 1 | 1 | | 1 | 1 | | | | 1 | 1 | 1 | | 1 |
| **Vitamin Biosynthesis (Folate)** | | | | | | | | | | | | | | | |
| COG1478 | Alternate folylglutamate synthase FolC2 | 1 | 1 | 1 | | 1 | 1 | 1 | 1 | 1 | 1 | 1 | 1 | 1 | 1 |
| 0ZGA4 | Dihydroneopterin aldolase FolB | 1 | 1 | 1 | | 1 | 1 | 1 | 1 | 1 | 1 | 1 | 1 | | 1 |
| **Metabolism** | | | | | | | | | | | | | | | |
| COG1218 | 3'(2'),5'bisphosphate nucleotidase | 1 | 1 | 1 | 1 | 1 | 1 | 1 | 1 | 1 | 1 | 1 | 1 | 1 | 1 |
| 0Z2QW | Adenosine AMP deaminase | | 1 | | 1 | | 1 | 1 | | | | 1 | 1 | | 1 |
| COG0352 | Thiamine monophosphate synthase | | | | | | | | 1 | | | 1 | 1 | 1 | 1 |
| **Gene Expression** | | | | | | | | | | | | | | | |
| PF07382 | Histone H1-like nucleoprotein HC2 | 1 | 1 | 2 | 1 | 1 | 1 | 1 | 1 | 1 | 1 | 1 | 1 | 1 | 1 |
| PF17455 | Late transcription unit B protein | 1 | 1 | 1 | 1 | 1 | 1 | 1 | 1 | 1 | 1 | 1 | 1 | 1 | 1 |
| PF17446 | Late transcription unit A protein | 1 | 1 | 1 | | 1 | | 1 | 1 | 1 | | 1 | 1 | 1 | 1 |
| **Unknown Function** | | | | | | | | | | | | | | | |
| 11VHY | Conserved hypothetical protein | 1 | 1 | 1 | 1 | 1 | 1 | 1 | 1 | 1 | 1 | 1 | 1 | 1 | 1 |
| 0Y23D | DUF5398 - Domain of unknown function | 1 | 1 | 1 | 1 | 1 | 1 | 1 | 1 | 1 | 1 | 1 | 1 | 1 | 1 |
| PF06587 | DUF1137 - Domain of unknown function | 1 | 1 | 1 | 1 | 1 | 1 | 1 | 1 | 1 | 1 | 1 | 1 | 1 | 1 |
| PF16802 | DUF5070 - Domain of unknown function | 1 | 1 | 1 | 1 | 1 | 1 | 1 | 1 | 1 | 1 | 1 | 1 | 1 | 1 |
| PF07577 | DUF1547 - Domain of unknown function | 1 | 1 | 1 | 1 | 1 | | | 1 | | 1 | 1 | 1 | 1 | 1 |
| PF07146 | DUF1389 - Domain of unknown function | | | | 2 | 3 | 4 | 5 | 4 | 3 | 3 | 3 | 7 | 3 | 7 |
| PF07560; PF07579 | DUF1539 and DUF1548 - Domains of unknown function | | | | 2 | 1 | 1 | 1 | 1 | 1 | 1 | 3 | 3 | 3 | 3 |

**Supplementary Figure 6.** *Conserved gene content restricted to the Chlamydiaceae family.* Presence (dark grey), absence (light grey), and number of genes assigned to NOGs or with PF domains found uniquely within Chlamydiaceae lineages among Chlamydiae, and which is conserved across the family (in a third of representative genomes).

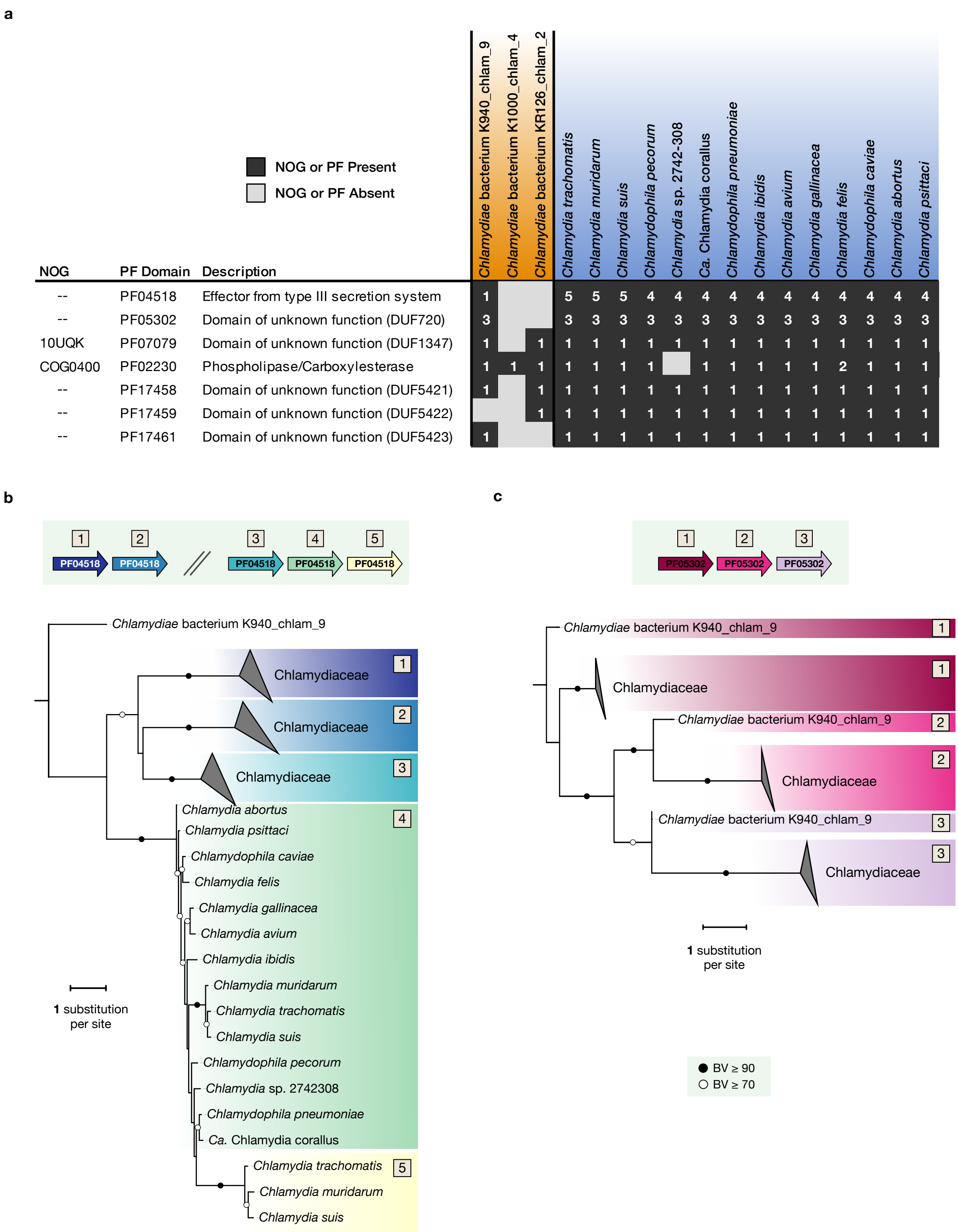

**Supplementary Figure 7.** *Evolutionary insights into gene content shared between Chlamydiaceae and CC-IV.* **a**, Presence (dark grey), absence (light grey), and number of genes assigned to NOGs or with PF domains found conserved uniquely in CC-IV and Chlamydiaceae lineages among Chlamydiae. Phylogenetic tree and typical genomic organization of gene families containing PF domains **b**, PF04518 and **c**, PF05302. Phylogenies were inferred with IQ-TREE under the PMSF approximation of LG+C20+G+F (PF04518: 395 sites, PF05302: 126 sites). Bipartitions are labeled with black and white circles representing non-parametric bootstrap values (BV) greater or equal to 90 and 70, respectively.

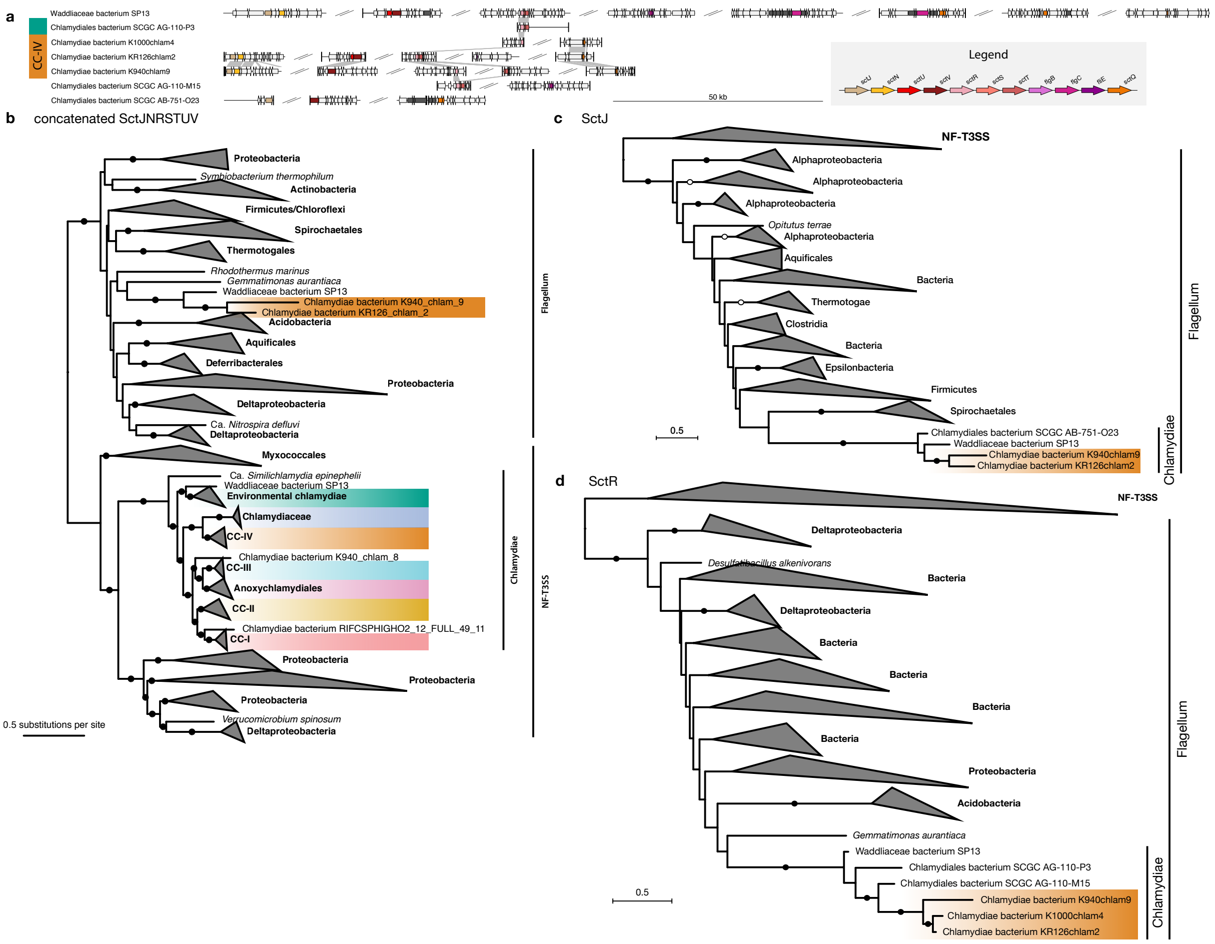

**Supplementary Figure 8.** *Synteny of flagellar components found in chlamydial lineages and phylogenetic analyses of homologous NF-T3SS and flagellar components*. **a**, Synteny of flagellar genes in *Chlamydiae*, following the gene nomenclature by Abby et al.[713030]. All genomes with at least one homolog of flagellar genes detected by MacSyFinder are included, except for those genomes with flagellear homologs of *sctN* and *sctV*, which have been reported to be co-opted by the NF-T3SS machinery (see Supplementary Discussion). Genes (arrows) are colored according to the legend next to the synteny plot. Genome regions are defined as 10 kb up- and downstream the colored genes, and are truncated at contig boundaries (thicker, vertical lines). Comparison lines between genes represent best reciprocal BLASTP hits with an e-value less than or equal to 0.001. Phylogenies of **b**, a concatenated dataset of the SctJNRSTUV proteins (PMSF approximation of LG+F+C50+R4, 626 sequences, 1635 sites), **c**, the SctJ protein (LG+F+C40+R4, 630 sequences, 126 sites) and **d**, the SctR protein (LG+F+C40+R4, 651 sequences, 171 sites). All phylogenies were reconstructed with IQ-TREE and were rooted with the respective paralogues. Bipartitions are labeled with black and white circles representing non-parametric bootstrap values (BV) greater or equal to 90 and 70, respectively.

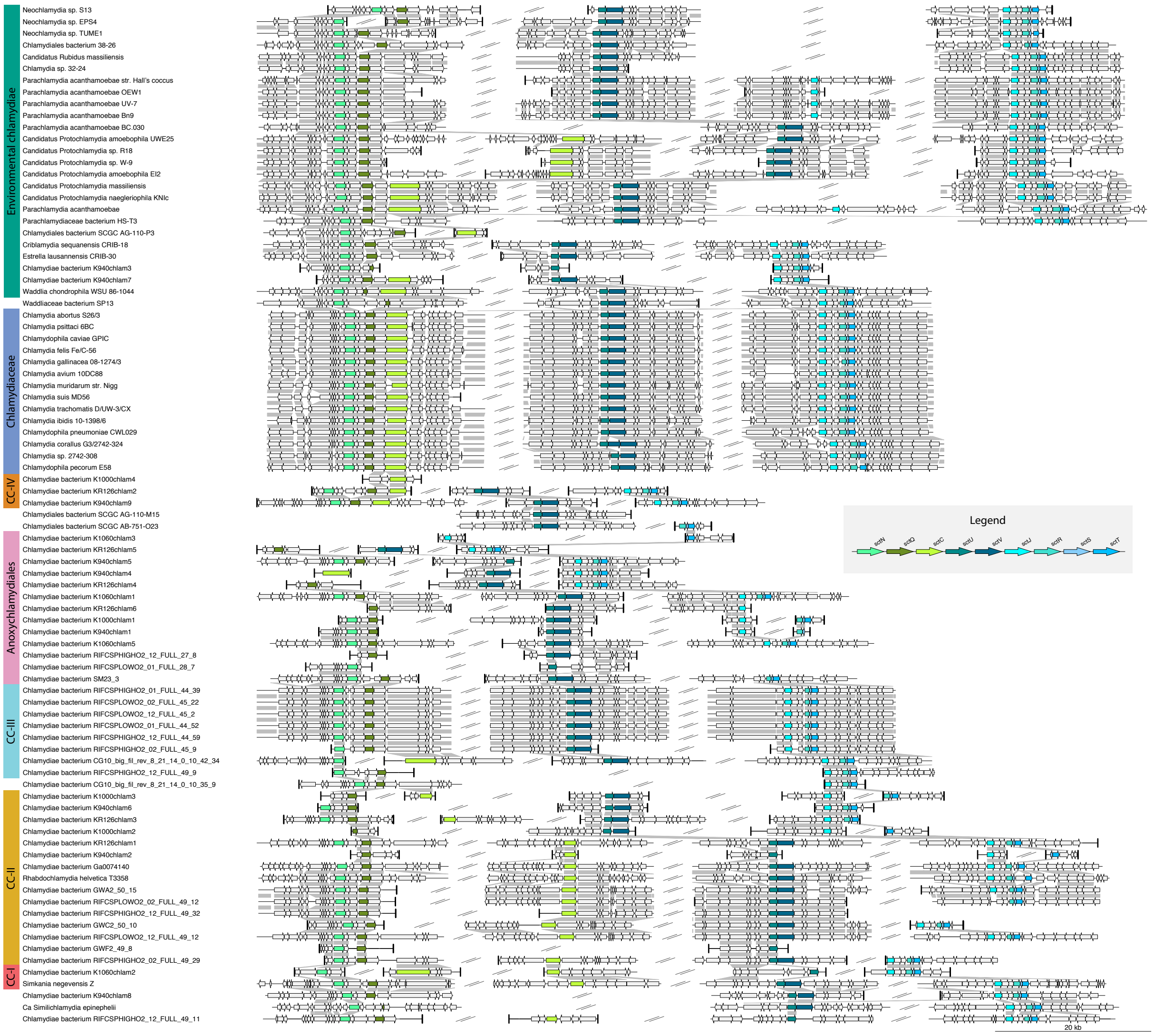

**Supplementary Figure 9**. *Conserved synteny of NF-T3SS components across Chlamydiae.* Synteny plot including genomes with at least one homolog of NF-T3SS genes detected by MacSyFinder. Genes (arrows) are colored according to the legend. Genome regions are defined as 10 kb up- and downstream of the colored genes, and are truncated at contig boundaries (thicker, vertical lines). Comparison lines between genes represent best reciprocal BLASTP hits with an e-value less than or equal to 0.001.

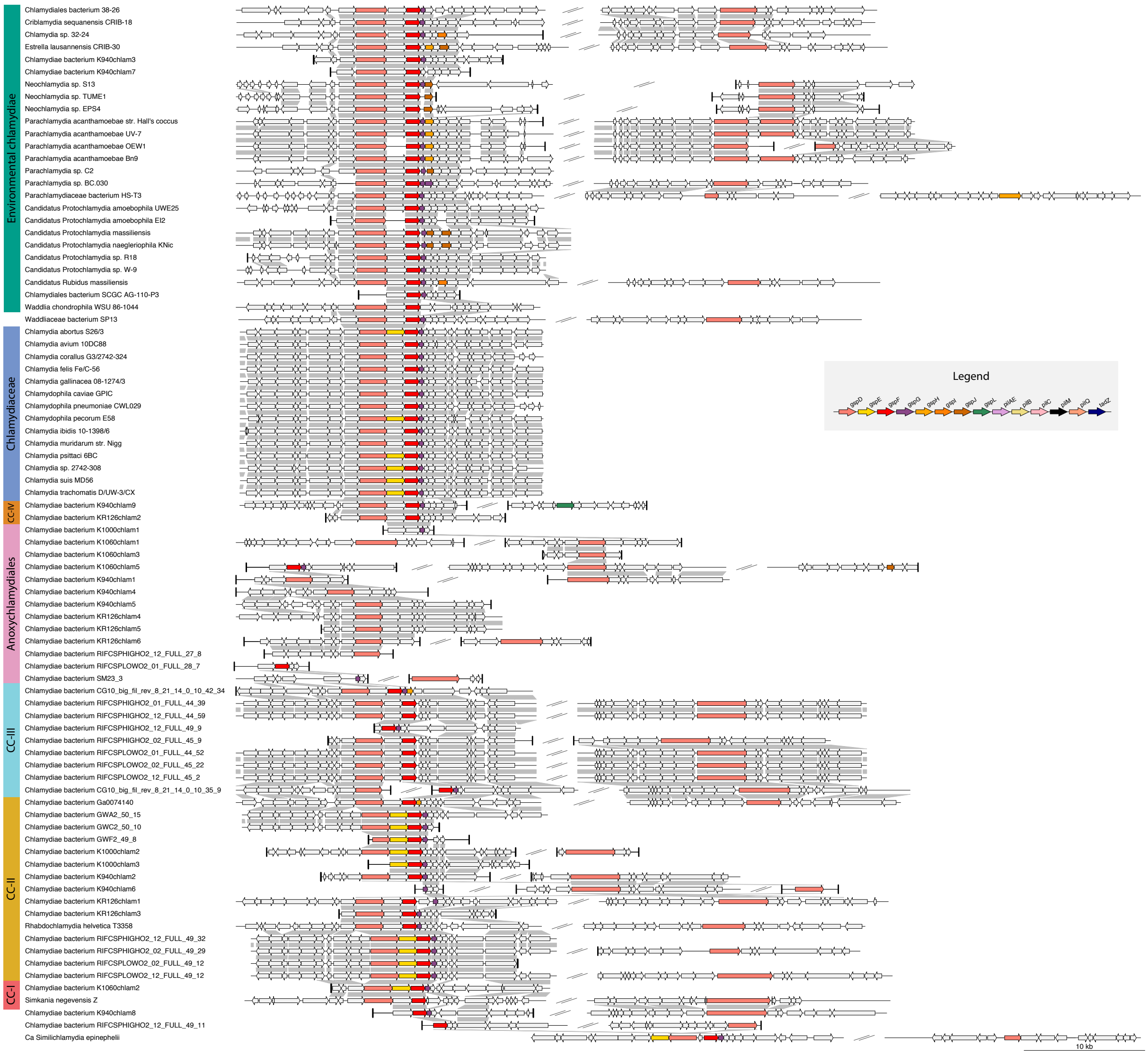

**Supplementary Figure 10.** *Conserved synteny of T2SS components across Chlamydiae.* Synteny plot including Chlamydiae genomes with at least one T2SS detected by MacSyFinder (Supplementary Data 3). Genes (arrows) are colored according to the legend. Genome regions are defined as 10 kb up- and downstream of the colored genes, and are truncated at contig boundaries (thicker, vertical lines). Comparison lines between genes represent best reciprocal BLASTP hits with an e-value less than or equal to 0.001.

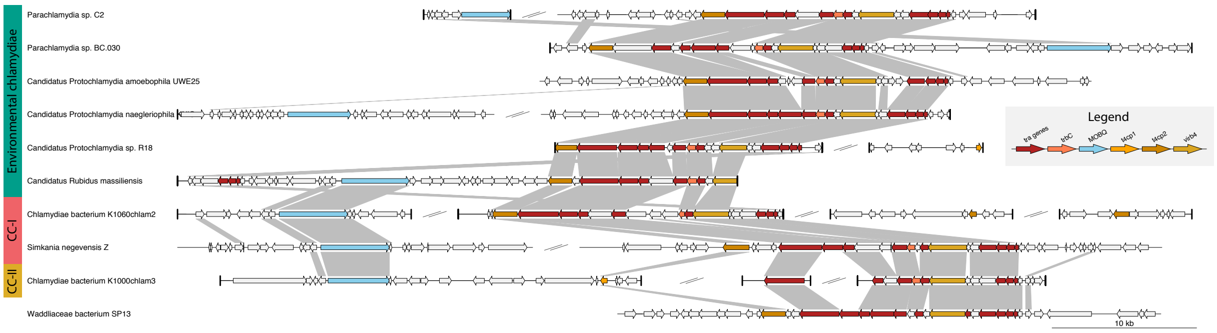

**Supplementary Figure 11**. *Conserved synteny of T4SS components in Chlamydiae.* Synteny plot including Chlamydiae genomes with at least one T4SS detected by MacSyFinder (Supplementary Data 3). Genes (arrows) are colored according to the legend. Genome regions are defined as 10 kb up- and downstream of the colored genes, and are truncated at contig boundaries (thicker, vertical lines). Comparison lines between genes represent best reciprocal BLASTP hits with an e-value less than or equal to 0.001.

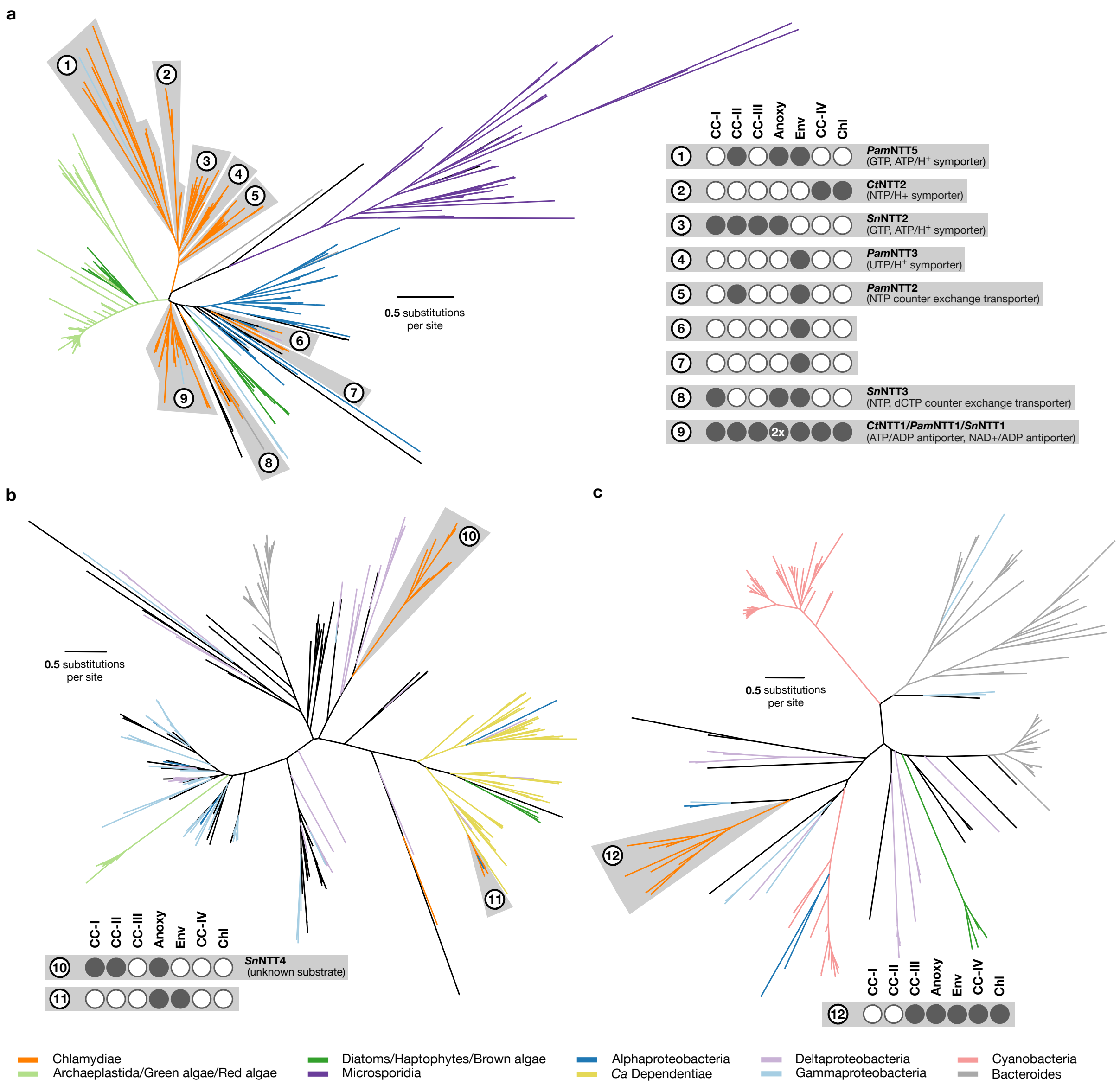

**Supplementary Figure 12**. *Phylogenetic inference of nucleotide transporters.* Phylogenetic trees of nucleotide transporter (NTT) proteins found in both prokaryotes and eukaryotes. Chlamydiae are shown in orange with clade affiliation of the chlamydiae within each chlamydial NTT cluster indicated by the coloured circles. Functionally characterized NTTs from each cluster are indicated. Species and clade name abbreviations: environmental chlamydiae (Env), Chlamydiaceae (Chl), Anoxychlamydiales (Anoxy), *Chlamydia trachomatis* (Ct), *Ca.* Protochlamydia acanthamoebae (Pam), *Simkania negevensis* (Sn). See legend for color scheme of additional lineages. **a**, ML phylogeny inferred using IQ-TREE with the LG+F+R8 model of "canonical NTTs" (400 taxa, 357 sites) **b**, ML phylogeny of "other NTTs" (that form a sister clade to the "canonical NTTs"), inferred using IQ-TREE with the LG+F+R8 model of evolution (302 taxa, 348 sites). **c**, ML phylogeny of "NTT-HEAT NTTs", inferred using IQ-TREE with the LG+F+R6 model of evolution (157 taxa, 329 sites).

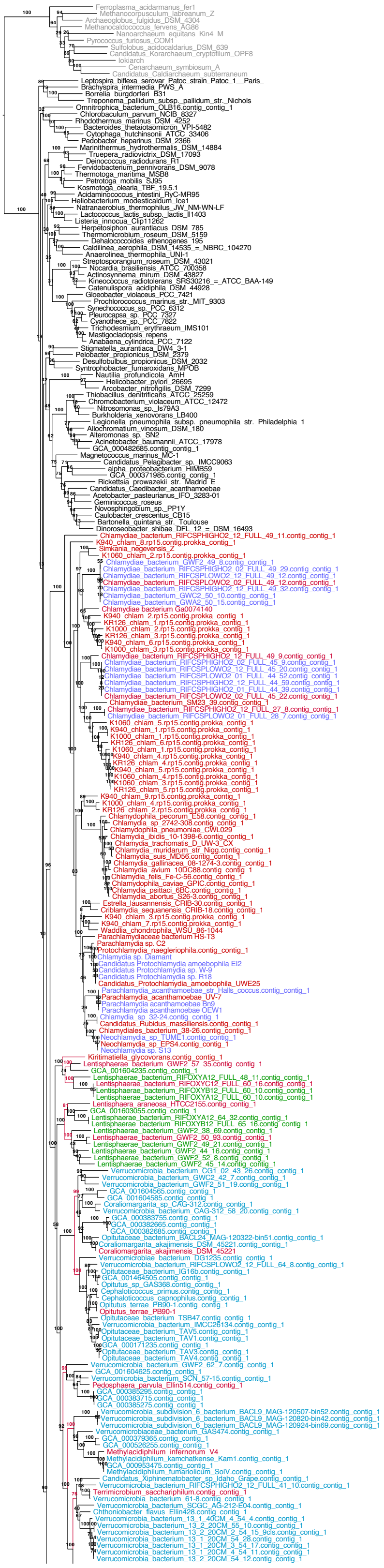
Ferroplasma_acidarmanus_fer1
Methanocorpusculum_labreanum_Z
Archaeoglobus_fulgidus_DSM_4304
Methanocaldococcus_fervens_AG86
Nanoarchaeum_equitans_Kin4_M
Pyrococcus_furiosus_COM1
Sulfolobus_acidocaldarius_DSM_639
Candidatus_Korarchaeum_cryptofilum_OPF8
lokiarch
Cenarchaeum_symbiosum_A
Candidatus_Caldiarchaeum_subterraneum
Leptospira_biflexa_serovar_Patoc_strain_Patoc_1__Paris_
Brachyspira_intermedia_PWS_A
Borrelia_burgdorferi_B31
Treponema_pallidum_subsp._pallidum_str._Nichols
Omnitrophica_bacterium_OLB16.contig_contig_1
Chlorobaculum_parvum_NCIB_8327
Rhodothermus_marinus_DSM_4252
Bacteroides_thetaiotaomicron_VPI-5482
Cytophaga_hutchinsonii_ATCC_33406
Pedobacter_heparinus_DSM_2366
Marinithermus_hydrothermalis_DSM_14884
Truepera_radiovictrix_DSM_17093
Deinococcus_radiodurans_R1
Fervidobacterium_pennivorans_DSM_9078
Thermotoga_maritima_MSB8
Petrotoga_mobilis_SJ95
Kosmotoga_olearia_TBF_19.5.1
Acidaminococcus_intestini_RyC-MR95
Heliobacterium_modesticaldum_Ice1
Natranaerobius_thermophilus_JW_NM-WN-LF
Lactococcus_lactis_subsp._lactis_Il1403
Listeria_innocua_Clip11262
Herpetosiphon_aurantiacus_DSM_785
Thermomicrobium_roseum_DSM_5159
Dehalococcoides_ethenogenes_195
Caldilinea_aerophila_DSM_14535_=_NBRC_104270
Anaerolinea_thermophila_UNI-1
Streptosporangium_roseum_DSM_43021
Nocardia_brasiliensis_ATCC_700358
Actinosynnema_mirum_DSM_43827
Kineococcus_radiotolerans_SRS30216_=_ATCC_BAA-149
Catenulispora_acidiphila_DSM_44928
Gloeobacter_violaceus_PCC_7421
Prochlorococcus_marinus_str._MIT_9303
Synechococcus_sp._PCC_6312
Pleurocapsa_sp._PCC_7327
Cyanothece_sp._PCC_7822
Trichodesmium_erythraeum_IMS101
Mastigocladopsis_repens
Anabaena_cylindrica_PCC_7122
Stigmatella_aurantiaca_DW4_3-1
Pelobacter_propionicus_DSM_2379
Desulfobulbus_propionicus_DSM_2032
Syntrophobacter_fumaroxidans_MPOB
Nautilia_profundicola_AmH
Helicobacter_pylori_26695
Arcobacter_nitrofigilis_DSM_7299
Thiobacillus_denitrificans_ATCC_25259
Chromobacterium_violaceum_ATCC_12472
Nitrosomonas_sp._Is79A3
Burkholderia_xenovorans_LB400
Legionella_pneumophila_subsp._pneumophila_str._Philadelphia_1
Allochromatium_vinosum_DSM_180
Alteromonas_sp._SN2
Acinetobacter_baumannii_ATCC_17978
GCA_000482685.contig_contig_1
Magnetococcus_marinus_MC-1
Candidatus_Pelagibacter_sp._IMCC9063
alpha_proteobacterium_HIMB59
GCA_000371985.contig_contig_1
Rickettsia_prowazekii_str._Madrid_E
Candidatus_Caedibacter_acanthamoebae
Acetobacter_pasteurianus_IFO_3283-01
Geminicoccus_roseus
Novosphingobium_sp._PP1Y
Caulobacter_crescentus_CB15
Bartonella_quintana_str._Toulouse
Dinoroseobacter_shibae_DFL_12_=_DSM_16493
Chlamydiae_bacterium_RIFCSPHIGHO2_12_FULL_49_11.contig_contig_1
K940_chlam_8.rp15.contig.prokka_contig_1
Simkania_negevensis_Z
K1060_chlam_2.rp15.contig.prokka_contig_1
Chlamydiae_bacterium_GWF2_49_8.contig_contig_1
Chlamydiae_bacterium_RIFCSPHIGHO2_02_FULL_49_29.contig_contig_1
Chlamydiae_bacterium_RIFCSPLOWO2_12_FULL_49_12.contig_contig_1
Chlamydiae_bacterium_RIFCSPLOWO2_02_FULL_49_12.contig_contig_1
Chlamydiae_bacterium_RIFCSPHIGHO2_12_FULL_49_32.contig_contig_1
Chlamydiae_bacterium_GWC2_50_10.contig_contig_1
Chlamydiae_bacterium_GWA2_50_15.contig_contig_1
Chlamydiae bacterium Ga0074140
K940_chlam_2.rp15.contig.prokka_contig_1
KR126_chlam_1.rp15.contig.prokka_contig_1
K1000_chlam_2.rp15.contig.prokka_contig_1
KR126_chlam_3.rp15.contig.prokka_contig_1
K940_chlam_6.rp15.contig.prokka_contig_1
K1000_chlam_3.rp15.contig.prokka_contig_1
Chlamydiae_bacterium_RIFCSPHIGHO2_12_FULL_49_9.contig_contig_1
Chlamydiae_bacterium_RIFCSPHIGHO2_02_FULL_45_9.contig_contig_1
Chlamydiae_bacterium_RIFCSPLOWO2_12_FULL_45_20.contig_contig_1
Chlamydiae_bacterium_RIFCSPLOWO2_01_FULL_44_52.contig_contig_1
Chlamydiae_bacterium_RIFCSPHIGHO2_12_FULL_44_59.contig_contig_1
Chlamydiae_bacterium_RIFCSPHIGHO2_01_FULL_44_39.contig_contig_1
Chlamydiae_bacterium_RIFCSPLOWO2_02_FULL_45_22.contig_contig_1
Chlamydiae_bacterium_SM23_39.contig_contig_1
Chlamydiae_bacterium_RIFCSPHIGHO2_12_FULL_27_8.contig_contig_1
Chlamydiae_bacterium_RIFCSPLOWO2_01_FULL_28_7.contig_contig_1
K1060_chlam_5.rp15.contig.prokka_contig_1
K940_chlam_1.rp15.contig.prokka_contig_1
K1000_chlam_1.rp15.contig.prokka_contig_1
KR126_chlam_6.rp15.contig.prokka_contig_1
K1060_chlam_1.rp15.contig.prokka_contig_1
K940_chlam_4.rp15.contig.prokka_contig_1
KR126_chlam_4.rp15.contig.prokka_contig_1
K940_chlam_5.rp15.contig.prokka_contig_1
K1060_chlam_4.rp15.contig.prokka_contig_1
K1060_chlam_3.rp15.contig.prokka_contig_1
KR126_chlam_5.rp15.contig.prokka_contig_1
K940_chlam_9.rp15.contig.prokka_contig_1
K1000_chlam_4.rp15.contig.prokka_contig_1
KR126_chlam_2.rp15.contig.prokka_contig_1
Chlamydophila_pecorum_E58.contig_contig_1
Chlamydia_sp_2742-308.contig_contig_1
Chlamydophila_pneumoniae_CWL029
Chlamydia_ibidis_10-1398-6.contig_contig_1
Chlamydia_trachomatis_D_UW-3_CX
Chlamydia_muridarum_str_Nigg.contig_contig_1
Chlamydia_suis_MD56.contig_contig_1
Chlamydia_gallinacea_08-1274-3.contig_contig_1
Chlamydia_avium_10DC88.contig_contig_1
Chlamydia_felis_Fe-C-56.contig_contig_1
Chlamydophila_caviae_GPIC.contig_contig_1
Chlamydia_psittaci_6BC.contig_contig_1
Chlamydia_abortus_S26-3.contig_contig_1
Estrella_lausannensis_CRIB-30.contig_contig_1
Criblamydia_sequanensis_CRIB-18.contig_contig_1
K940_chlam_3.rp15.contig.prokka_contig_1
K940_chlam_7.rp15.contig.prokka_contig_1
Waddlia_chondrophila_WSU_86-1044
Parachlamydiaceae bacterium HS-T3
Parachlamydia sp. C2
Protochlamydia_naegleriophila.contig_contig_1
Chlamydia sp. Diamant
Candidatus Protochlamydia amoebophila EI2
Candidatus Protochlamydia sp. W-9
Candidatus Protochlamydia sp. R18
Candidatus_Protochlamydia_amoebophila_UWE25
Parachlamydia_acanthamoebae_str_Halls_coccus.contig_contig_1
Parachlamydia_acanthamoebae_UV-7
Parachlamydia acanthamoebae Bn9
Parachlamydia acanthamoebae OEW1
Chlamydia_sp_32-24.contig_contig_1
Candidatus_Rubidus_massiliensis.contig_contig_1
Chlamydiales_bacterium_38-26.contig_contig_1
Neochlamydia_sp_TUME1.contig_contig_1
Neochlamydia_sp_EPS4.contig_contig_1
Neochlamydia sp. S13
Kiritimatiella_glycovorans.contig_contig_1
Lentisphaerae_bacterium_GWF2_57_35.contig_contig_1
GCA_001604235.contig_contig_1
Lentisphaerae_bacterium_RIFOXYA12_FULL_48_11.contig_contig_1
Lentisphaerae_bacterium_RIFOXYC12_FULL_60_16.contig_contig_1
Lentisphaerae_bacterium_RIFOXYB12_FULL_60_10.contig_contig_1
Lentisphaerae_bacterium_RIFOXYA12_FULL_60_10.contig_contig_1
Lentisphaera_araneosa_HTCC2155.contig_contig_1
GCA_001603055.contig_contig_1
Lentisphaerae_bacterium_RIFOXYA12_64_32.contig_contig_1
Lentisphaerae_bacterium_RIFOXYB12_FULL_65_16.contig_contig_1
Lentisphaerae_bacterium_GWF2_38_69.contig_contig_1
Lentisphaerae_bacterium_GWF2_50_93.contig_contig_1
Lentisphaerae_bacterium_GWF2_49_21.contig_contig_1
Lentisphaerae_bacterium_GWF2_44_16.contig_contig_1
Lentisphaerae_bacterium_GWF2_52_8.contig_contig_1
Lentisphaerae_bacterium_GWF2_45_14.contig_contig_1
Verrucomicrobia_bacterium_CG1_02_43_26.contig_contig_1
Verrucomicrobia_bacterium_GWC2_42_7.contig_contig_1
Verrucomicrobia_bacterium_GWF2_51_19.contig_contig_1
GCA_001604565.contig_contig_1
GCA_001604585.contig_contig_1
Coraliomargarita_sp_CAG-312.contig_contig_1
Verrucomicrobia_bacterium_CAG-312_58_20.contig_contig_1
GCA_000383755.contig_contig_1
GCA_000382665.contig_contig_1
GCA_000382685.contig_contig_1
Opitutaceae_bacterium_BACL24_MAG-120322-bin51.contig_contig_1
Coraliomargarita_akajimensis_DSM_45221.contig_contig_1
Coraliomargarita_akajimensis_DSM_45221
Verrucomicrobiae_bacterium_DG1235.contig_contig_1
Verrucomicrobia_bacterium_RIFCSPLOWO2_12_FULL_64_8.contig_contig_1
Opitutaceae_bacterium_IG16b.contig_contig_1
GCA_001464505.contig_contig_1
Opitutus_sp_GAS368.contig_contig_1
Cephaloticoccus_primus.contig_contig_1
Cephaloticoccus_capnophilus.contig_contig_1
Opitutus_terrae_PB90-1.contig_contig_1
Opitutus_terrae_PB90-1
Opitutaceae_bacterium_TSB47.contig_contig_1
Verrucomicrobia_bacterium_IMCC26134.contig_contig_1
Opitutaceae_bacterium_TAV5.contig_contig_1
Opitutaceae_bacterium_TAV1.contig_contig_1
GCA_000171235.contig_contig_1
Opitutaceae_bacterium_TAV3.contig_contig_1
Opitutaceae_bacterium_TAV4.contig_contig_1
Verrucomicrobia_bacterium_GWF2_62_7.contig_contig_1
GCA_001604625.contig_contig_1
Verrucomicrobia_bacterium_SCN_57-15.contig_contig_1
Pedosphaera_parvula_Ellin514.contig_contig_1
GCA_000385295.contig_contig_1
GCA_000383715.contig_contig_1
GCA_000385275.contig_contig_1
Verrucomicrobia_subdivision_6_bacterium_BACL9_MAG-120507-bin52.contig_contig_1
Verrucomicrobia_subdivision_6_bacterium_BACL9_MAG-120820-bin42.contig_contig_1
Verrucomicrobia_subdivision_6_bacterium_BACL9_MAG-120924-bin69.contig_contig_1
Verrucomicrobiaceae_bacterium_GAS474.contig_contig_1
GCA_000379365.contig_contig_1
GCA_000526255.contig_contig_1
Methylacidiphilum_infernorum_V4
Methylacidiphilum_kamchatkense_Kam1.contig_contig_1
GCA_000953475.contig_contig_1
Methylacidiphilum_fumariolicum_SolV.contig_contig_1
Candidatus_Xiphinematobacter_sp_Idaho_Grape.contig_contig_1
Verrucomicrobia_bacterium_RIFCSPHIGHO2_12_FULL_41_10.contig_contig_1
Terrimicrobium_sacchariphilum.contig_contig_1
Verrucomicrobia_bacterium_61-8.contig_contig_1
Verrucomicrobia_bacterium_SCGC_AG-212-E04.contig_contig_1
Chthoniobacter_flavus_Ellin428.contig_contig_1
Verrucomicrobia_bacterium_13_1_40CM_4_54_4.contig_contig_1
Verrucomicrobia_bacterium_13_2_20CM_55_10.contig_contig_1
Verrucomicrobia_bacterium_13_2_20CM_2_54_15_9cls.contig_contig_1
Verrucomicrobia_bacterium_13_1_20CM_54_28.contig_contig_1
Verrucomicrobia_bacterium_13_1_20CM_3_54_17.contig_contig_1
Verrucomicrobia_bacterium_13_1_20CM_4_54_11.contig_contig_1
Verrucomicrobia_bacterium_13_2_20CM_54_12.contig_contig_1

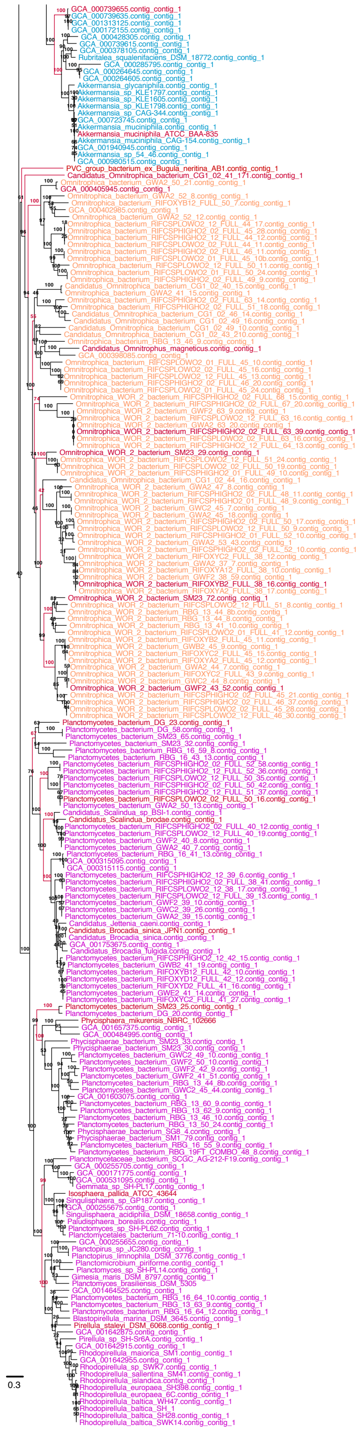

**Supplementary Figure 13**. *Selection of representative PVC genomes*. ML phylogeny, inferred from an alignment of 438 taxa and 2301 sites of concatenated orthologous ribosomal proteins from ribocontigs using RAxML under the PROTCATLG model of evolution. Branch support was estimated with 100 rapid bootstrap replicates. PVC phyla are coloured: Planctomycetes in pink, *Candidatus* Omnitrophica in orange, Verrucomicrobia in blue, Lentisphaerae in green and Chlamydiae in purple. Representative bacterial lineages are in black and the archaeal outgroup in grey. Branches leading to clades from which to select a representative and selected representatives (Supplementary Table 3 and 6) are in red.

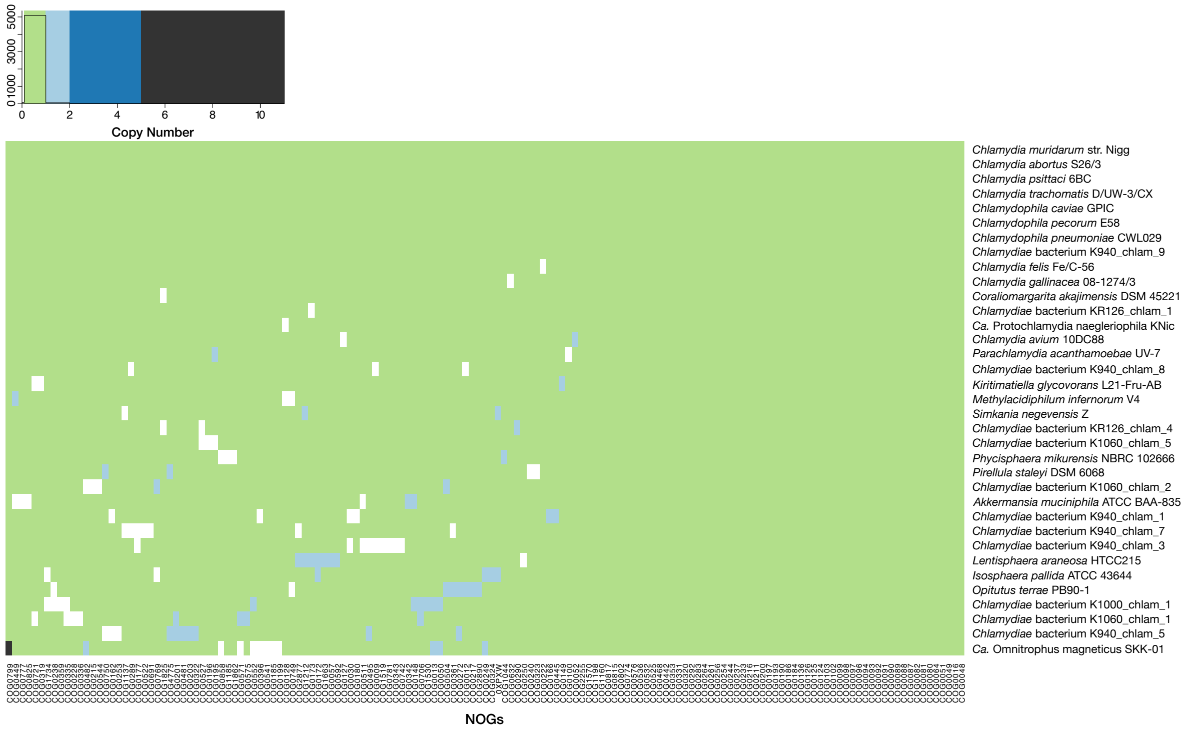

**Supplementary Figure 14**. *Heatmap of copy-number for potential marker gene NOGs*. Presence, absence and copy number of single-copy marker gene NOGs from complete and near complete PVC genomes, used for reconstructing species phylogenies.

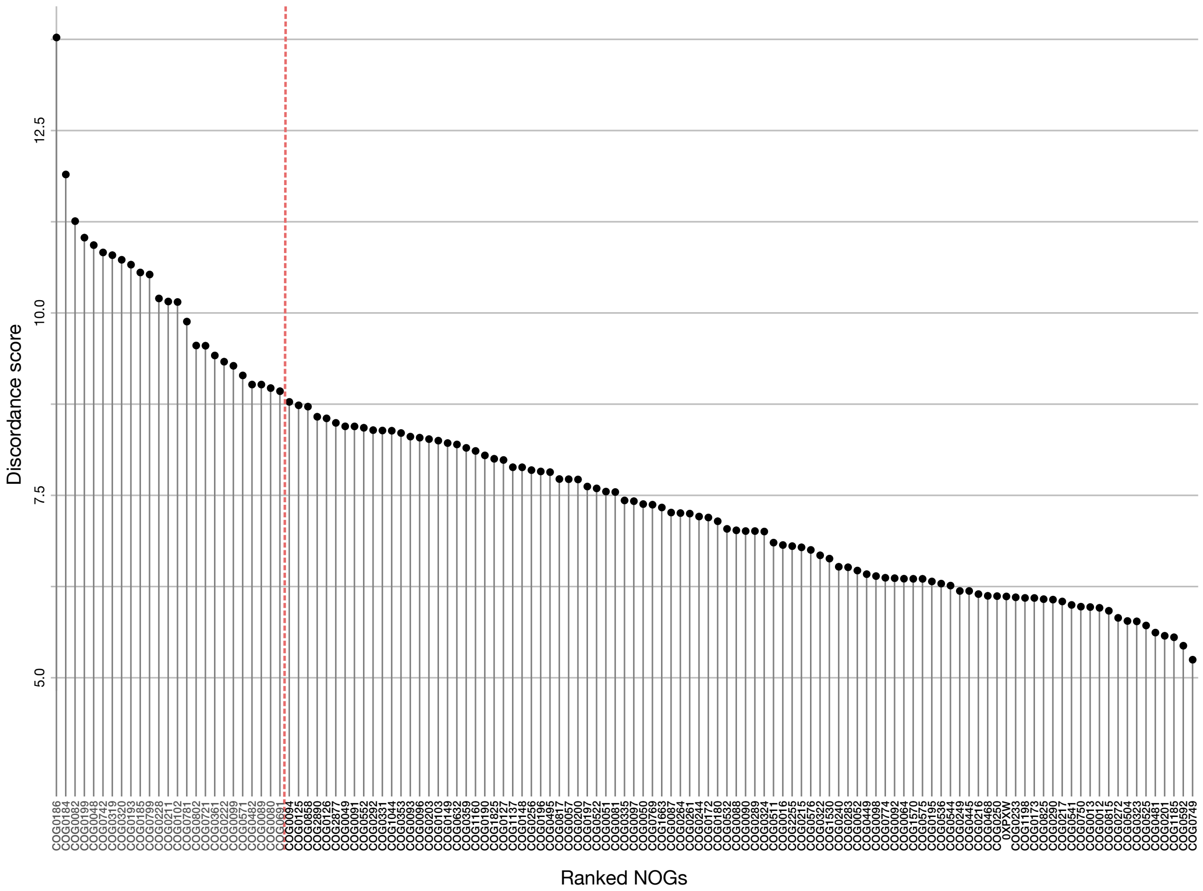

**Supplementary Figure 15**. *Discordance filtering of single-copy marker genes*. Discordance scores across single-copy marker protein NOGs. The 20% most discordant markers are left of the red dotted line.

.

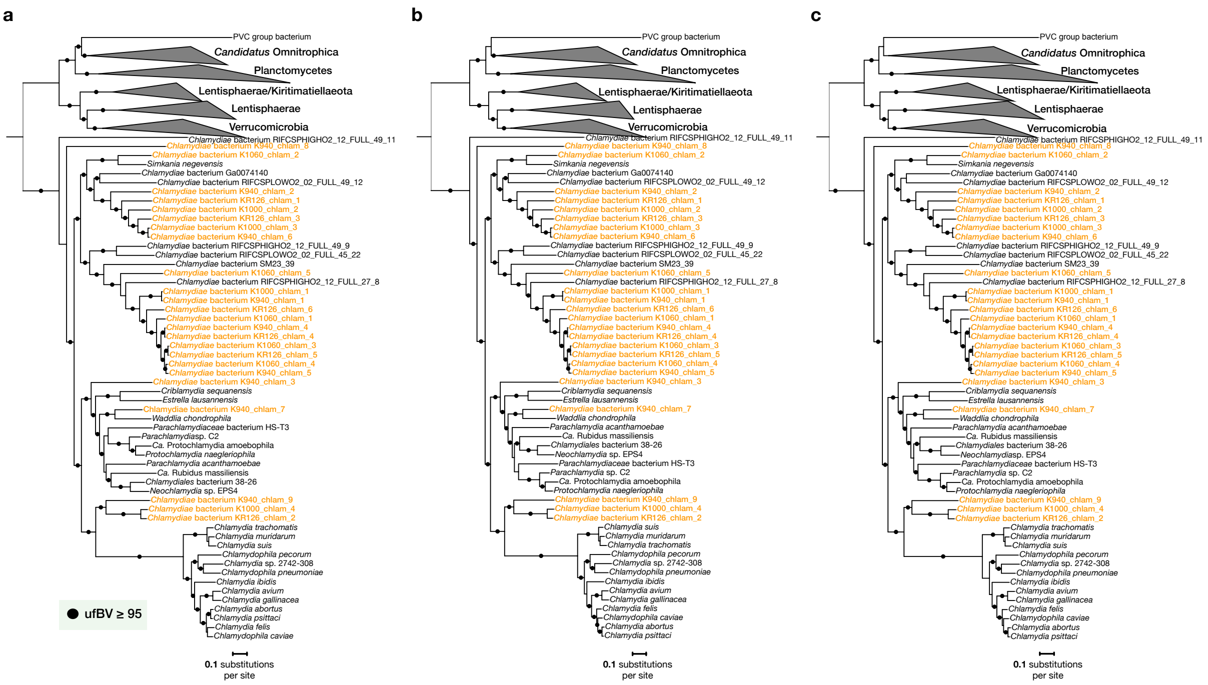

**Supplementary Figure 16**. *ML concatenated species phylogenetic trees of* Chlamydiae. ML phylogenetic trees inferred using IQ-TREE under the LG+C60+R4+F model of evolution with (**a**) 98 (28,286 sites), (**b**) 55 (14,212 sites), and (**c**) 38 (7,894 sites) concatenated single-copy marker genes. Datasets of 55 and 38 single-copy marker genes are subsets of the 98 based on best representation among genomes. Phylogenies include an extensive outgroup with representatives from across the PVC phyla: Kiritimatiellaeota, Lentisphaerae, Verrucomicrobia, *Candidatus* Omnitrophica and Planctomycetes. Ultrafast bootstrap (ufBV) support is indicated at branches following the legend.

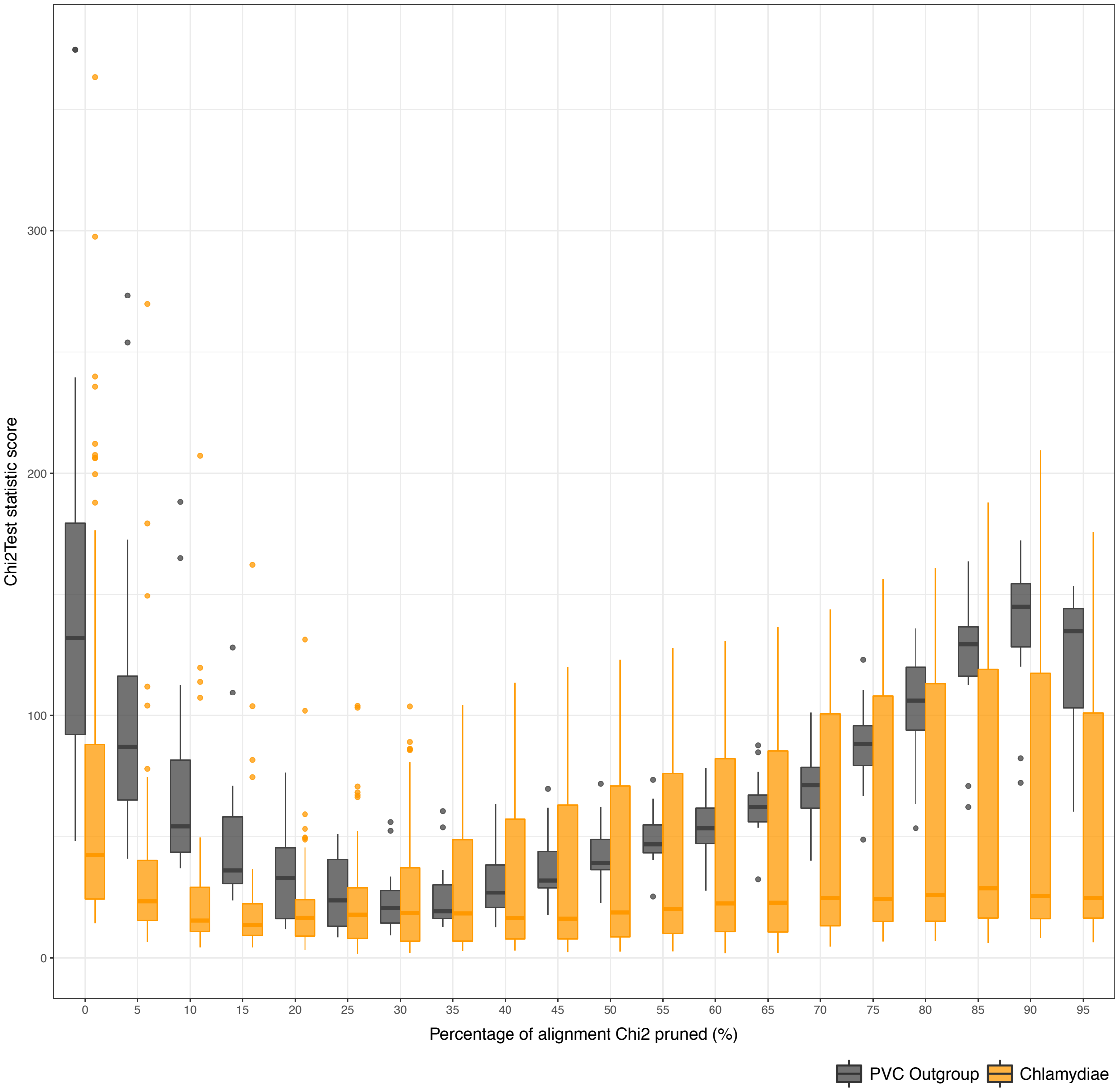

**Supplementary Figure 17**. *Step-wise removal of the most compositionally heterogeneous sites from a concatenated alignment of single-copy marker proteins, based on $\chi^2$ statistics*. The plot shows the $\chi^2$ test statistic across Chlamydiae and outgroup PVC taxa after pruning 5% to 95% of the sites with the highest compositional heterogeneity.

### Supplementary Tables

**Supplementary Table 1**. *Loki's Castle sediment sample information and summary of amplicon sequencing results.*

| Sample ID | Sediment Core | Depth (mbsf) | Chlamydiae Relative Abundance (%) | Number of Chlamydiae OTUs | Chlamydiae OTUs Over 0.1% Relative Abundance | Chlamydiae OTUs Over 1% Relative Abundance |
|---|---|---|---|---|---|---|
| GS10_PC15_10 | GS10_PC15 | 11.58 | –[a] | – | – | – |
| GS10_PC15_40 | GS10_PC15 | 11.23 | – | – | – | – |
| GS10_PC15_70 | GS10_PC15 | 10.93 | – | – | – | – |
| GS10_PC15_100 | GS10_PC15 | 10.63 | – | – | – | – |
| GS10_PC15_130 | GS10_PC15 | 10.33 | – | – | – | – |
| GS10_PC15_160 | GS10_PC15 | 10.03 | – | – | – | – |
| GS10_PC15_190 | GS10_PC15 | 9.73 | – | – | – | – |
| GS10_PC15_220 | GS10_PC15 | 9,43 | 0 | 0 | 0 | 0 |
| GS10_PC15_250 | GS10_PC15 | 9.13 | 0.068 | 1 | 0 | 0 |
| GS10_PC15_280 | GS10_PC15 | 8.83 | – | – | – | – |
| GS10_PC15_310 | GS10_PC15 | 8.53 | – | – | – | – |
| GS10_PC15_340 | GS10_PC15 | 8.23 | – | – | – | – |
| GS10_PC15_370 | GS10_PC15 | 7.93 | – | – | – | – |
| GS10_PC15_400 | GS10_PC15 | 7.63 | – | – | – | – |
| GS10_PC15_430 | GS10_PC15 | 7.33 | – | – | – | – |
| GS10_PC15_460 | GS10_PC15 | 7.03 | – | – | – | – |
| GS10_PC15_490 | GS10_PC15 | 6.73 | – | – | – | – |
| GS10_PC15_520 | GS10_PC15 | 6.43 | 0.177 | 1 | 1 | 0 |
| GS10_PC15_550 | GS10_PC15 | 6.13 | 0 | 0 | 0 | 0 |
| GS10_PC15_580 | GS10_PC15 | 5.83 | – | – | – | – |
| GS10_PC15_610 | GS10_PC15 | 5.53 | 0 | 0 | 0 | 0 |
| GS10_PC15_640 | GS10_PC15 | 5.23 | – | – | – | – |
| GS10_PC15_670 | GS10_PC15 | 4.93 | – | – | – | – |
| GS10_PC15_700 | GS10_PC15 | 4.36 | – | – | – | – |
| GS10_PC15_730 | GS10_PC15 | 4.33 | – | – | – | – |
| GS10_PC15_760 | GS10_PC15 | 4.03 | – | – | – | – |
| GS10_PC15_790 | GS10_PC15 | 3.73 | – | – | – | – |
| GS10_PC15_820 | GS10_PC15 | 3.43 | – | – | – | – |
| GS10_PC15_850 | GS10_PC15 | 3.13 | – | – | – | – |
| GS10_PC15_880 | GS10_PC15 | 2.83 | 3.808 | 14 | 6 | 1 |
| GS10_PC15_910 | GS10_PC15 | 2.53 | – | – | – | – |
| GS10_PC15_940 | GS10_PC15 | 2.23 | 11.148 | 26 | 10 | 2 |
| GS10_PC15_1000 | GS10_PC15 | 1.63 | 5.666 | 82 | 16 | 0 |
| GS10_PC15_1060 | GS10_PC15 | 1.03 | 12.43 | 163 | 29 | 1 |
| GS10_PC15_1090 | GS10_PC15 | 0.73 | – | – | – | – |
| GS10_PC15_1120 | GS10_PC15 | 0.43 | 1.239 | 25 | 2 | 0 |
| GS10_GC14_5 | GS10_GC14 | 0.05 | – | – | – | – |
| GS10_GC14_10 | GS10_GC14 | 0.10 | – | – | – | – |
| GS10_GC14_40 | GS10_GC14 | 0.40 | – | – | – | – |
| GS10_GC14_75 | GS10_GC14 | 0.75 | 8.929 | 10 | 4 | 1 |
| GS10_GC14_100 | GS10_GC14 | 1.00 | – | – | – | – |
| GS10_GC14_115 | GS10_GC14 | 1.15 | – | – | – | – |
| GS10_GC14_130 | GS10_GC14 | 1.30 | – | – | – | – |
| GS10_GC14_150 | GS10_GC14 | 1.50 | – | – | – | – |
| GS10_GC14_176 | GS10_GC14 | 1.76 | – | – | – | – |
| GS10_GC14_180 | GS10_GC14 | 1.80 | – | – | – | – |
| GS10_GC14_200 | GS10_GC14 | 2.00 | – | – | – | – |
| GS14_GC12_10 | GS14_GC12 | 0.10 | 1.024 | 25 | 2 | 0 |
| GS14_GC12_20 | GS14_GC12 | 0.20 | 1.028 | 19 | 1 | 0 |
| GS14_GC12_30 | GS14_GC12 | 0.30 | 1.079 | 94 | 2 | 0 |
| GS14_GC12_40 | GS14_GC12 | 0.40 | 0.467 | 25 | 1 | 0 |
| GS14_GC12_50 | GS14_GC12 | 0.50 | 1.107 | 17 | 3 | 0 |
| GS14_GC12_75 | GS14_GC12 | 0.75 | 2.101 | 45 | 3 | 0 |
| GS14_GC12_100 | GS14_GC12 | 1.00 | 1.648 | 25 | 6 | 0 |
| GS14_GC12_130 | GS14_GC12 | 1.30 | 1.393 | 6 | 3 | 1 |
| GS14_GC12_160 | GS14_GC12 | 1.60 | 3.829 | 4 | 1 | 1 |
| GS14_GC12_175 | GS14_GC12 | 1.75 | 0.848 | 5 | 2 | 0 |
| GS14_GC12_190 | GS14_GC12 | 1.90 | 0.098 | 2 | 0 | 0 |
| GS14_GC12_220 | GS14_GC12 | 2.20 | 0 | 0 | 0 | 0 |
| GS14_GC12_250 | GS14_GC12 | 2.50 | 0 | 0 | 0 | 0 |
| GS14_GC12_280 | GS14_GC12 | 2.80 | 0 | 0 | 0 | 0 |
| GS14_GC12_310 | GS14_GC12 | 3.10 | 0 | 0 | 0 | 0 |
| GS14_GC12_340 | GS14_GC12 | 3.40 | 0.218 | 6 | 0 | 0 |
| GS14_GC12_357 | GS14_GC12 | 3.57 | 0 | 0 | 0 | 0 |
| GS14_GC12_360 | GS14_GC12 | 3.60 | 0.027 | 5 | 0 | 0 |
| GS08_GC12_38 | GS08_GC12 | 0.38 | – | – | – | – |
| GS08_GC12_80 | GS08_GC12 | 0.80 | – | – | – | – |
| GS08_GC12_126 | GS08_GC12 | 1.26 | 43.063 | 37 | 8 | 2 |
| GS08_GC12_310 | GS08_GC12 | 3.10 | – | – | – | – |

[a]Not applicable (–), PCR screened, but amplicon sequencing not performed for sample

#### **Supplementary Table 2**. *Characteristics of marine sediment chlamydiae MAGs.*

| Genome | *Chlamydiae* Clade Affiliation | Metagenome Sample | Completeness (%)[a] | Redundancy[a] | GC (%) | Number of Contigs | Bin Size (Mbp) | Estimated Genome Size (Mbp)[b] | Median Intergenic Space (bp) | iRep[c] | N50 | RP15 Contig | 16S rRNA Gene |
|---|---|---|---|---|---|---|---|---|---|---|---|---|---|
| *Chlamydiae* bacterium K940_chlam_8 | Unresolved | GS10_PC15_940 | 98 | 1 | 37.48 | 89 | 1.4 | 1.43 | 38 | – | 26189 | contig-124_1042 | full (contig-124_2389) |
| *Chlamydiae* bacterium K1060_chlam_2 | CC-I | GS10_PC15_1060 | 97 | 1 | 46.89 | 143 | 1.63 | 1.68´ | 28 | – | 16440 | contig-124_2150 | partial (contig-124_100491) |
| *Chlamydiae* bacterium K940_chlam_2 | CC-II | GS10_PC15_940 | 94 | 1.09 | 48.88 | 285 | 1.66 | 1.61 | 18 | – | 7020 | contig-124_2839 | none |
| *Chlamydiae* bacterium KR126_chlam_1 | CC-II | GS08_GC12_126 | 99 | 1.01 | 46.8 | 25 | 1.61 | 1.61 | 32 | 1.19 | 116054 | contig-100_216 | partial (contig-100_165) |
| *Chlamydiae* bacterium K1000_chlam_2 | CC-II | GS10_PC15_1000 | 88 | 1.01 | 44.48 | 252 | 1.74 | 1.96 | 20 | – | 8520 | contig-124_9157 | partial (contig-124_16903) |
| *Chlamydiae* bacterium KR126_chlam_3 | CC-II | GS08_GC12_126 | 94 | 1 | 42.98 | 185 | 1.68 | 1.79 | 40 | 1.4 | 11258 | contig-100_1304 | partial (contig-100_50271) |
| *Chlamydiae* bacterium K940_chlam_6 | CC-II | GS10_PC15_940 | 91 | 1 | 42.93 | 432 | 1.51 | 1.66 | 24 | – | 4668 | contig-124_54975/contig-124_65424 | none |
| *Chlamydiae* bacterium K1000_chlam_3 | CC-II | GS10_PC15_1000 | 86 | 1 | 41.77 | 270 | 1.66 | 1.93 | 31 | 1.9 | 7870 | contig-124_1375 | none |
| *Chlamydiae* bacterium K1060_chlam_5 | CC-IV | GS10_PC15_1060 | 98 | 1.01 | 26.37 | 57 | 1.39 | 1.40 | 39 | – | 39842 | contig-124_32386 | none |
| *Chlamydiae* bacterium K940_chlam_1 | CC-IV | GS10_PC15_940 | 100 | 1 | 30.93 | 156 | 1.33 | 1.33 | 50 | 1.54 | 14062 | contig-124_2150 | none |
| *Chlamydiae* bacterium K1000_chlam_1 | CC-IV | GS10_PC15_1000 | 96 | 1 | 30.85 | 252 | 1.6 | 1.67 | 57 | 1.51 | 9399 | contig-124_1902 | partial (contig-124_139984) |
| *Chlamydiae* bacterium KR126_chlam_6 | CC-IV | GS08_GC12_126 | 94 | 1.02 | 29.65 | 162 | 1.53 | 1.60 | 50 | 1.36 | 12844 | contig-100_3930 | none |
| *Chlamydiae* bacterium K1060_chlam_1 | CC-IV | GS10_PC15_1060 | 96 | 1 | 29.32 | 120 | 1.59 | 1.66 | 68 | 1.48 | 18228 | contig-124_9382 | none |
| *Chlamydiae* bacterium KR126_chlam_4 | CC-IV | GS08_GC12_126 | 100 | 1.01 | 29.43 | 107 | 1.58 | 1.56 | 63 | 1.12 | 21756 | contig-100_2916 | none |
| *Chlamydiae* bacterium K940_chlam_4 | CC-IV | GS10_PC15_940 | 94 | 1 | 29.51 | 235 | 1.42 | 1.51 | 64 | 1.58 | 8098 | contig-124_7225 | none |
| *Chlamydiae* bacterium K940_chlam_5 | CC-IV | GS10_PC15_940 | 99 | 1.01 | 30.16 | 191 | 1.76 | 1.76 | 65 | 1.58 | 12088 | contig-124_5410 | none |
| *Chlamydiae* bacterium K1060_chlam_4 | CC-IV | GS10_PC15_1060 | 67 | 1.04 | 30.3 | 514 | 1.38 | 1.98 | 75 | – | 3239 | contig-124_114082 | none |
| *Chlamydiae* bacterium KR126_chlam_5 | CC-IV | GS08_GC12_126 | 94 | 1.01 | 30.25 | 264 | 1.57 | 1.65 | 54 | 1.19 | 7741 | contig-100_14168 | none |
| *Chlamydiae* bacterium K1060_chlam_3 | CC-IV | GS10_PC15_1060 | 71 | 1.03 | 30.34 | 167 | 0.98 | 1.34 | 60.5 | – | 8308 | contig-124_29952 | none |
| *Chlamydiae* bacterium K940_chlam_3 | Environmental chlamydiae | GS10_PC15_940 | 96 | 1 | 41.7 | 221 | 1.91 | 1.99 | 50 | 1.73 | 10099 | contig-124_6068 | none |
| *Chlamydiae* bacterium K940_chlam_7 | Environmental chlamydiae | GS10_PC15_940 | 96 | 1 | 43.37 | 408 | 2.49 | 2.59 | 53 | 1.4 | 7595 | contig-124_4023 | none |
| *Chlamydiae* bacterium K940_chlam_9 | CC-V | GS10_PC15_940 | 99 | 1 | 47.21 | 240 | 2.07 | 2.09 | 35.5 | 1.79 | 16710 | contig-124_6236 | partial (contig-124_2865) |
| *Chlamydiae* bacterium KR126_chlam_2 | CC-V | GS08_GC12_126 | 93 | 1.04 | 47.37 | 141 | 1.37 | 1.41 | 24 | 1.4 | 12860 | contig-100_6141/contig-100_8385 | none |
| *Chlamydiae* bacterium K1000_chlam_4 | CC-V | GS10_PC15_1000 | 71 | 1.04 | 46.82 | 229 | 0.94 | 1.27 | 38 | – | 4244 | contig-124_70302 | none |

[a]Estimated using micomplete (See Methods) with Chlamydiae-specific single-copy gene set (Supplementary Table 6)

[b]Based on estimated completeness, corrected by estimated proportion of genome redundancy

[c]Not applicable (–), genome didn't meet coverage (5X) and completeness (70%) requirements for inferring replication rate (iRep)(Brown et al., 2016)

[d]Percentage of reads in respective metagenome mapped to contigs in each genome (out of all reads mapped to contigs in the complete metagenome assembly)

### **Supplementary Table 3**. *Characteristics of Chlamydiae reference genomes.*

| Organism Name | *Chlamydiae* Clade Affiliation | Genbank Accession | Assembly Level | Genome Source | Number of Contigs | Completeness (%)[b] | Redundancy[b] | GC (%) | Genome/Bin Size (Mbp) | Estimated Genome Size (Mbp) | Number of ORFs[c] | Median Intergenic Space (bp) | 16S rRNA Gene |
|---|---|---|---|---|---|---|---|---|---|---|---|---|---|
| ***Chlamydiae* Species Representatives (Available Prior to February 2017)** | | | | | | | | | | | | | |
| *Chlamydiae* bacterium RIFCSPHIGHO2_12_FULL_49_11 | Unresolved | GCA_001794905.1 | Scaffold | aquifer groundwater metagenome[96] | 134 | 91 | 1 | 48.45 | 1.26 | 1.38 | 1065 | 22.5 | no |
| *Simkania negevensis* Z | CC-I | GCA_000237205.1 | Complete Genome | co-culture | – | 100 | 1 | 41.60 | 2.63 | – | 2518 | 36 | yes |
| *Chlamydiae* bacterium RIFCSPLOWO2_02_FULL_49_12 | CC-II | GCA_001796275.1 | Scaffold | aquifer groundwater metagenome[96] | 156 | 95 | 1 | 48.99 | 1.41 | 1.49 | 1175 | 34 | yes |
| *Chlamydiae* bacterium Ga0074140 | CC-II | GCA_001464115.1 | Contig | water treatment plant metagenome[88] | 6 | 99 | 1 | 47.82 | 1.72 | 1.74 | 1648 | 35 | yes |
| *Chlamydiae* bacterium RIFCSPHIGHO2_12_FULL_49_9 | CC-III | GCA_001794935.1 | Scaffold | aquifer groundwater metagenome[96] | 211 | 65 | 1 | 48.59 | 1.32 | 2.02 | 1166 | 27 | no |
| *Chlamydiae* bacterium RIFCSPLOWO2_02_FULL_45_22 | CC-III | GCA_001796255.1 | Scaffold | aquifer groundwater metagenome[96] | 31 | 99 | 1 | 44.70 | 1.58 | 1.59 | 1475 | 23.5 | yes |
| *Chlamydiae* bacterium SM23_39 | CC-IV | GCA_001303765.1 | Contig | aquifer groundwater metagenome[96] | 67 | 93 | 1 | 26.23 | 1.13 | 1.21 | 986 | 35 | yes |
| *Chlamydiae* bacterium RIFCSPHIGHO2_12_FULL_27_8 | CC-IV | GCA_001796155.1 | Scaffold | aquifer groundwater metagenome[96] | 222 | 71 | 1 | 27.43 | 0.97 | 1.38 | 817 | 29 | no |
| *Criblamydia sequanensis* CRIB-18 | Environmental chlamydiae | GCA_000750955.1 | Contig | co-culture | 23 | 99 | 1 | 38.24 | 2.97 | 3.00 | 2418 | 64 | yes |
| *Estrella lausannensis* CRIB-30 | Environmental chlamydiae | GCA_900000175.1 | Scaffold | co-culture | 34 | 99 | 1 | 48.22 | 2.83 | 2.85 | 2217 | 92 | yes |
| *Waddlia chondrophila* WSU 86-1044 | Environmental chlamydiae | GCA_000092785.1 | Complete Genome | co-culture | – | 100 | 1 | 43.74 | 2.13 | – | 1956 | 22 | yes |
| *Parachlamydia acanthamoebae* UV-7 | Environmental chlamydiae | GCA_000253035.1 | Complete Genome | co-culture | – | 99 | 1 | 39.04 | 3.07 | – | 2788 | 56 | yes |
| *Ca.* Rubidus massiliensis (*ex.* Chlamydia sp. Rubis) | Environmental chlamydiae | GCA_000756735.1 | Contig | co-culture | 5 | 100 | 1 | 32.64 | 2.82 | 2.82 | 2446 | 53 | yes |
| *Chlamydiales* bacterium 38-26 | Environmental chlamydiae | GCA_001897225.1 | Scaffold | thiocyanate bioreactor metagenome[87] | 10 | 100 | 1 | 38.12 | 2.83 | 2.83 | 2327 | 88 | no |
| *Neochlamydia* sp. EPS4 | Environmental chlamydiae | GCA_000813665.1 | Contig | co-culture | 112 | 99 | 1 | 38.09 | 2.53 | 2.55 | 2173 | 142 | yes |
| *Parachlamydiaceae* bacterium HS-T3 | Environmental chlamydiae | GCA_000829755.1 | Contig | co-culture | 34 | 99 | 1 | 38.71 | 2.31 | 2.33 | 2025 | 44.5 | yes |
| *Parachlamydia* sp. C2 (*ex.* Protochlamydia greubae) | Environmental chlamydiae | GCA_001545115.1 | Scaffold | co-culture | 33 | 100 | 1 | 42.05 | 3.42 | 3.42 | 2766 | 117 | yes |
| *Ca.* Protochlamydia amoebophila UWE25 | Environmental chlamydiae | GCA_000011565.1 | Chromosome | co-culture | 2 | 100 | 1 | 34.72 | 2.41 | 2.41 | 2031 | 120 | yes |
| *Ca.* Protochlamydia naegleriophila KNic | Environmental chlamydiae | GCA_001499655.1 | Complete Genome | co-culture | –[a] | 100 | 1 | 42.44 | 3.03 | – | 2575 | 113 | yes |
| *Chlamydia trachomatis* D/UW-3/CX | *Clamydiaceae* | GCA_000008725.1 | Complete Genome | co-culture | – | 100 | 1 | 41.31 | 1.04 | – | 894 | 53 | yes |
| *Chlamydia muridarum* str. Nigg | *Clamydiaceae* | GCA_000006685.1 | Complete Genome | co-culture | – | 100 | 1 | 40.31 | 1.08 | – | 911 | 45.5 | yes |
| *Chlamydia suis* MD56 | *Clamydiaceae* | GCA_000493885.1 | Scaffold | co-culture | 47 | 100 | 1 | 42.01 | 1.08 | 1.08 | 931 | 51 | yes |
| *Chlamydophila pecorum* E58 | *Clamydiaceae* | GCA_000204135.1 | Complete Genome | co-culture | – | 100 | 1 | 41.08 | 1.11 | – | 988 | 28 | yes |
| *Chlamydia* sp. 2742-308 | *Clamydiaceae* | GCA_001653975.1 | Chromosome | co-culture | 2 | 100 | 1 | 38.50 | 1.12 | 1.12 | 1004 | 44 | yes |
| *Chlamydophila pneumoniae* CWL029 | *Clamydiaceae* | GCA_000008745.1 | Complete Genome | co-culture | – | 100 | 1 | 40.58 | 1.23 | – | 1052 | 55 | yes |
| *Chlamydia ibidis* 10-1398/6 | *Clamydiaceae* | GCA_000454725.1 | Contig | co-culture | 4 | 100 | 1 | 38.32 | 1.15 | 1.15 | 1018 | 50 | yes |
| *Chlamydia avium* 10DC88 | *Clamydiaceae* | GCA_000583875.1 | Complete Genome | co-culture | – | 100 | 1 | 36.88 | 1.05 | – | 947 | 31 | yes |
| *Chlamydia gallinacea* 08-1274/3 | *Clamydiaceae* | GCA_000471025.2 | Complete Genome | co-culture | – | 99 | 1 | 37.90 | 1.07 | – | 900 | 40.5 | yes |
| *Chlamydia felis* Fe/C-56 | *Clamydiaceae* | GCA_000009945.1 | Complete Genome | co-culture | – | 100 | 1 | 39.34 | 1.17 | – | 1013 | 39.5 | yes |
| *Chlamydophila caviae* GPIC | *Clamydiaceae* | GCA_000007605.1 | Complete Genome | co-culture | – | 100 | 1 | 39.19 | 1.18 | – | 1005 | 48 | yes |
| *Chlamydia abortus* S26/3 | *Clamydiaceae* | GCA_000026025.1 | Complete Genome | co-culture | – | 100 | 1 | 39.87 | 1.14 | – | 932 | 45 | yes |
| *Chlamydia psittaci* 6BC | *Clamydiaceae* | GCA_000204255.1 | Complete Genome | co-culture | – | 100 | 1 | 39.02 | 1.18 | – | 1009 | 42 | yes |
| **Other *Chlamydiae* Species Representatives (Released Between February 2017 and April 2018)** | | | | | | | | | | | | | |
| *Ca.* Similichlamydia epinephelii GCCT14 | Unresolved | GCA_003056015.1 | Scaffold | infected gill tissue metagenome[2] | 170 | 80 | 1.4 | 39.54 | 0.98 | 0.71 | 940 | 36 | yes |
| *Rhabdochlamydia helvetica* T3358 | CC-II | Pillonel et al.. 2018 | Contig | tick metagenome[3] | 38 | 99 | 1 | 36.18 | 1.83 | 1.85 | 1692 | 43 | yes |
| *Chlamydiae* bacterium CG10_big_fil_rev_8_21_14_0_10_42_34 | CC-III | GCA_002773795.1 | Scaffold | cold $CO_2$ driven geyser metagenome[95] | 34 | 97 | 1 | 42.36 | 1.68 | 1.73 | 1581 | 21.5 | no |
| *Chlamydiae* bacterium CG10_big_fil_rev_8_21_14_0_10_35_9 | Unresolved | GCA_002773835.1 | Scaffold | cold $CO_2$ driven geyser metagenome[95] | 108 | 85 | 1.04 | 35.13 | 1.74 | 1.96 | 1720 | 24 | no |
| *Chlamydiales* bacterium SCGC AB-751-O23 | Unresolved | GCA_900093645.1 | Scaffold | single-cell from marine water[36] | 89 | 42 | 1 | 35.45 | 0.99 | 2.37 | 876 | 30.5 | yes |
| *Waddliaceae* bacterium SP13 | Unresolved | GCA_002709385.1 | Contig | marine water metagenome[37] | 49 | 97 | 1 | 38.48 | 3.15 | 3.24 | 2460 | 65 | yes |
| *Chlamydiales* bacterium SCGC AG-110-P3 | Environmental chlamydiae | GCA_900093655.1 | Scaffold | single-cell from marine water[36] | 102 | 50 | 1 | 46.83 | 1.30 | 2.58 | 1235 | 84 | yes |
| *Chlamydiales* bacterium SCGC AG-110-M15 | Unresolved | GCA_900093625.1 | Scaffold | single-cell from marine water[36] | 59 | 50 | 1 | 41.80 | 0.93 | 1.84 | 851 | 53 | no |
| *Parachlamydia* sp. BC.030 | Environmental chlamydiae | GCA_002786175.1 | Contig | urban drinking water system metagenome[89] | 39 | 100 | 1 | 41.53 | 3.04 | 3.04 | 2540 | 68 | no |
| *Ca.* Chlamydia corallus G3/2742-324 | *Clamydiaceae* | GCA_002817655.1 | Contig | snake choana metagenome[97] | 7 | 100 | 1.01 | 39.28 | 1.20 | 1.19 | 996 | 48.5 | yes |
| **Non-representative *Chlamydiae* Included in Select Analyses** | | | | | | | | | | | | | |
| *Chlamydiae* bacterium GWA2_50_15 | CC-II | GCA_001796065.1 | Scaffold | aquifer groundwater metagenome[96] | 56 | 94 | 1 | 49.34 | 1.18 | 1.26 | 993 | 29.5 | yes |
| *Chlamydiae* bacterium GWC2_50_10 | CC-II | GCA_001796095.1 | Scaffold | aquifer groundwater metagenome[96] | 135 | 83 | 1 | 48.94 | 1.17 | 1.41 | 966 | 31 | yes |
| *Chlamydiae* bacterium GWF2_49_8 | CC-II | GCA_001796105.1 | Scaffold | aquifer groundwater metagenome[96] | 280 | 71 | 1 | 49.23 | 1.02 | 1.44 | 760 | 26 | no |
| *Chlamydiae* bacterium RIFCSPHIGHO2_02_FULL_49_29 | CC-II | GCA_001796185.1 | Scaffold | aquifer groundwater metagenome[96] | 85 | 91 | 1 | 49.08 | 1.39 | 1.52 | 1187 | 31 | no |
| *Chlamydiae* bacterium RIFCSPHIGHO2_12_FULL_49_32 | CC-II | GCA_001796175.1 | Scaffold | aquifer groundwater metagenome[96] | 94 | 89 | 1 | 48.91 | 1.40 | 1.56 | 1190 | 29 | yes |
| *Chlamydiae* bacterium RIFCSPLOWO2_12_FULL_49_12 | CC-II | GCA_001796315.1 | Scaffold | aquifer groundwater metagenome[96] | 70 | 90 | 1 | 49.16 | 1.42 | 1.57 | 1224 | 34.5 | yes |
| *Chlamydiae* bacterium RIFCSPHIGHO2_01_FULL_44_39 | CC-III | GCA_001794865.1 | Scaffold | aquifer groundwater metagenome[96] | 32 | 99 | 1 | 44.72 | 1.57 | 1.58 | 1466 | 23 | yes |
| *Chlamydiae* bacterium RIFCSPHIGHO2_02_FULL_45_9 | CC-III | GCA_001796125.1 | Scaffold | aquifer groundwater metagenome[96] | 222 | 80 | 1 | 44.66 | 1.34 | 1.69 | 1156 | 26 | no |
| *Chlamydiae* bacterium RIFCSPLOWO2_01_FULL_44_52 | CC-III | GCA_001796235.1 | Scaffold | aquifer groundwater metagenome[96] | 30 | 99 | 1 | 44.74 | 1.54 | 1.55 | 1438 | 24 | yes |
| *Chlamydiae* bacterium RIFCSPLOWO2_12_FULL_45_20 | CC-III | GCA_001796285.1 | Scaffold | aquifer groundwater metagenome[96] | 29 | 99 | 1 | 44.75 | 1.54 | 1.56 | 1443 | 23 | yes |
| *Chlamydiae* bacterium RIFCSPHIGHO2_12_FULL_44_59 | CC-III | GCA_001794895.1 | Scaffold | aquifer groundwater metagenome[96] | 30 | 99 | 1 | 44.72 | 1.57 | 1.58 | 1470 | 23.5 | yes |
| *Chlamydiae* bacterium RIFCSPLOWO2_01_FULL_28_7 | CC-IV | GCA_001796195.1 | Scaffold | aquifer groundwater metagenome[96] | 193 | 62 | 1 | 28.07 | 0.71 | 1.14 | 572 | 31 | no |
| *Chlamydia* sp. 32-24 | Environmental chlamydiae | GCA_001897185.1 | Scaffold | thiocyanate bioreactor metagenome[87] | 100 | 99 | 1 | 32.42 | 2.53 | 2.55 | 2075 | 56 | no |
| *Neochlamydia* sp. S13 | Environmental chlamydiae | GCA_000648235.1 | Contig | co-culture | 1342 | 99 | 1.07 | 38.03 | 3.19 | 2.99 | 2232 | 161 | yes |
| *Neochlamydia* sp. TUME1 | Environmental chlamydiae | GCA_000813645.1 | Contig | co-culture | 254 | 100 | 1 | 38.02 | 2.55 | 2.55 | 2344 | 121 | yes |
| *Parachlamydia acanthamoebae* OEW1 | Environmental chlamydiae | GCA_000812225.1 | Contig | co-culture | 162 | 97 | 1 | 39.04 | 3.01 | 3.09 | 2755 | 53 | yes |
| *Parachlamydia acanthamoebae* Bn9 | Environmental chlamydiae | GCA_000875975.1 | Contig | co-culture | 72 | 99 | 1 | 38.94 | 3.00 | 3.03 | 2409 | 60 | yes |
| *Parachlamydia acanthamoebae* Hall's coccus | Environmental chlamydiae | GCA_000176075.1 | Contig | co-culture | 95 | 98 | 1 | 38.97 | 2.97 | 3.02 | 2809 | 54 | no |
| *Ca.* Protochlamydia amoebophila EI2 | Environmental chlamydiae | GCA_000813625.1 | Contig | co-culture | 178 | 96 | 1 | 34.82 | 2.40 | 2.51 | 2149 | 99 | yes |
| *Ca.* Protochlamydia massiliensis (*ex.* Chlamydia sp. 'Diamant') | Environmental chlamydiae | GCA_000751535.1 | Contig | co-culture | 5 | 100 | 1 | 42.75 | 2.96 | 2.96 | 2451 | 110 | yes |
| *Ca.* Protochlamydia sp. R18 str. S13 | Environmental chlamydiae | GCA_000648255.1 | Contig | co-culture | 795 | 100 | 1.02 | 34.74 | 2.72 | 2.67 | 2017 | 110 | yes |
| *Ca.* Protochlamydia sp. W-9 | Environmental chlamydiae | GCA_001950615.1 | Contig | co-culture | 402 | 100 | 1 | 34.43 | 2.48 | 2.48 | 1817 | 109 | yes |

[a] Not applicable (–), complete genome

[b] Estimated using micomplete (See Methods) with Chlamydiae-specific single-copy gene set (Supplementary Table 6)

[c] Open Reading Frames (ORFs)

**Supplementary Table 4**. *Number of 16/18S rRNA gene fragments identified per phyla in marine sediment sample metagenomes.*

| Phylum | Domain | GS10_PC15_1060 | GS10_PC15_1000 | GS10_PC15_940 | GS08_GC12_126 |
|---|---|---|---|---|---|
| Chloroplast | – | 0 | 0 | 1 | 0 |
| Platyhelminthes | Eukaryota | 1 | 0 | 0 | 0 |
| Chordata | Eukaryota | 0 | 1 | 2 | 0 |
| Abeoformidae | Eukaryota | 0 | 0 | 2 | 0 |
| Chlorophyta | Eukaryota | 0 | 0 | 1 | 0 |
| Bacteroidetes | Bacteria | 8 | 6 | 9 | 3 |
| Marinimicrobia (SAR406 clade) | Bacteria | 4 | 3 | 3 | 3 |
| Gemmatimonadetes | Bacteria | 3 | 2 | 3 | 2 |
| Fibrobacteres | Bacteria | 3 | 2 | 1 | 1 |
| Caldithrix phylum incertae sedis | Bacteria | 2 | 3 | 0 | 1 |
| *Ca.* Latescibacteria (WS3) | Bacteria | 4 | 4 | 2 | 2 |
| Cloacimonetes | Bacteria | 1 | 0 | 0 | 0 |
| *Ca.* Zixibacteria (RBG-1) | Bacteria | 5 | 3 | 0 | 5 |
| GN01 | Bacteria | 0 | 2 | 0 | 0 |
| Ignavibacteriae | Bacteria | 0 | 0 | 1 | 0 |
| Proteobacteria | Bacteria | 51 | 22 | 30 | 14 |
| Firmicutes | Bacteria | 0 | 1 | 1 | 0 |
| Actinobacteria | Bacteria | 17 | 18 | 21 | 8 |
| Chloroflexi | Bacteria | 63 | 50 | 22 | 92 |
| Deinococcus-Thermus | Bacteria | 1 | 2 | 0 | 0 |
| Armatimonadetes | Bacteria | 1 | 1 | 1 | 0 |
| WS1 | Bacteria | 3 | 2 | 1 | 5 |
| Spirochaetes | Bacteria | 9 | 8 | 7 | 3 |
| **Chlamydiae** | **Bacteria** | **16** | **17** | **12** | **6** |
| Planctomycetes | Bacteria | 69 | 45 | 55 | 31 |
| Lentisphaerae | Bacteria | 1 | 0 | 0 | 0 |
| *Ca.* Omnitrophica (OP3) | Bacteria | 7 | 1 | 0 | 11 |
| Epsilonbacteraeota | Bacteria | 3 | 0 | 3 | 0 |
| Acidobacteria | Bacteria | 6 | 5 | 5 | 0 |
| *Ca.* Acetothermia (OP1) | Bacteria | 0 | 4 | 0 | 1 |
| Nitrospinae | Bacteria | 0 | 0 | 0 | 1 |
| Elusimicrobia | Bacteria | 2 | 1 | 0 | 1 |
| *Ca.* Hydrogenedentes (NKB19) | Bacteria | 1 | 0 | 2 | 1 |
| *Ca.* Atribacteria | Bacteria | 0 | 0 | 2 | 0 |
| *Ca.* Saccharibacteria (TM7) | Bacteria | 1 | 1 | 0 | 0 |
| *Ca.* Parcubacteria | Bacteria | 49 | 22 | 13 | 11 |
| CPR2 | Bacteria | 2 | 0 | 0 | 0 |
| *Ca.* Microgenomates (OP11) | Bacteria | 9 | 6 | 4 | 9 |
| *Ca.* Berkelbacteria | Bacteria | 2 | 2 | 1 | 1 |
| *Ca.* Gracilibacteria | Bacteria | 2 | 0 | 0 | 0 |
| *Ca.* Peregrinibacteria | Bacteria | 4 | 0 | 0 | 1 |
| *Ca.* Katanobacteria (WWE3) | Bacteria | 1 | 0 | 1 | 0 |
| *Ca.* Absconditabacteria (SR1) | Bacteria | 0 | 1 | 0 | 0 |
| MD2896-B216 | Bacteria | 2 | 1 | 0 | 0 |
| Unknown CPR phylum 1 | Bacteria | 1 | 1 | 0 | 0 |
| Unknown CPR phylum 2 | Bacteria | 1 | 0 | 0 | 0 |
| Unknown CPR phylum 3 | Bacteria | 1 | 0 | 0 | 0 |
| Poribacteria | Bacteria | 4 | 1 | 0 | 3 |
| Halanaerobiales phylum incertae sedis | Bacteria | 4 | 5 | 0 | 2 |
| *Ca.* Aminicenantes (OP8) | Bacteria | 3 | 3 | 1 | 1 |
| *Ca.* Dependentiae (TM6) | Bacteria | 42 | 17 | 18 | 6 |
| *Ca.* Aerophobetes (CD12) | Bacteria | 6 | 6 | 1 | 3 |
| NC10 | Bacteria | 1 | 0 | 0 | 1 |
| Nitrospirae | Bacteria | 2 | 1 | 5 | 0 |
| BRC1 | Bacteria | 1 | 1 | 5 | 0 |
| Euryarchaeota | Archaea | 1 | 1 | 0 | 4 |
| Unknown Euryarchaeota (superphylum) phylum 1 | Archaea | 0 | 0 | 1 | 0 |
| Thaumarchaeota | Archaea | 2 | 8 | 4 | 1 |
| *Ca.* Bathyarchaeota | Archaea | 2 | 2 | 0 | 1 |
| Group C3 | Archaea | 2 | 4 | 0 | 0 |
| *Ca.* Woesearchaeota | Archaea | 20 | 11 | 30 | 8 |
| *Ca.* Diapherotrites | Archaea | 3 | 0 | 0 | 0 |
| *Ca.* Parvarchaeota | Archaea | 1 | 0 | 0 | 0 |
| Unknown DPANN phylum 1 | Archaea | 1 | 0 | 0 | 0 |
| *Ca.* Aenigmarchaeota | Archaea | 0 | 0 | 0 | 1 |
| *Ca.* Altiarchaeota | Archaea | 0 | 0 | 0 | 1 |
| *Ca.* Lokiarchaeota | Archaea | 5 | 10 | 1 | 3 |
| **TOTAL** | ALL | 456 | 307 | 271 | 248 |

**Supplementary Table 5**. *PCR primer pairs, taxonomic coverage and reaction conditions.*

| Primer Pair | Taxonomic Target | Taxonomic Coverage (No Mismatches)[a] | Taxonomic Coverage (One Mismatch)[a] | PCR Amplication Reaction Conditions[b] |
|---|---|---|---|---|
| Chla-310-a-20 (CGCCAACAYTGGGACTGAGA) and S-*-Univ-1100-a-A-15 (GGGTYKCGCTCGTTR)[98] | Chlamydiae | · 0% of Eukaryota<br>· 0% of Archaea<br>· 0% of Bacteria<br>· 96% of characterized Chlamydiae (0.1% of Armatimonadetes, no additional bacterial phyla) | · 0% of Eukaryota<br>· 0% of Archaea<br>· 0% of Bacteria<br>· 99 % of characterized Chlamydiae (4.5 % of Marinimicrobia and small percentages (0.01-0.81 %) of additional bacterial phyla) | · 15 min of polymerase heat activation at 95 °C<br>· 35 cycles of 94 °C (60 s), 60 °C (60 s) and 72 °C (60 s) · final extension at 72 °C (10 min) |
| 574*f (CGGTAAYTCCAGCTCYV)[99] and 1132 (CCGTCAATTHCTTYAART)[99] | Eukarya | · 88 % of Eukaryota<br>· 0% of Archaea<br>· 0% of Bacteria | · 94% of Eukaryota<br>· 10% of Archaea<br>· 0% of Bacteria | · 15 min of polymerase heat activation at 95 °C<br>· 35 cycles of 94 °C (60 s), a step-down to 70 °C (1 s), followed by a ramping rate of 0.4 °C/s to 50 °C (60 s), and a ramping rate of 0.8 °C/s to 72 °C (60 s)<br>· final extension at 72 °C (10 min) |
| S-D-0564-a-S-15 (AYTGGGYDTAAAGNG)[98] and S-D-Bact-1061-a-A-17 (CRRCACGAGCTGACGAC)[98] | Bacteria | · 0% of Eukaryota<br>· 0.2% of Archaea<br>· 92% of Bacteria<br>· 94% of characterized Chlamydiae | · 0% of Eukaryota<br>· 4.6% of Archaea<br>· 97% of Bacteria<br>· 99% of characterized Chlamydiae | · 15 min of polymerase heat activation at 95 °C<br>· 28 cycles of 94 °C (60 s), a step-down to 70 °C (1 s), followed by a ramping rate of 0.4 °C/s to 50 °C (60 s), and a ramping rate of 0.8 °C/s to 72 °C (60 s)<br>· final extension at 72 °C (10 min) |
| A519F (CAGCMGCCGCGGTAA)[100] and Uni1391R (ACGGGCGGTGWGTRC)[93] | Eukarya, Archaea and Bacteria | · 86% of Eukaryota<br>· 70% of Archaea<br>· 86% of Bacteria<br>· 0.7% of characterized Chlamydiae | · 93% of Eukaryota<br>· 74% of Archaea<br>· 91% of Bacteria<br>· 93% of characterized Chlamydiae | not used in this study |
| Earth Microbiome Project[92] primers: 515F (GTGYCAGCMGCCGCGGTAA) and 806R (GGACTACNVGGGTWTCTAAT) | Archaea and Bacteria | · 0% of Eukaryota<br>· 0% of Archaea<br>· 92% of Bacteria<br>· 0.7% of characterized Chlamydiae | · 0% of Eukaryota<br>· 0% of Archaea<br>· 92% of Bacteria<br>· 95% of characterized Chlamydiae | not used in this study |

[a]Using SILVA TestPrime (Klindworth et al., 2013) with the SSU r132 database and RefNR sequence collection
[b]Using HotStarTaq DNA Polymerase (QIAGEN)

**Supplementary Table 6**. *PVC outgroup genomes used in phylogenomic analyses.*

| Phylum | Species Name | Genbank Accession |
|---|---|---|
| Kiritimatiellaeota | *Kiritimatiella glycovorans* L21-Fru-AB | GCA_001017655.1 |
| Lentisphaerae | *Lentisphaerae* bacterium GWF2_57_35 | GCA_001804865.1 |
| Lentisphaerae | *Lentisphaerae* bacterium RIFOXYC12_FULL_60_16 | GCA_001803315.1 |
| Lentisphaerae | *Lentisphaera araneosa* HTCC215 | GCA_000170755.1 |
| Lentisphaerae | *Lentisphaerae* bacterium GWF2_50_93 | GCA_001804815.1 |
| Verrucomicrobia | *Coraliomargarita akajimensis* DSM 45221 | GCA_000025905.1 |
| Verrucomicrobia | *Opitutus terrae* PB90-1 | GCA_000019965.1 |
| Verrucomicrobia | *Pedosphaera parvula* Ellin514 | GCA_000172555.1 |
| Verrucomicrobia | *Methylacidiphilum infernorum* V4 | GCA_000019665.1 |
| Verrucomicrobia | *Terrimicrobium sacchariphilum* NM-5 | GCA_001613545.1 |
| Verrucomicrobia | *Verrucomicrobium* sp. BvORR106 | GCA_000739655.1 |
| Verrucomicrobia | *Akkermansia muciniphila* ATCC BAA-835 | GCA_000020225.1 |
| Unclassified PVC | PVC group bacterium (*ex.* Bugula neritina AB1) AB1-3 | GCA_001730085.1 |
| *Candidatus* Omnitrophica | *Ca.* Omnitrophica bacterium CG1_02_41_171 | GCA_001871865.1 |
| *Candidatus* Omnitrophica | *Ca.* Omnitrophus fodinae SCGC AAA011-A17 | GCA_000405945.1 |
| *Candidatus* Omnitrophica | *Ca.* Omnitrophus magneticus SKK-01 | GCA_000954095.1 |
| *Candidatus* Omnitrophica | *Omnitrophica* WOR_2 bacterium RIFCSPHIGHO2_02_FULL_63_39 | GCA_001805685.1 |
| *Candidatus* Omnitrophica | *Omnitrophica* WOR_2 bacterium RIFCSPLOWO2_02_FULL_50_19 | GCA_001805805.1 |
| *Candidatus* Omnitrophica | *Omnitrophica* WOR_2 bacterium RIFOXYB2_FULL_38_16 | GCA_001805995.1 |
| *Candidatus* Omnitrophica | *Omnitrophica* WOR_2 bacterium RBG_13_41_10 | GCA_001805465.1 |
| *Candidatus* Omnitrophica | *Omnitrophica* WOR_2 bacterium GWF2_43_52 | GCA_001805445.1 |
| Planctomycetes | *Planctomycetes* bacterium DG_23 | GCA_001302825.1 |
| Planctomycetes | *Planctomycetes* bacterium RIFCSPLOWO2_02_FULL_50_16 | GCA_001828565.1 |
| Planctomycetes | *Ca.* Scalindua brodae RU1 | GCA_000786775.1 |
| Planctomycetes | *Ca.* JPN1 | GCA_000949635.1 |
| Planctomycetes | *Phycisphaera mikurensis* NBRC 102666 | GCA_000284115.1 |
| Planctomycetes | *Isosphaera pallida* ATCC 43644 | GCA_000186345.1 |
| Planctomycetes | *Pirellula staleyi* DSM 6068 | GCA_000025185.1 |

**Supplementary Table 7**. *Single-copy marker genes used to assess chlamydial MAGs completeness and redundancy.*

| **Chlamydiae Micomplete Gene/Domain Names** | |
|---|---|
| Aminoacyl tRNA synthetase class II, N-terminal domain | Ribosomal protein S10p/S20e |
| Arginyl tRNA synthetase N terminal domain | Ribosomal protein S11 |
| Bacterial RNA polymerase, alpha chain C terminal domain | Ribosomal protein S12/S23 |
| Bacterial trigger factor protein (TF) | Ribosomal protein S13/S18 |
| Bacterial trigger factor protein (TF) C-terminus | Ribosomal protein S15 |
| ClpX C4-type zinc finger | Ribosomal protein S16 |
| Conserved hypothetical protein 95 | Ribosomal protein S17 |
| CTP synthase N-terminus | Ribosomal protein S18 |
| Cytidylate kinase | Ribosomal protein S19 |
| Dephospho-CoA kinase | Ribosomal protein S2 |
| DNA polymerase III beta subunit, C-terminal domain | Ribosomal protein S20 |
| DNA polymerase III beta subunit, central domain | Ribosomal protein S3, C-terminal domain |
| DNA primase catalytic core, N-terminal domain | Ribosomal protein S4/S9 N-terminal domain |
| Double-stranded RNA binding motif | Ribosomal protein S5, C-terminal domain |
| Elongation factor TS | Ribosomal protein S5, N-terminal domain |
| Enolase, C-terminal TIM barrel domain | Ribosomal protein S6 |
| Enolase, N-terminal domain | Ribosomal protein S7p/S5e |
| FAD synthetase | Ribosomal protein S8 |
| Ferredoxin-fold anticodon binding domain | Ribosomal protein S9/S16 |
| GAD domain | Ribosomal Proteins L2, C-terminal domain |
| GrpE | Ribosomal Proteins L2, RNA binding domain |
| GTP-binding protein LepA C-terminus | Ribosome recycling factor |
| GTP1/OBG | RNA polymerase beta subunit |
| Holliday junction DNA helicase ruvB C-terminus | RNA polymerase beta subunit external 1 domain |
| IPP transferase | RNA polymerase Rpb1, domain 1 |
| MraW methylase family | RNA polymerase Rpb1, domain 2 |
| NusA N-terminal domain | RNA polymerase Rpb1, domain 3 |
| Oligomerisation domain | RNA polymerase Rpb1, domain 4 |
| Peptidyl-tRNA hydrolase | RNA polymerase Rpb1, domain 5 |
| Phosphoglycerate kinase | RNA polymerase Rpb2, domain 2 |
| Protein of unknown function (DUF933) | RNA polymerase Rpb2, domain 3 |
| recA bacterial DNA recombination protein | RNA polymerase Rpb2, domain 6 |
| Ribosomal L18p/L5e family | RNA polymerase Rpb2, domain 7 |
| Ribosomal L27 protein | RNA polymerase Rpb3/Rpb11 dimerisation domain |
| Ribosomal L28 family | RuvA N terminal domain |
| Ribosomal L29 protein | SecY translocase |
| ribosomal L5P family C-terminus | Seryl-tRNA synthetase N-terminal domain |
| Ribosomal prokaryotic L21 protein | Signal peptidase (SPase) II |
| Ribosomal protein L10 | Signal peptide binding domain |
| Ribosomal protein L11, N-terminal domain | SmpB protein |
| Ribosomal protein L11, RNA binding domain | Tetrahydrofolate dehydrogenase/cyclohydrolase, NAD(P)-binding domain |
| Ribosomal protein L13 | Translation initiation factor 1A / IF-1 |
| Ribosomal protein L14p/L23e | Translation initiation factor IF-3, C-terminal domain |
| Ribosomal protein L16p/L10e | Translation initiation factor IF-3, N-terminal domain |
| Ribosomal protein L17 | Translation-initiation factor 2 |
| Ribosomal protein L18e/L15 | TRCF domain |
| Ribosomal protein L19 | tRNA (Guanine-1)-methyltransferase |
| Ribosomal protein L1p/L10e family | tRNA synthetase B5 domain |
| Ribosomal protein L20 | tRNA synthetases class I (R) |
| Ribosomal protein L22p/L17e | tRNA synthetases class II core domain (F) |
| Ribosomal protein L23 | UDP-N-acetylenolpyruvoylglucosamine reductase, C-terminal domain |
| Ribosomal protein L3 | Ultra-violet resistance protein B |
| Ribosomal protein L35 | Uncharacterised P-loop hydrolase UPF0079 |
| Ribosomal protein L5 | Uncharacterised protein family (UPF0081) |
| Ribosomal protein L6 | Uncharacterized protein family UPF0054 |
| Ribosomal protein L9, C-terminal domain | UvrC Helix-hairpin-helix N-terminal |
| Ribosomal protein L9, N-terminal domain | |

**Supplementary Table 8.** *Single-copy marker genes used concatenation-based species tree inference.*

| Used in 38, 55 and 98 NOG marker protein sets | | Used in 55 and 98 NOG marker protein sets | | Used in 98 NOG marker protein sets | |
|---|---|---|---|---|---|
| **NOG** | **COG description** | **NOG** | **COG description** | **NOG** | **COG description** |
| COG0049 | Ribosomal protein S7 | 0XPXW | | COG0012 | Ribosome-binding ATPase YchF, GTP1/OBG family |
| COG0051 | Ribosomal protein S10 | COG0013 | Alanyl-tRNA synthetase | COG0064 | Asp-tRNAAsn/Glu-tRNAGln amidotransferase B subunit |
| COG0057 | Glyceraldehyde-3-phosphate dehydrogenase/erythrose-4-phosphate dehydrogenase | COG0016 | Phenylalanyl-tRNA synthetase alpha subunit | COG0125 | Thymidylate kinase |
| COG0081 | Ribosomal protein L1 | COG0050 | Translation elongation factor EF-Tu, a GTPase | COG0148 | Enolase |
| COG0087 | Ribosomal protein L3 | COG0052 | Ribosomal protein S2 | COG0149 | Triosephosphate isomerase |
| COG0088 | Ribosomal protein L4 | COG0172 | Seryl-tRNA synthetase | COG0173 | Aspartyl-tRNA synthetase |
| COG0090 | Ribosomal protein L2 | COG0240 | Glycerol-3-phosphate dehydrogenase | COG0180 | Tryptophanyl-tRNA synthetase |
| COG0091 | Ribosomal protein L22 | COG0292 | Ribosomal protein L20 | COG0190 | 5,10-methylene-tetrahydrofolate dehydrogenase/Methenyl tetrahydrofolate cyclohydrolase |
| COG0092 | Ribosomal protein S3 | COG0359 | Ribosomal protein L9 | COG0195 | Transcription antitermination factor NusA, contains S1 and KH domains |
| COG0093 | Ribosomal protein L14 | COG0504 | CTP synthase (UTP-ammonia lyase) | COG0196 | FAD synthase |
| COG0094 | Ribosomal protein L5 | COG0536 | GTPase involved in cell partioning and DNA repair | COG0215 | Cysteinyl-tRNA synthetase |
| COG0096 | Ribosomal protein S8 | COG0544 | FKBP-type peptidyl-prolyl cis-trans isomerase (trigger factor) | COG0217 | Transcriptional and/or translational regulatory protein YebC/TACO1 |
| COG0097 | Ribosomal protein L6P/L9E | COG0592 | DNA polymerase III sliding clamp (beta) subunit, PCNA homolog | COG0249 | DNA mismatch repair ATPase MutS |
| COG0098 | Ribosomal protein S5 | COG0750 | Membrane-associated protease RseP, regulator of RpoE activity | COG0250 | Transcription antitermination factor NusG |
| COG0103 | Ribosomal protein S9 | COG1185 | Polyribonucleotide nucleotidyltransferase (polynucleotide phosphorylase) | COG0272 | NAD-dependent DNA ligase |
| COG0126 | 3-phosphoglycerate kinase | COG1198 | Primosomal protein N' (replication factor Y) - superfamily II helicase | COG0283 | Cytidylate kinase |
| COG0127 | Inosine/xanthosine triphosphate pyrophosphatase, all-alpha NTP-PPase family | COG1530 | Ribonuclease G or E | COG0289 | Dihydrodipicolinate reductase |
| COG0197 | Ribosomal protein L16/L10AE | | | COG0322 | Excinuclease UvrABC, nuclease subunit |
| COG0200 | Ribosomal protein L15 | | | COG0323 | DNA mismatch repair ATPase MutL |
| COG0201 | Preprotein translocase subunit SecY | | | COG0324 | tRNA A37 N6-isopentenylltransferase MiaA |
| COG0203 | Ribosomal protein L17 | | | COG0335 | Ribosomal protein L19 |
| COG0216 | Protein chain release factor A | | | COG0445 | tRNA U34 5-carboxymethylaminomethyl modifying enzyme MnmG/GidA |
| COG0233 | Ribosome recycling factor | | | COG0449 | Glucosamine 6-phosphate synthetase, contains amidotransferase and phosphosugar isomerase domains |
| COG0244 | Ribosomal protein L10 | | | COG0481 | Translation elongation factor EF-4, membrane-bound GTPase |
| COG0256 | Ribosomal protein L18 | | | COG0511 | Biotin carboxyl carrier protein |
| COG0261 | Ribosomal protein L21 | | | COG0522 | Ribosomal protein S4 or related protein |
| COG0264 | Translation elongation factor EF-Ts | | | COG0525 | Valyl-tRNA synthetase |
| COG0290 | Translation initiation factor IF-3 | | | COG0532 | Translation initiation factor IF-2, a GTPase |
| COG0331 | Malonyl CoA-acyl carrier protein transacylase | | | COG0541 | Signal recognition particle GTPase |
| COG0353 | Recombinational DNA repair protein RecR | | | COG0552 | Signal recognition particle GTPase |
| COG0468 | RecA/RadA recombinase | | | COG0575 | CDP-diglyceride synthetase |
| COG0495 | Leucyl-tRNA synthetase | | | COG0749 | DNA polymerase I - 3'-5' exonuclease and polymerase domains |
| COG0576 | Molecular chaperone GrpE (heat shock protein) | | | COG0774 | UDP-3-O-acyl-N-acetylglucosamine deacetylase |
| COG0632 | Holliday junction resolvasome RuvABC DNA-binding subunit | | | COG0825 | Acetyl-CoA carboxylase alpha subunit |
| COG0769 | UDP-N-acetylmuramyl tripeptide synthase | | | COG0858 | Ribosome-binding factor A |
| COG0815 | Apolipoprotein N-acyltransferase | | | COG1044 | UDP-3-O-[3-hydroxymyristoyl] glucosamine N-acyltransferase |
| COG0817 | Holliday junction resolvasome RuvABC endonuclease subunit | | | COG1137 | ABC-type lipopolysaccharide export system, ATPase component |
| COG2255 | Holliday junction resolvasome RuvABC, ATP-dependent DNA helicase subunit | | | COG1160 | Predicted GTPases |
| | | | | COG1570 | Exonuclease VII, large subunit |
| | | | | COG1663 | Tetraacyldisaccharide-1-P 4'-kinase |
| | | | | COG1825 | Ribosomal protein L25 (general stress protein Ctc) |
| | | | | COG2877 | 3-deoxy-D-manno-octulosonic acid (KDO) 8-phosphate synthase |
| | | | | COG2890 | Methylase of polypeptide chain release factors |

#### Supplementary Data Descriptions

**Supplementary Data 1.** Chlamydiae 16S rRNA amplicon OTUs in FASTA format

**Supplementary Data 2.** Relative abundance of Chlamydiae OTUs across Loki's Castle marine sediments.

**Supplementary Data 3.** Pathway overviews, selected gene annotations and raw data.

**Tab 1.** Presence and absence of bacterial level NOGs across Chlamydiae

**Tab 2.** EffectiveDB results

**Tab 3.** Overview of KEGG pathways and their presence across Chlamydiae including central carbon metabolism, carbon fixation, amino acid and nucleotide biosynthesis

**Tab 4.** Secretion systems and flagellar components identified by MacSyFinder

**Tab 5.** Selected gene annotations

**Tab 6.** IMNGS results

**Supplementary Data 4.** Unprocessed phylogenetic trees presented in this study.
