## Supplementary Data 4 for "Marine sediments illuminate Chlamydiae diversity and evolution"

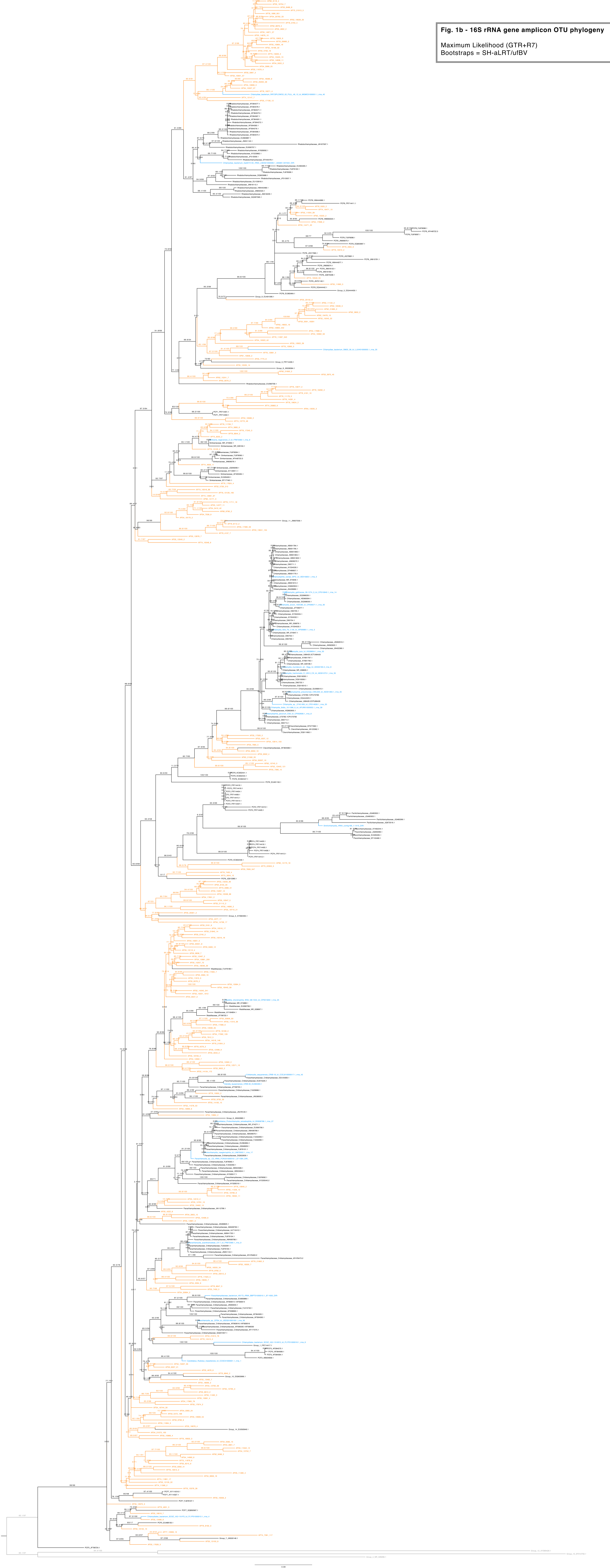

**Fig. 1b - 16S rRNA gene amplicon OTU phylogeny**

Maximum Likelihood (GTR+R7)
Bootstraps = SH-aLRT/ufBV

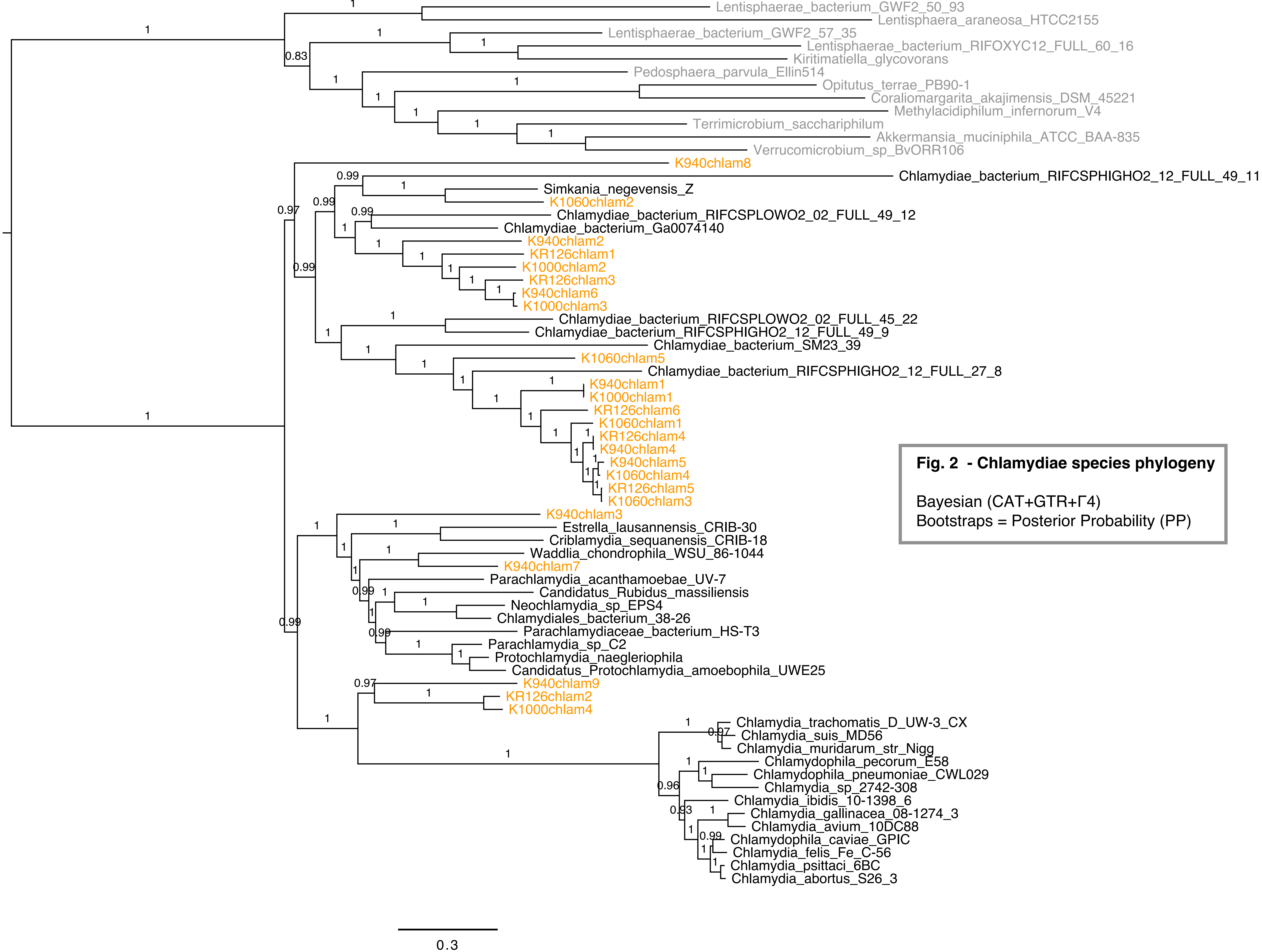


**Fig. 2 - Chlamydiae species phylogeny**

Bayesian (CAT+GTR+Γ4)
Bootstraps = Posterior Probability (PP)

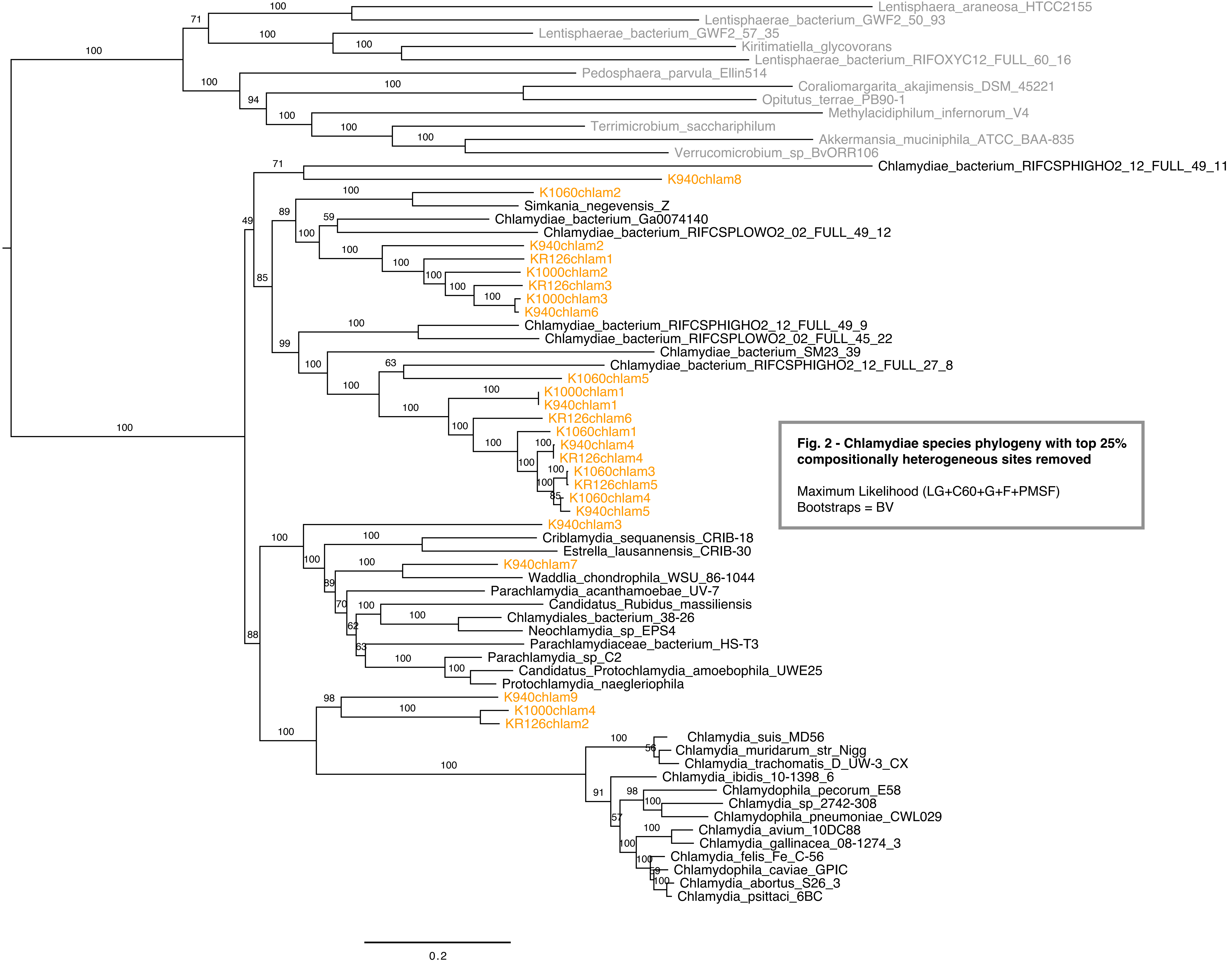


**Fig. 2 - Chlamydiae species phylogeny with top 25% compositionally heterogeneous sites removed**

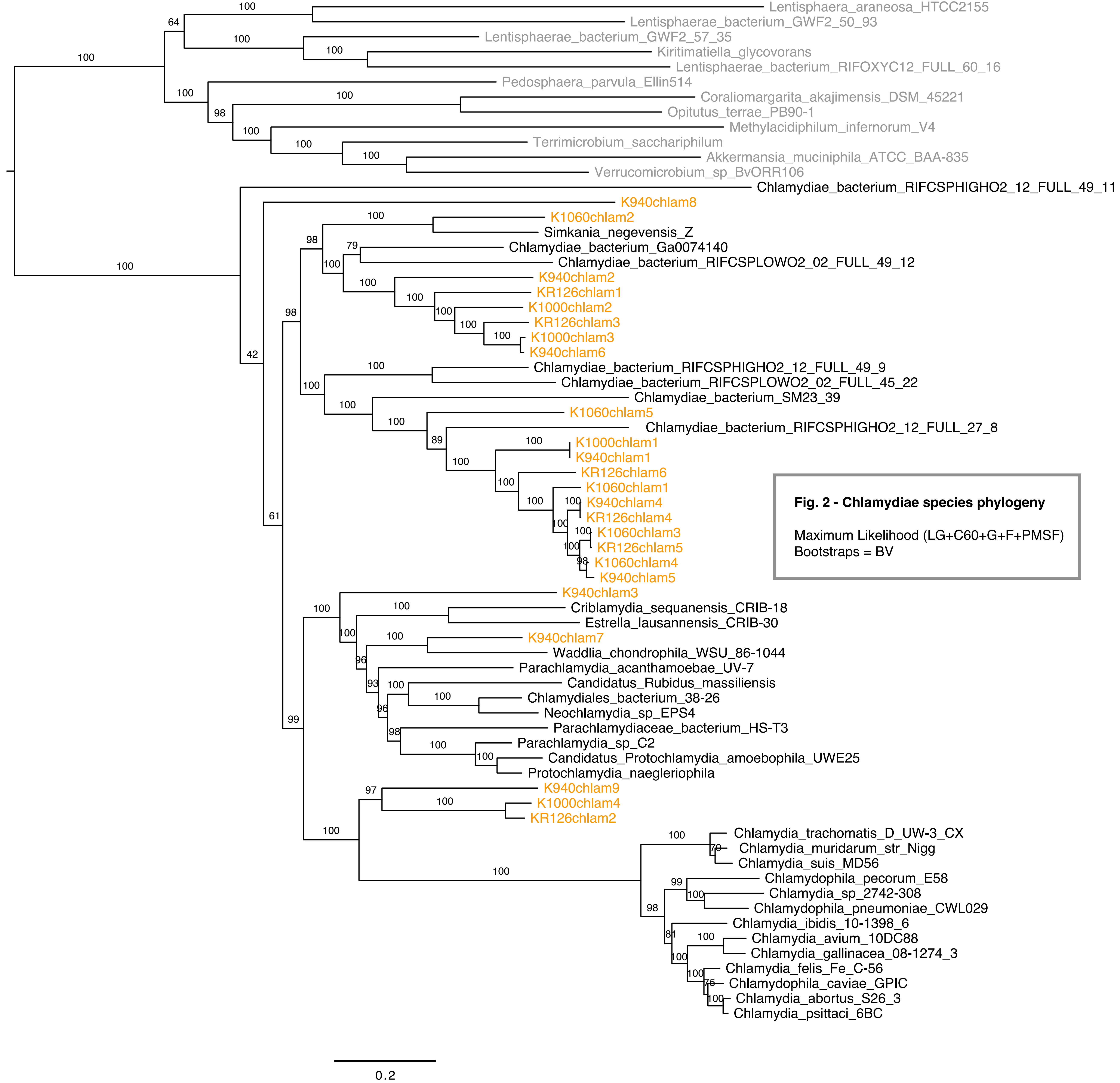


**Fig. 2 - Chlamydiae species phylogeny**

Maximum Likelihood (LG+C60+G+F+PMSF)
Bootstraps = BV

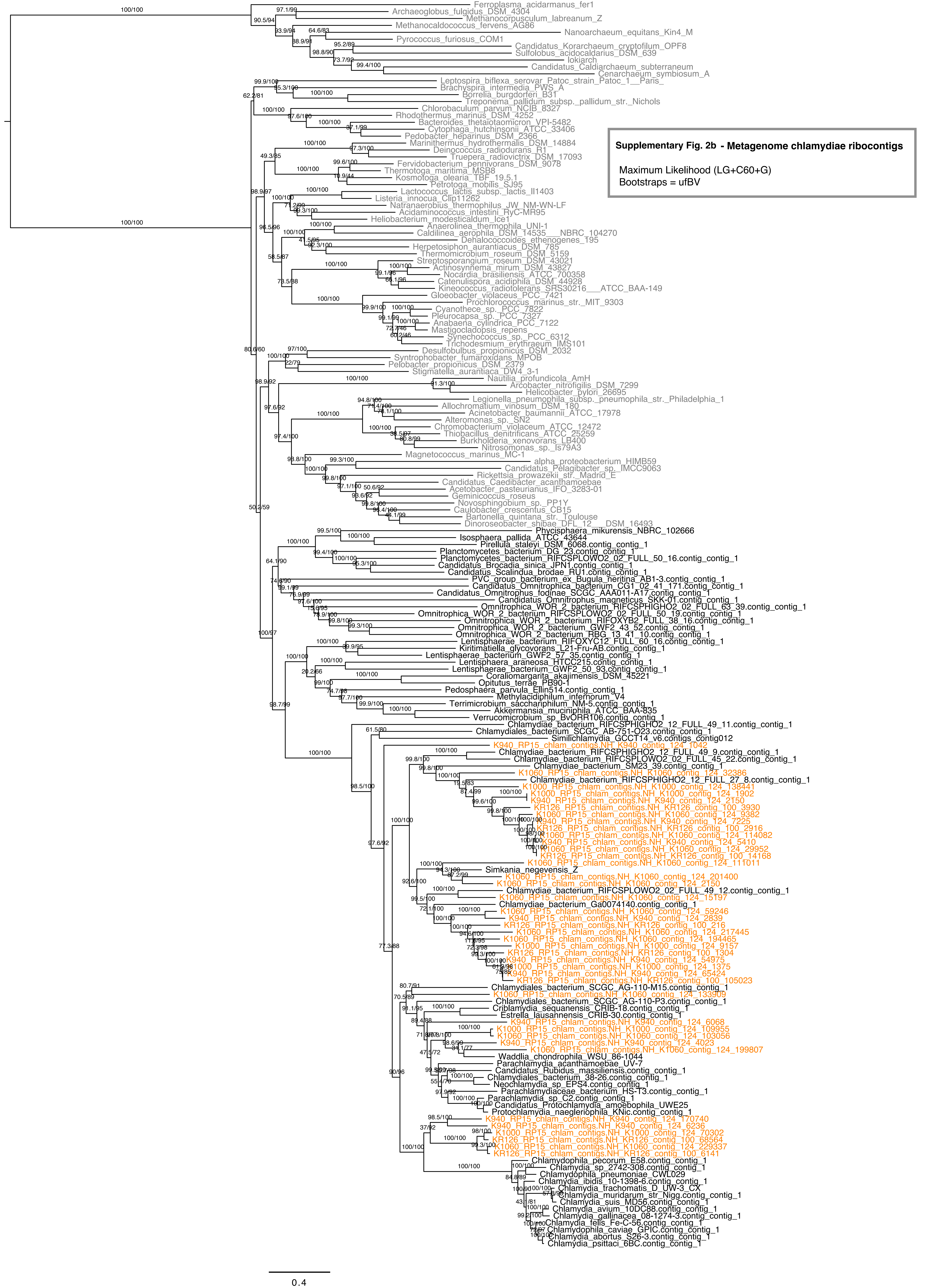


**Supplementary Fig. 2b - Metagenome chlamydiae ribocontigs**

Maximum Likelihood (LG+C60+G)
Bootstraps = ufBV

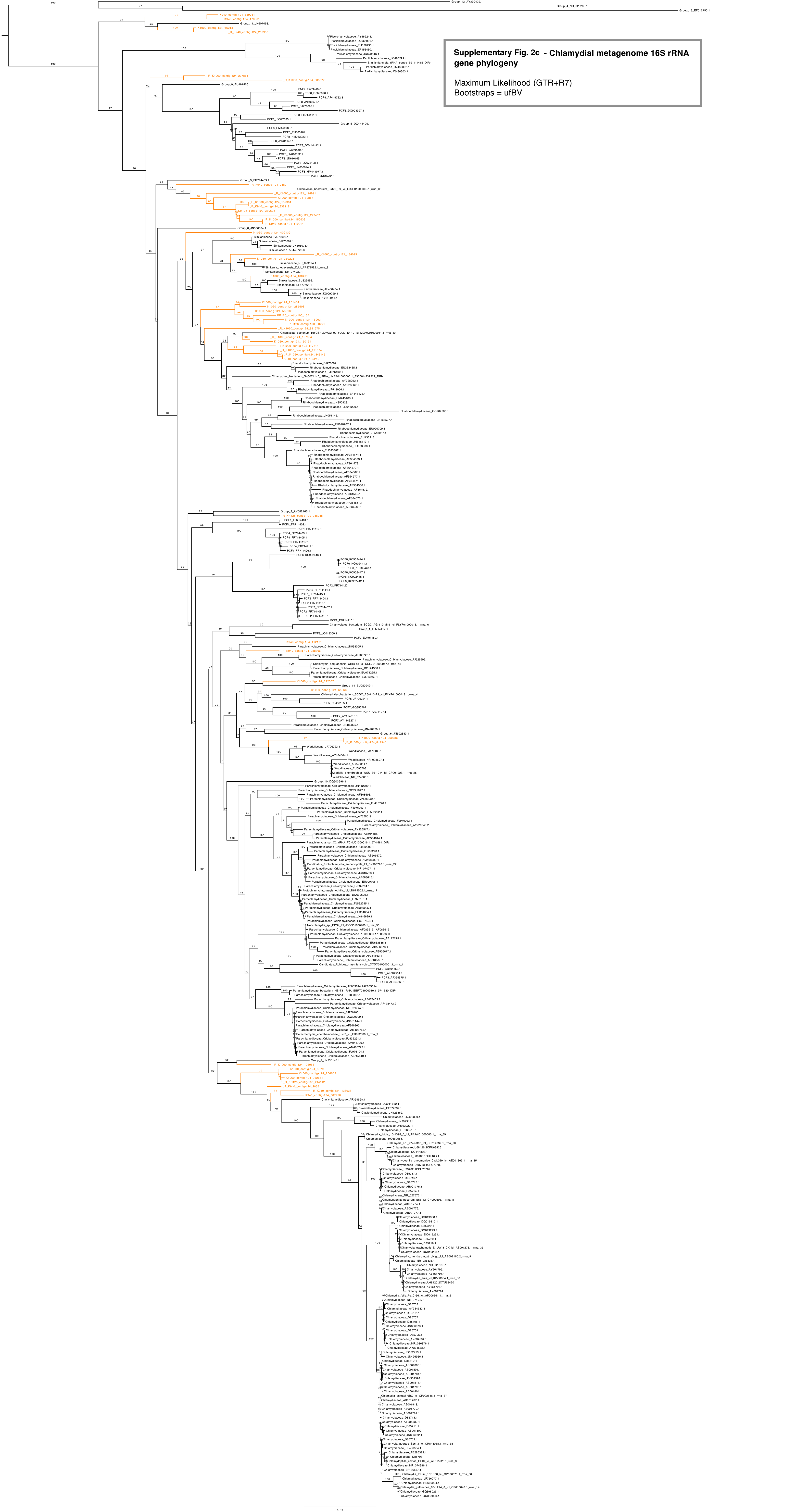

**Supplementary Fig. 2c - Chlamydial metagenome 16S rRNA gene phylogeny**

Maximum Likelihood (GTR+R7)
Bootstraps = ufBV

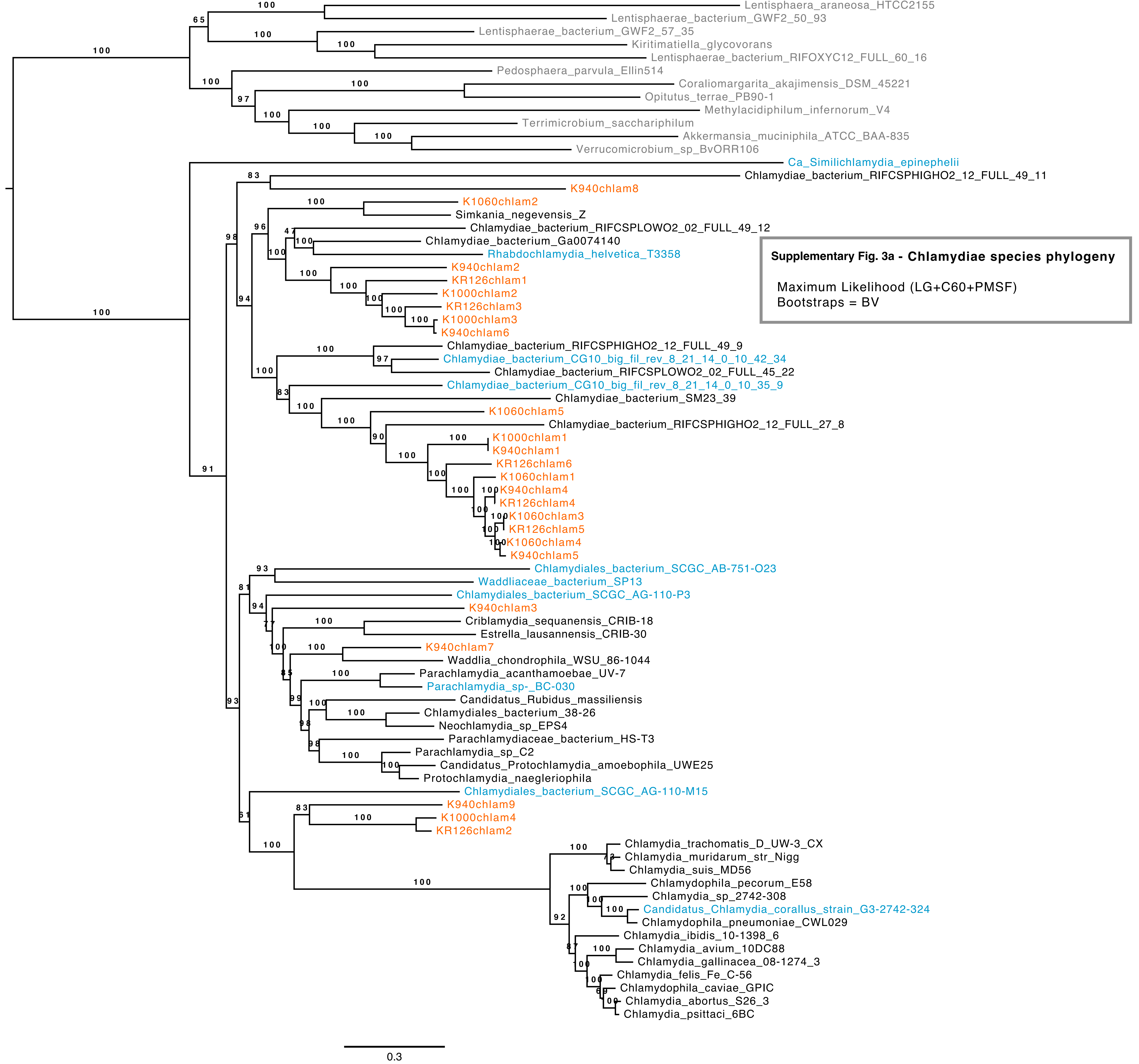


**Supplementary Fig. 3a - Chlamydiae species phylogeny**

Maximum Likelihood (LG+C60+PMSF)
Bootstraps = BV

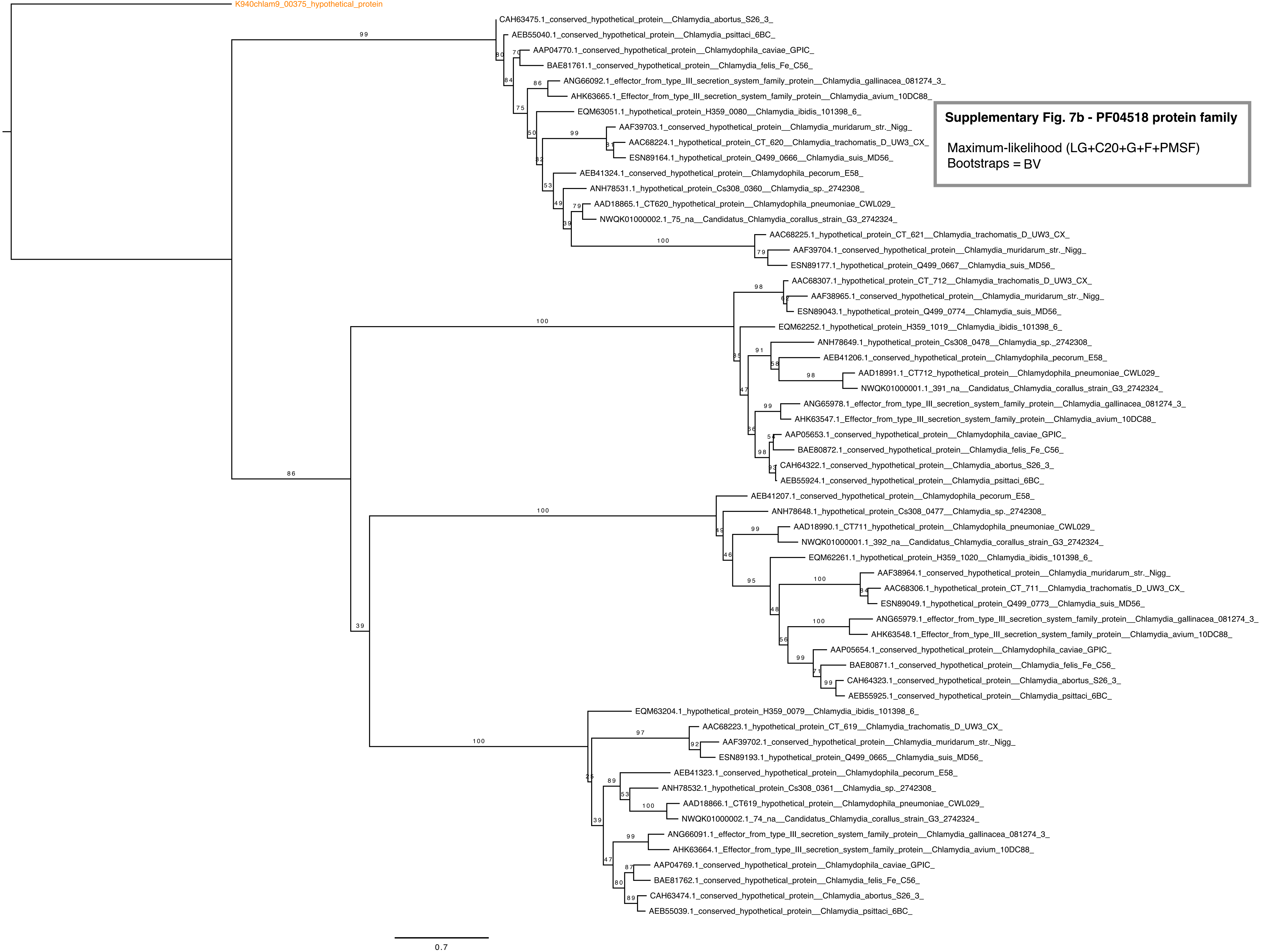


**Supplementary Fig. 7b - PF04518 protein family**

Maximum-likelihood (LG+C20+G+F+PMSF)
Bootstraps = BV

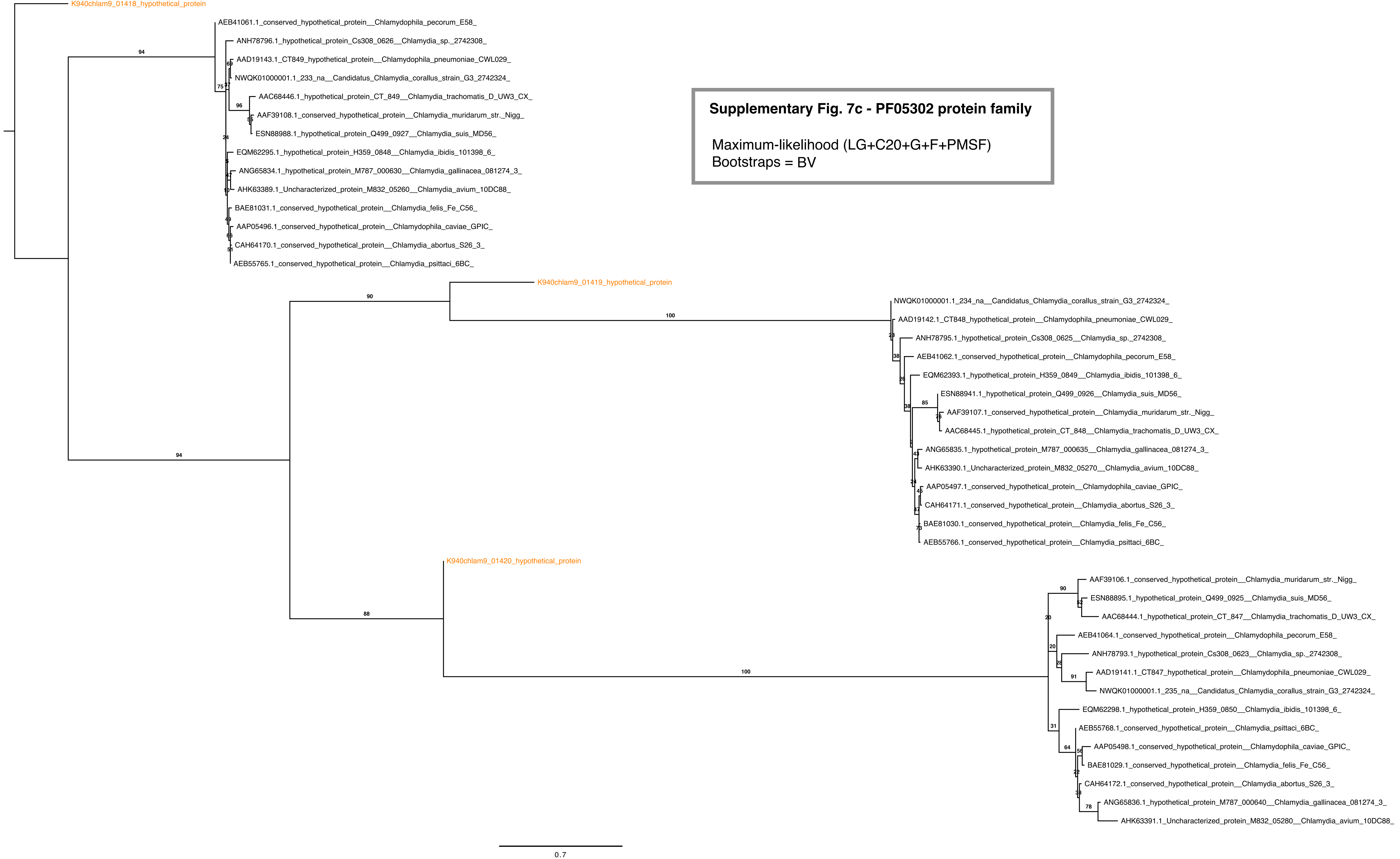


**Supplementary Fig. 7c - PF05302 protein family**

Maximum-likelihood (LG+C20+G+F+PMSF)
Bootstraps = BV

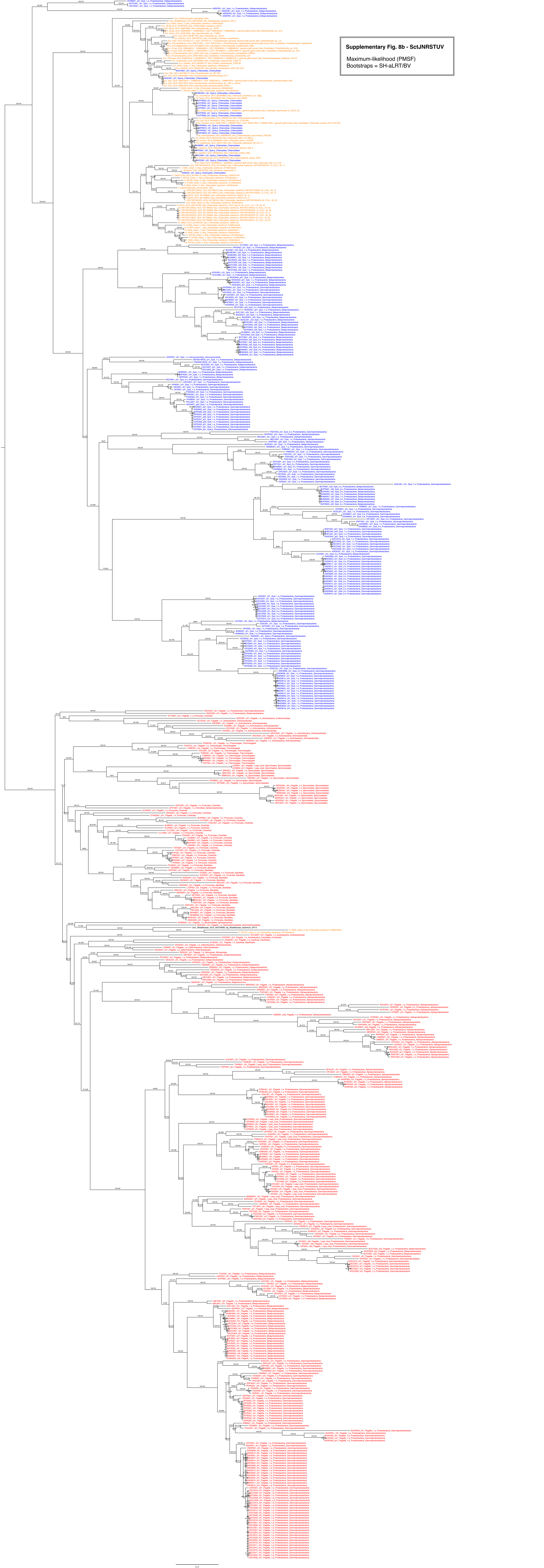
Supplementary Fig. 8b - SctJNRSTUV
Maximum-likelihood (PMSF)
Bootstraps = SH-aLRT/BV
0.3

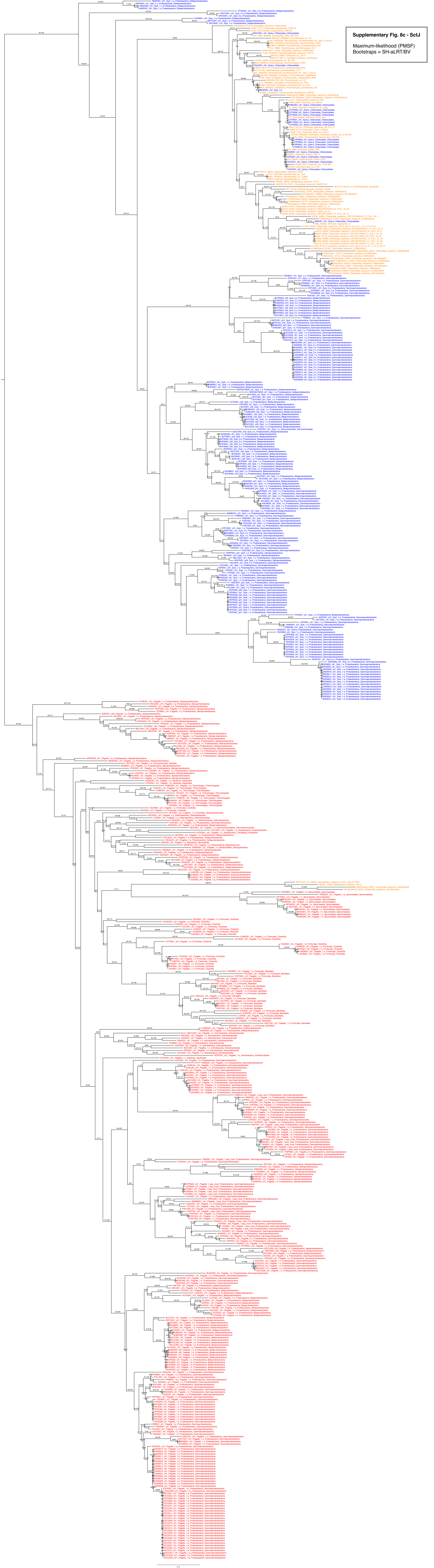
Supplementary Fig. 8c - SctJ
Maximum-likelihood (PMSF)
Bootstraps = SH-aLRT/BV

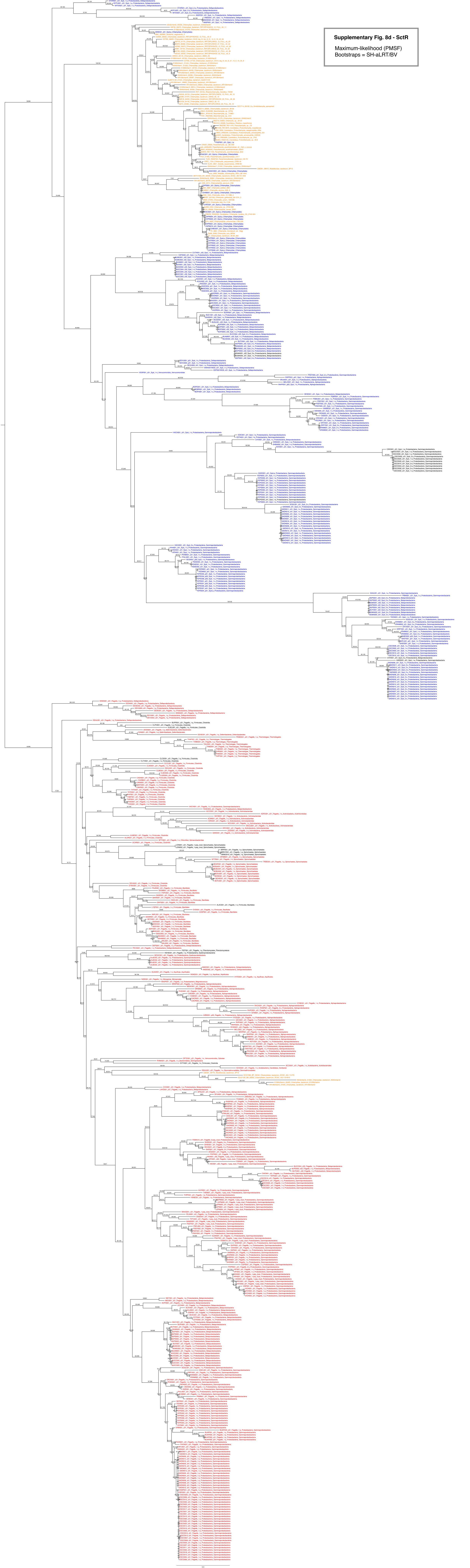

Supplementary Fig. 8d - SctR
Maximum-likelihood (PMSF)
Bootstraps = SH-aLRT/BV

**Supplementary Fig. 12a - "Canonical NTTs"**

Maximum Likelihood (LG+F+R8)
Bootstraps = SH-aLRT/ufBV

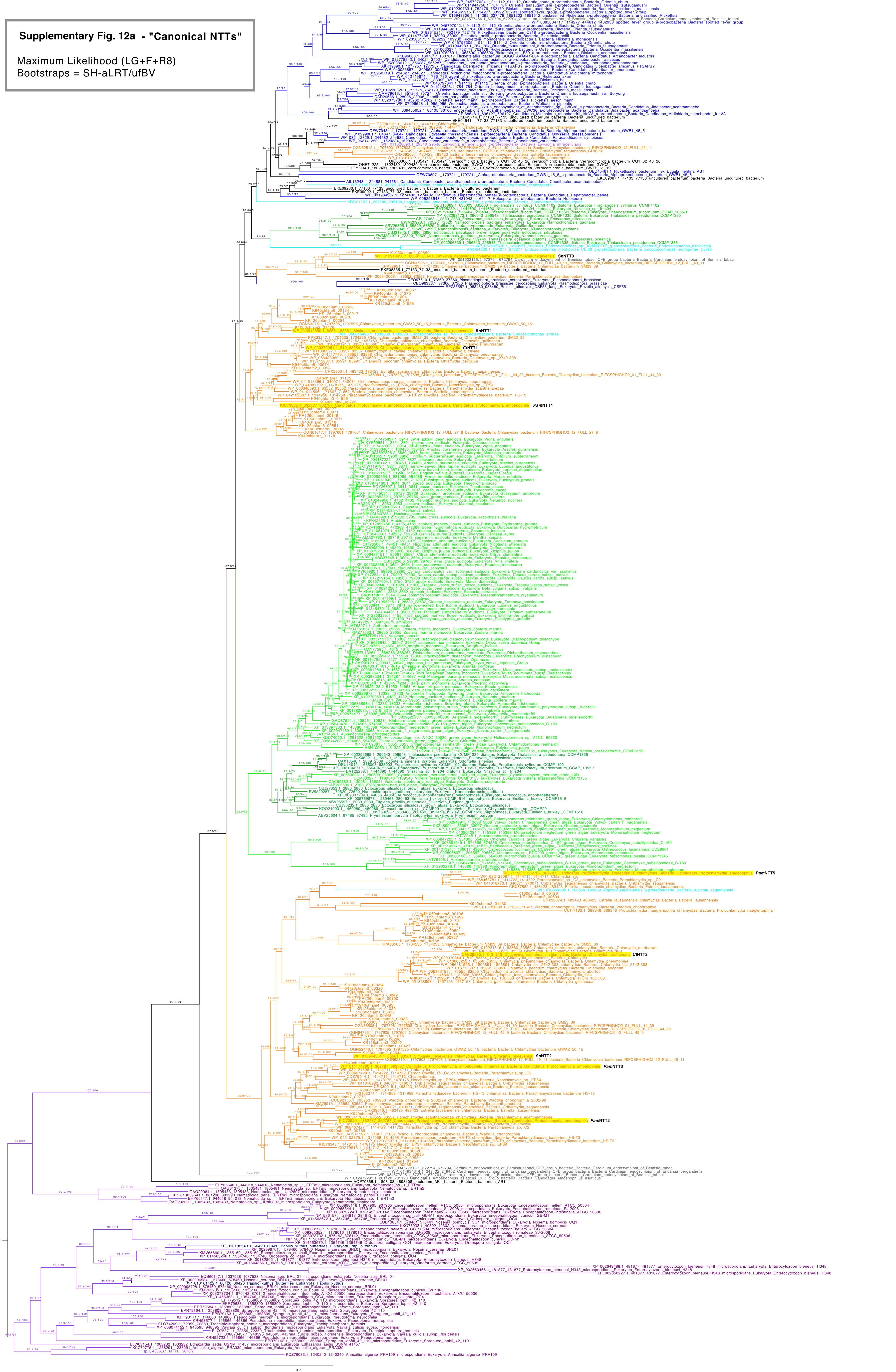

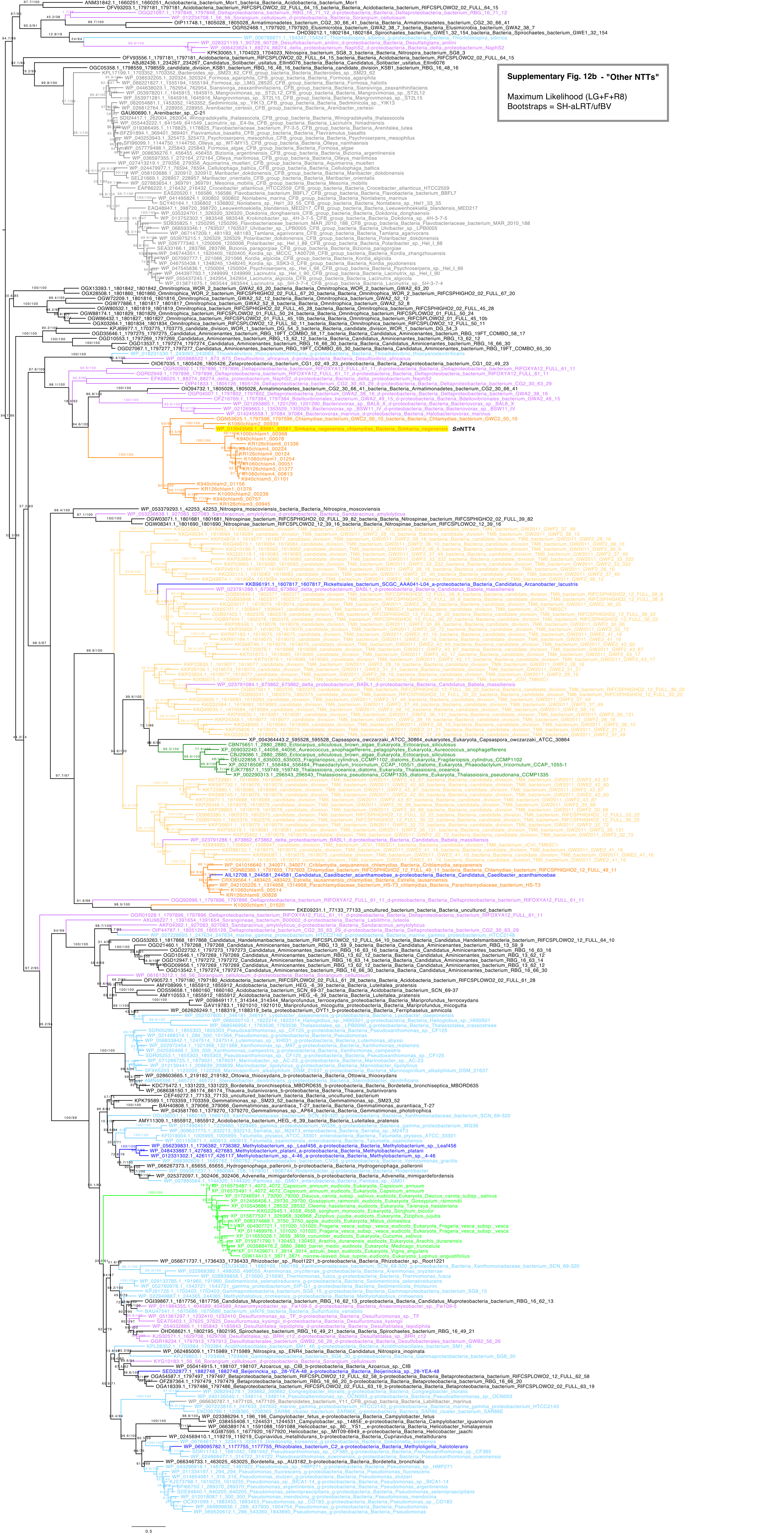

**Supplementary Fig. 12b - "Other NTTs"**

Maximum Likelihood (LG+F+R8)
Bootstraps = SH-aLRT/ufBV

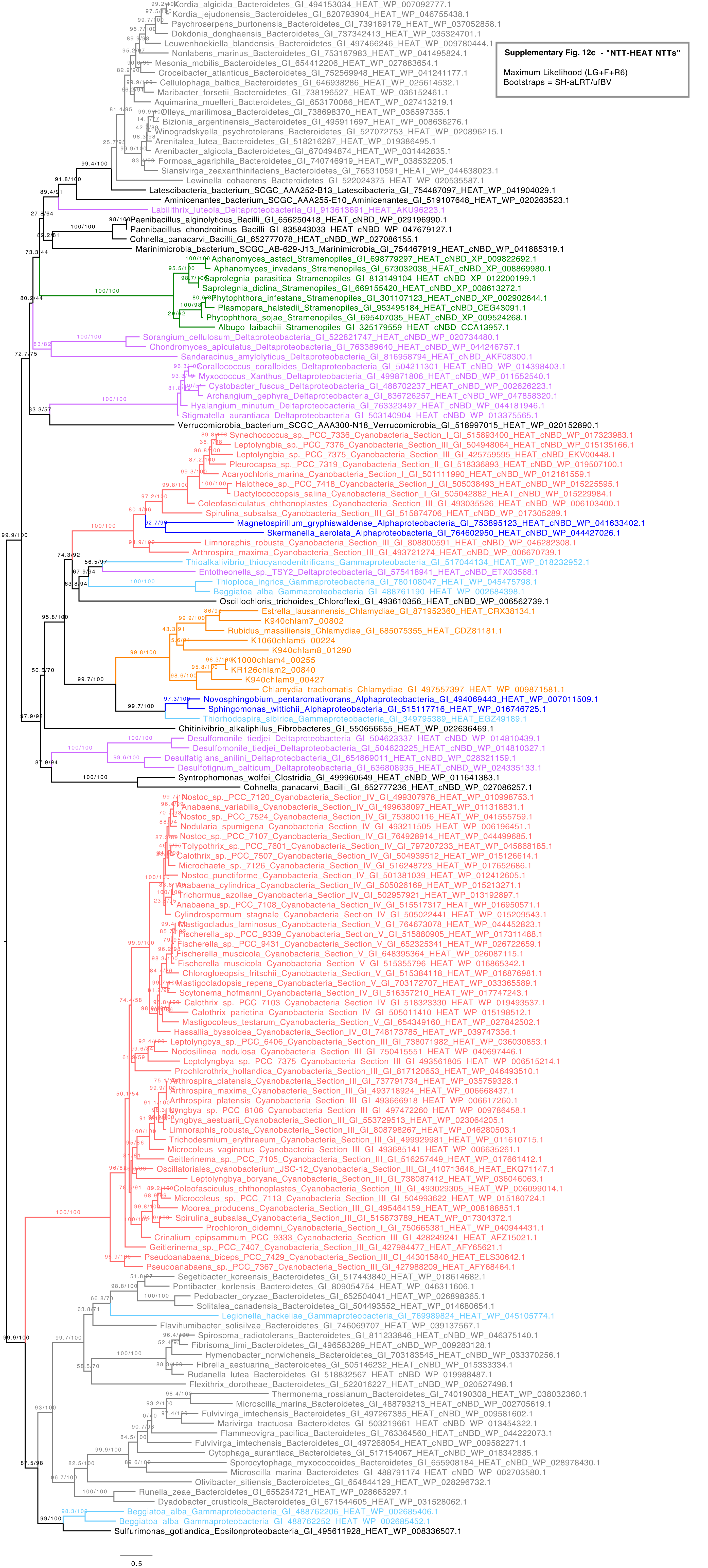


**Supplementary Fig. 12c - "NTT-HEAT NTTs"**

Maximum Likelihood (LG+F+R6)
Bootstraps = SH-aLRT/ufBV

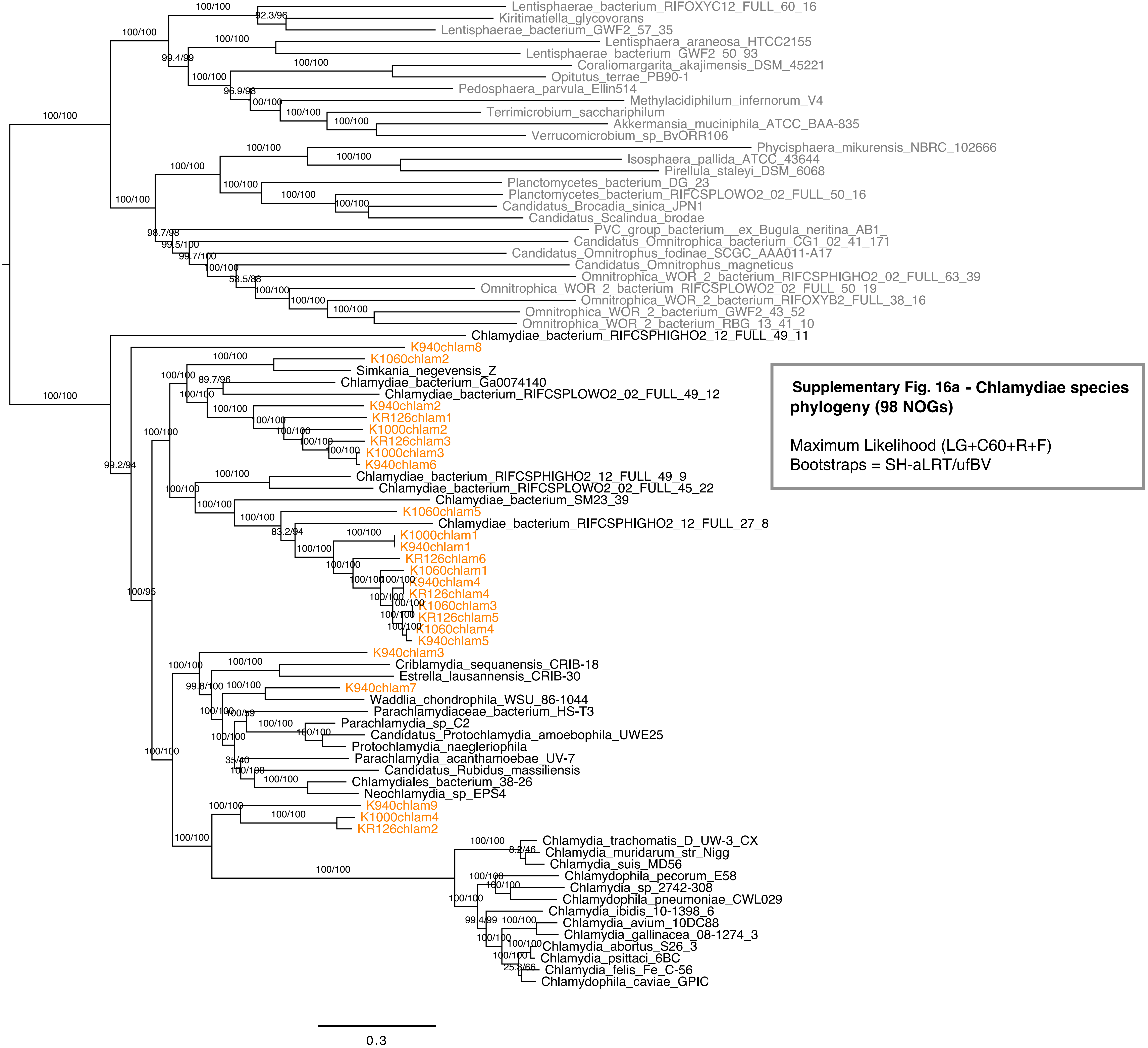


**Supplementary Fig. 16a - Chlamydiae species phylogeny (98 NOGs)**

Maximum Likelihood (LG+C60+R+F)
Bootstraps = SH-aLRT/ufBV

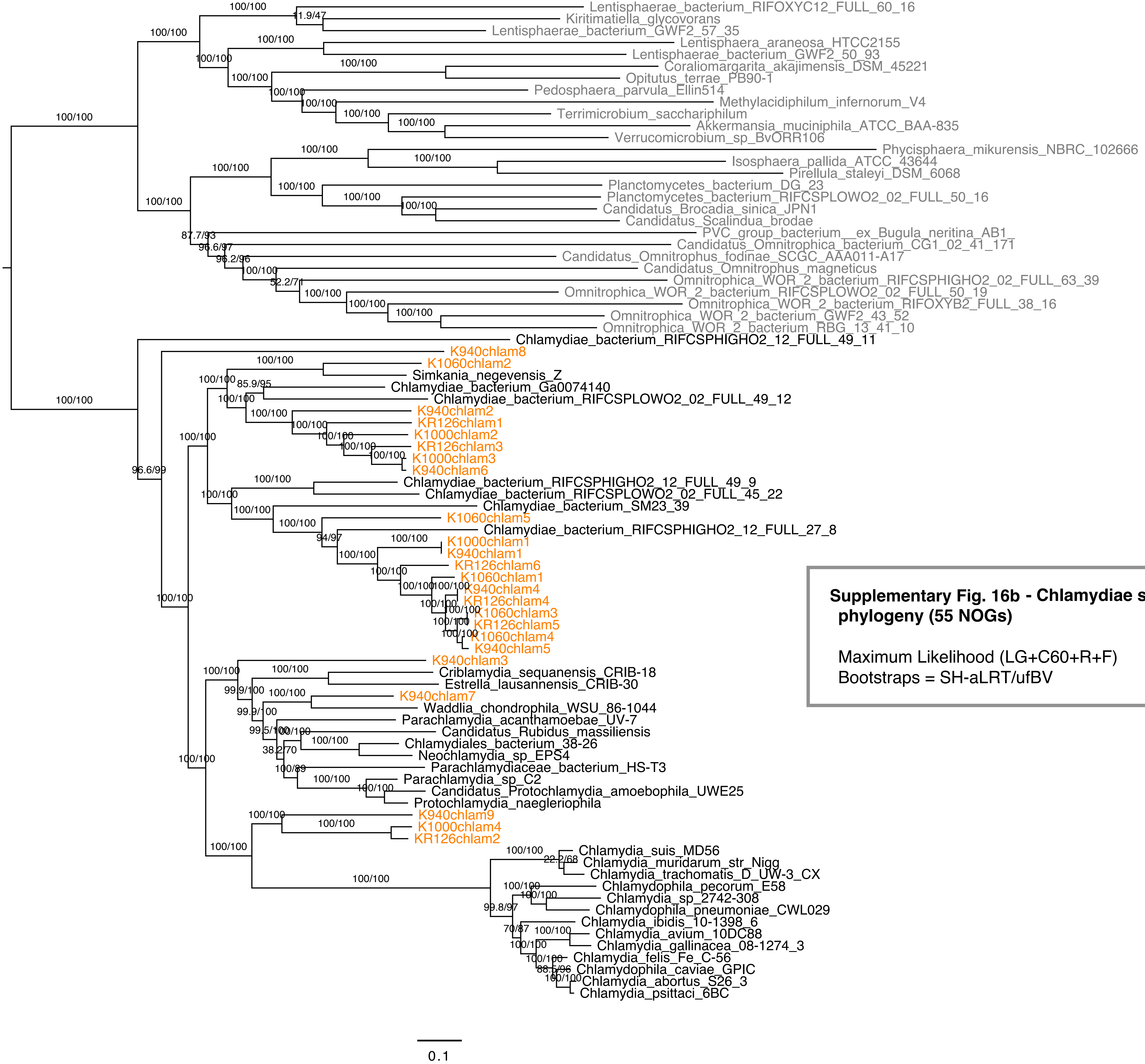


**Supplementary Fig. 16b - Chlamydiae species phylogeny (55 NOGs)**

Maximum Likelihood (LG+C60+R+F)
Bootstraps = SH-aLRT/ufBV

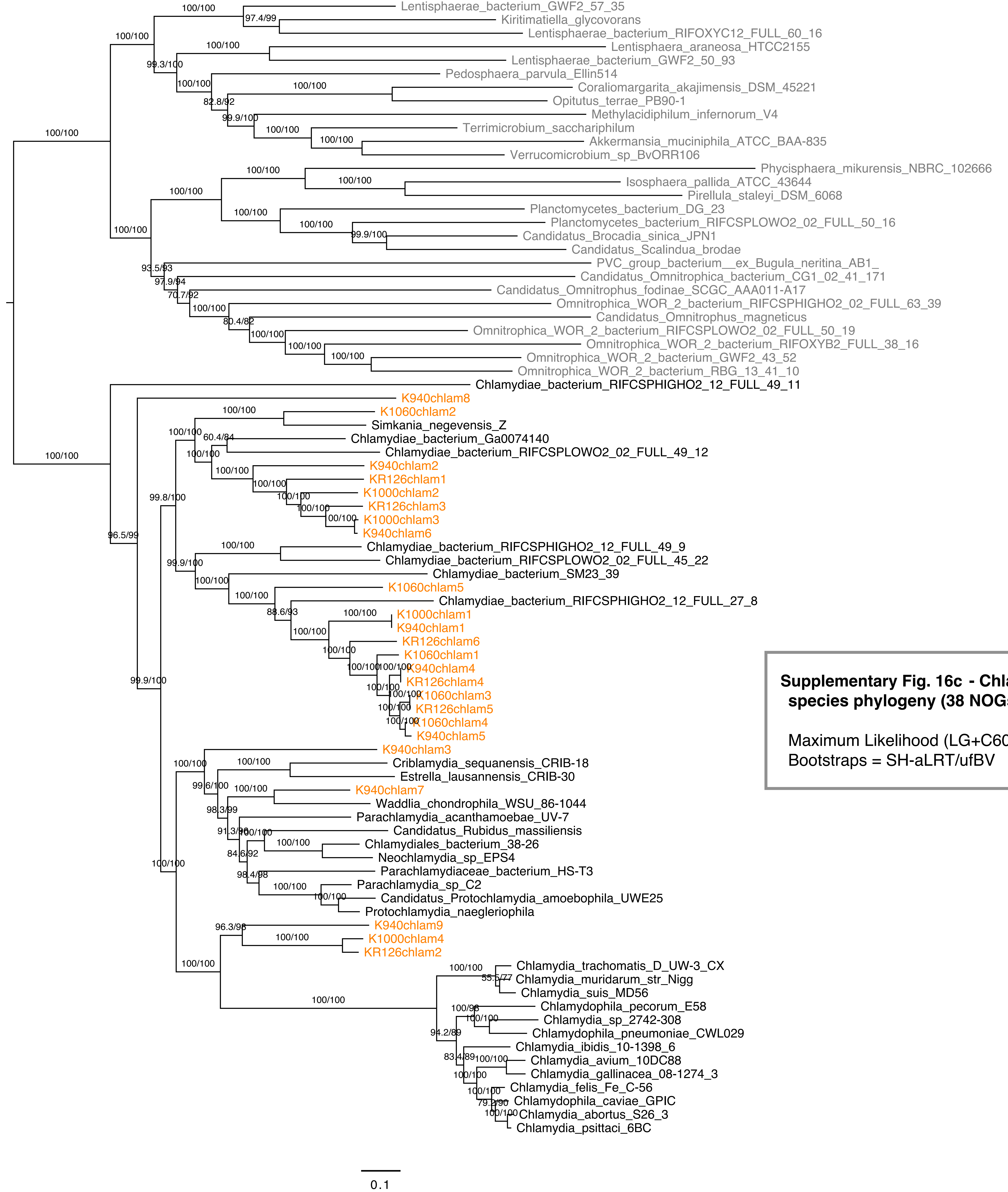


**Supplementary Fig. 16c - Chlamydiae species phylogeny (38 NOGs)**

Maximum Likelihood (LG+C60+R+F)
Bootstraps = SH-aLRT/ufBV

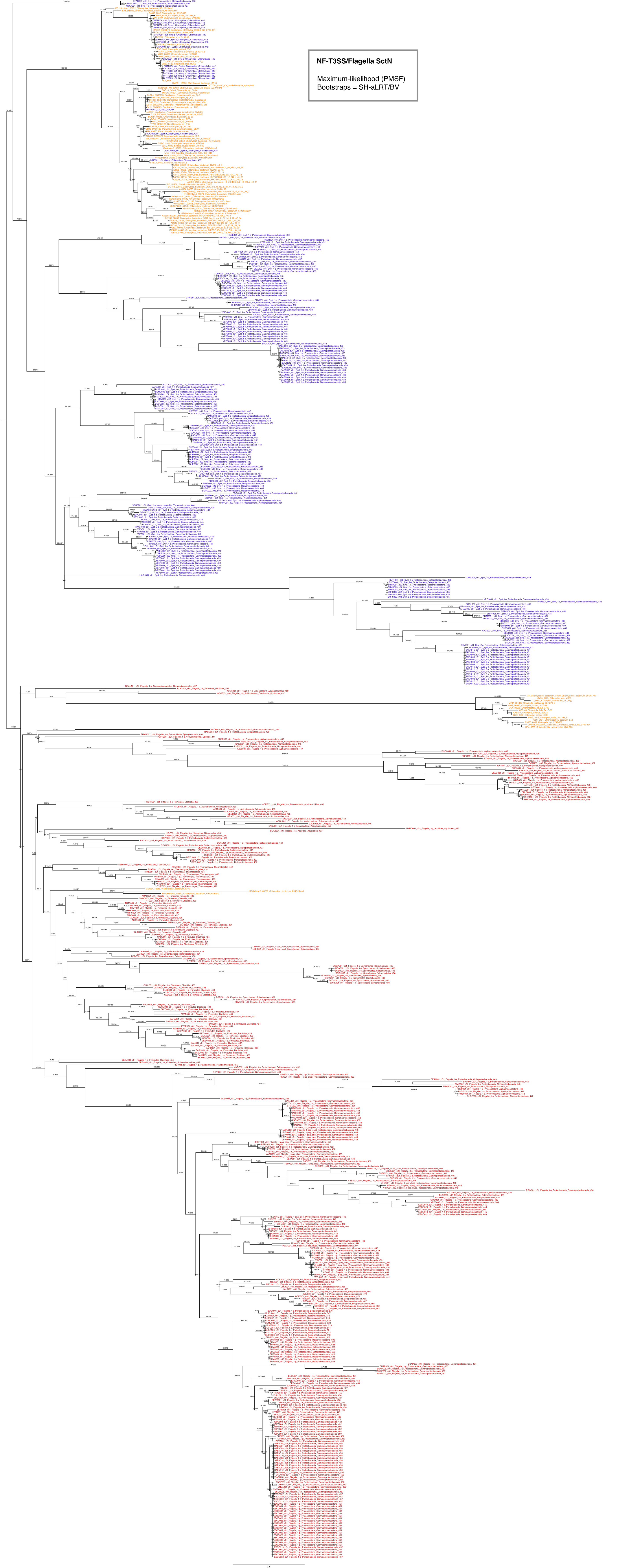
NF-T3SS/Flagella SctN
Maximum-likelihood (PMSF)
Bootstraps = SH-aLRT/BV

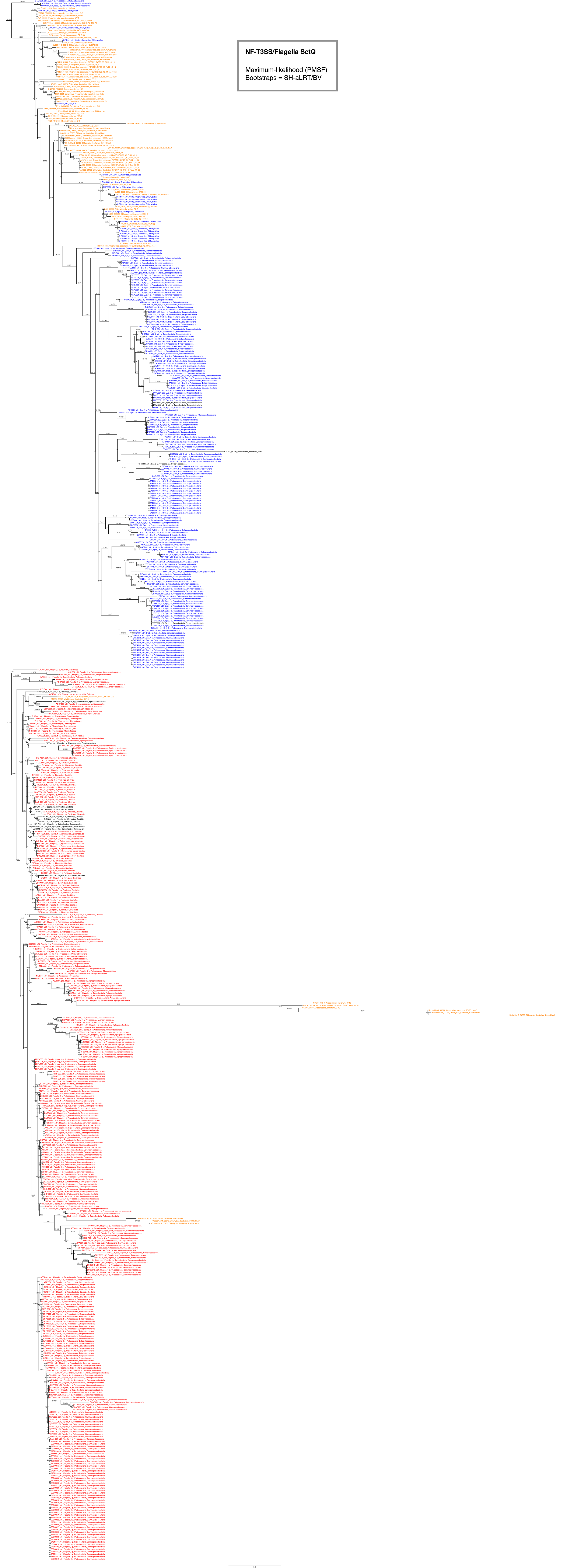
NF-T3SS/Flagella SctQ
Maximum-likelihood (PMSF)
Bootstraps = SH-aLRT/BV

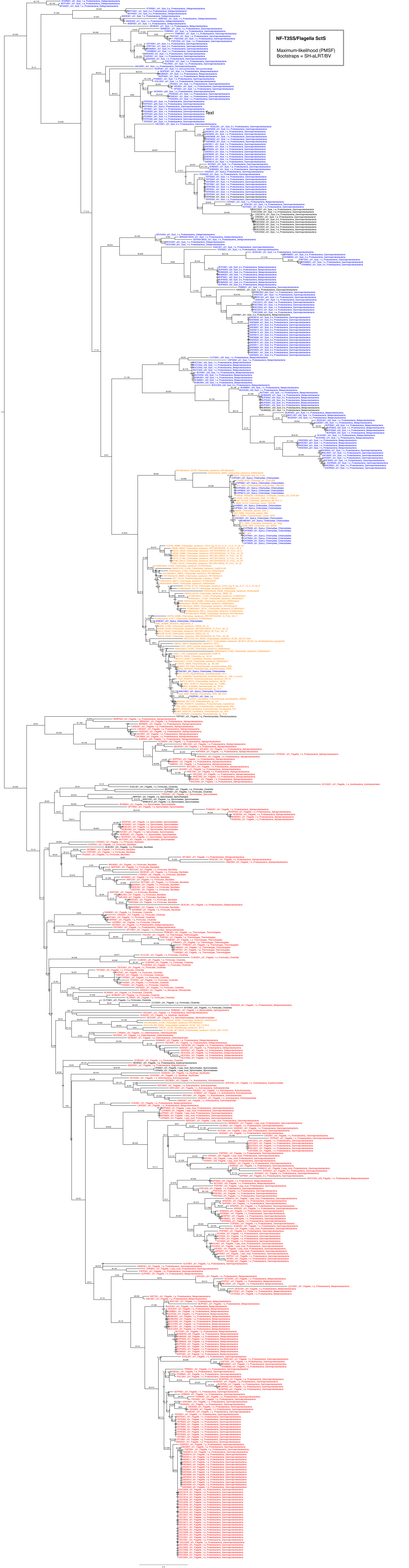
NF-T3SS/Flagella SctS
Maximum-likelihood (PMSF)
Bootstraps = SH-aLRT/BV
Text

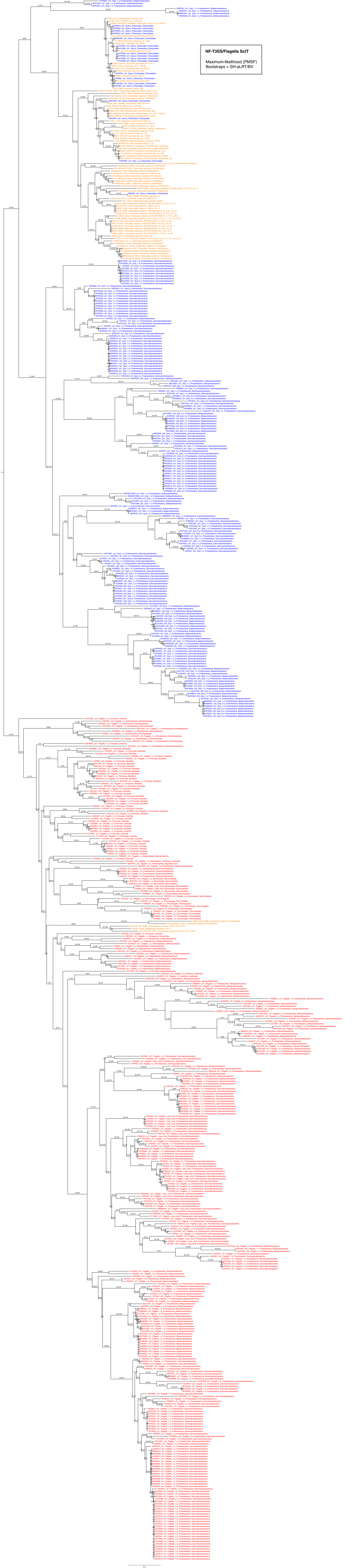
NF-T3SS/Flagella SctT
Maximum-likelihood (PMSF)
Bootstraps = SH-aLRT/BV

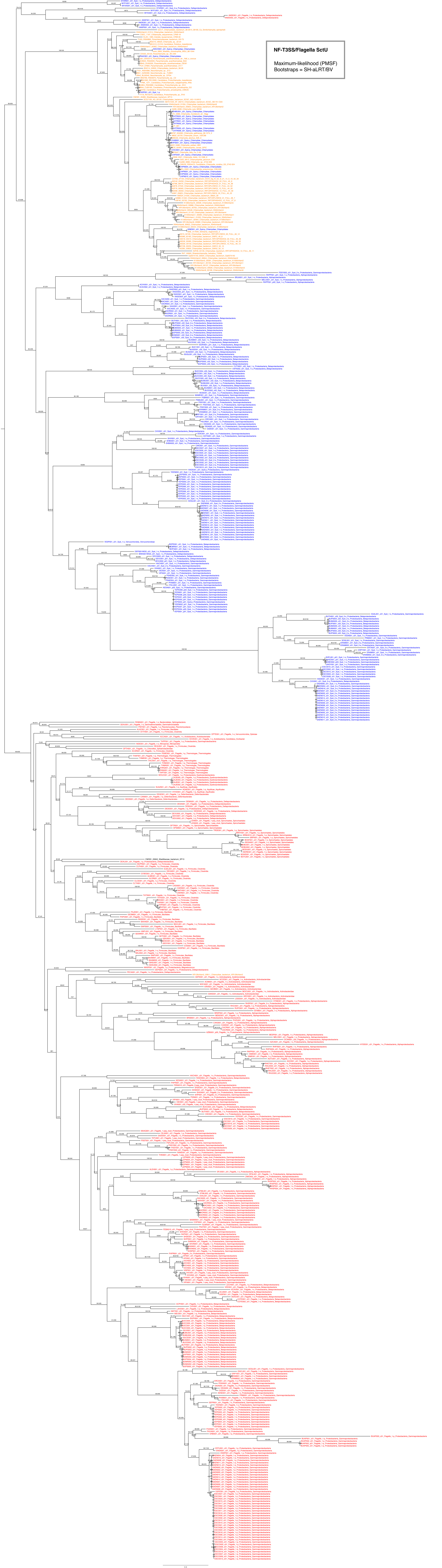
NF-T3SS/Flagella SctU
Maximum-likelihood (PMSF)
Bootstraps = SH-aLRT/BV

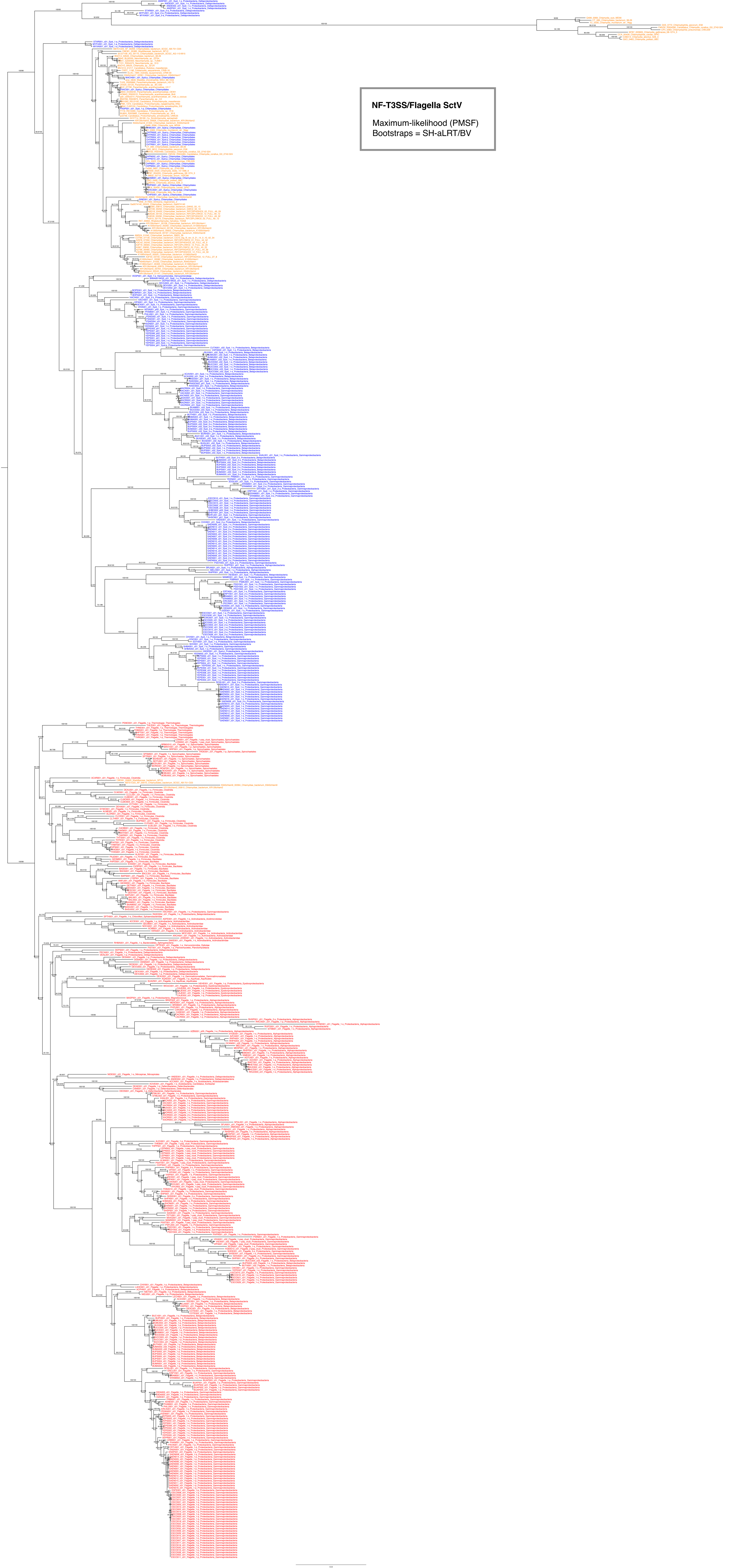
NF-T3SS/Flagella SctV
Maximum-likelihood (PMSF)
Bootstraps = SH-aLRT/BV
